# The differential impacts of dataset imbalance in single-cell data integration

**DOI:** 10.1101/2022.10.06.511156

**Authors:** Hassaan Maan, Lin Zhang, Chengxin Yu, Michael Geuenich, Kieran R Campbell, Bo Wang

## Abstract

Single-cell transcriptomic data measured across distinct samples has led to a surge in computational methods for data integration. Few studies have explicitly examined the common case of cell-type imbalance between datasets to be integrated, and none have characterized its impact on downstream analyses. To address this gap, we developed the *Iniquitate* pipeline for assessing the stability of single-cell RNA sequencing (scRNA-seq) integration results after perturbing the degree of imbalance between datasets. Through benchmarking 5 state-of-the-art scRNA-seq integration techniques in 1600 perturbed integration scenarios for a multi-sample peripheral blood mononuclear cell (PBMC) dataset, our results indicate that sample imbalance has significant impacts on downstream analyses and the biological interpretation of integration results. We observed significant variation in clustering, cell-type classification, marker gene-based annotation, and query-to-reference mapping in imbalanced settings. Two key factors were found to lead to quantitation differences after scRNA-seq integration - the cell-type imbalance within and between samples (*relative cell-type support*) and the relatedness of cell-types across samples (*minimum cell-type center distance*). To account for evaluation gaps in imbalanced contexts, we developed novel clustering metrics robust to sample imbalance, including the balanced Adjusted Rand Index (bARI) and balanced Adjusted Mutual Information (bAMI). Our analysis quantifies biologically-relevant effects of dataset imbalance in integration scenarios and introduces guidelines and novel metrics for integration of disparate datasets. The Iniquitate pipeline and balanced clustering metrics are available at https://github.com/hsmaan/Iniquitate and https://github.com/hsmaan/balanced-clustering, respectively.

## Introduction

Single-cell sequencing technologies developed in the past decade have led to breakthrough discoveries due to the high resolution that they offer in determining biological heterogeneity [1–3]. A major challenge associated with the analysis of high throughput sequencing data is that of accounting for batch effects, which are technical artifacts caused by factors such as differences in sequencing protocols, experimental reagents, and ambient conditions that lead to quantification changes that are not biologically driven [4]. Batch effects can lead to major discrepancies in comparisons of similar experimental groups that can easily be misinterpreted as biological signal [5]. The amount of mRNA captured and reads sequenced per cell in single-cell RNA sequencing (scRNA-seq) assays is very low compared to their bulk counterparts, leading to measurements that tend to be sparse and noisy [6, 7]. These factors, combined with measurements often conducted across separate experimental groups without balanced designs [7], leads to a higher susceptibility of scRNA-seq data to batch effects. Methods for removing batch effects from bulk RNA sequencing data have demonstrated poor performance in single-cell settings due to invalid assumptions of shared populations and linear application of technical effects [8]. To account for this gap in methodology, batch correction/integration techniques have been developed specifically for scRNA-seq data [8].

Current single-cell integration methods underperform in settings where datasets are imbalanced based on cell-types [9]. More specifically, this form of imbalance is dictated by differences in the cell-types present, number of cells per cell-type, and cell-type proportions across samples [9, 10]. Imbalanced datasets occur in many integration contexts, including developmental and cancer biology. In developmental data, it is unlikely that cell populations and proportions will be shared across samples from different developmental time-points due to factors such as depletion of stem-like progenitors and differentiation [11]. In tumor samples, both clonal and subclonal heterogeneity can be present, as well as different levels of immune and stromal cell infiltration, both within and across samples [12]. Therefore, as imbalanced contexts can be common in single-cell data analysis, integration methods and analysis pipelines must be able to explicitly address these imbalances or integration results may lead to inaccurate biological conclusions.

In comprehensive single-cell integration benchmarking studies by Tran et al. and Luecken et al. [9, 13], scRNA-seq integration methods were found to perform poorly in terms of both batch-correction and cell-type identity conservation metrics, particularly in large and imbalanced datasets. Ming et al. [10] highlighted dataset imbalance limitations through simulation studies for balanced and imbalanced cell-type compositions in scRNA-seq integration settings, and demonstrated that cell-type proportion imbalance leads to skewed distributions in standardized gene expression values between datasets. This drives major changes in the dimensionality reduction step in scRNA-seq analysis, and subsequently leads to inaccurate integration results [10]. Currently, no existing study has quantified the effects of dataset imbalance on both integration results and downstream biological conclusions. This aspect is highly relevant, as mechanisms to account for dataset imbalance do not readily exist in frequently utilized integration techniques [9, 13].

Here, we present an extensive analysis of the effects of dataset imbalance on scRNA-seq data integration. We begin by examining two balanced scRNA-seq batches of human peripheral blood mononuclear cell (PBMC) data [9, 14, 15] as a controlled setting. To determine the effects of dataset imbalance on integration results and downstream analyses, we perform 1600 perturbation experiments using the *Iniquitate* pipeline that involve control, downsampling, and ablation simulations in a cell-type-specific manner with replicates. Downstream analyses tested include unsupervised clustering [8], differential expression to determine marker genes [8], nearest-neighbor-based cell-type classification [16], and query-to-reference cell-type annotation [17]. To extend the analyses to more complex settings, we analyze datasets with prevalent imbalance, including imbalanced PBMC datasets [18], longitudinal mouse hindbrain developmental data [19], and pancreatic ductal adenocarcinoma (PDAC) samples from different patients [20]. Our analyses reveals that dataset imbalance has cell-type-specific effects on integration performance, as well as the downstream results, and that these effects are largely method-agnostic. We further define two key aspects of multi-sample single-cell data that act in concert to affect downstream results - *relative cell-type support* and *minimum cell-type center distance*. To address limitations with respect to dataset imbalance in benchmarking single-cell integration, we reformulate current integration metrics to consider imbalance explicitly. Finally, we provide a series of guidelines and recommendations to help minimize and mitigate the impacts of dataset imbalance in scRNA-seq integration settings.

## Results

### Development of a comprehensive perturbation pipeline to determine the impacts of imbalance in scRNA-seq integration

To assess the impacts of dataset imbalance in scRNA-seq integration, we developed a pipeline termed *Iniquitate*, that quantifies imbalance prevalent in datasets using global and per-cell-type statistics, determines the differences in these quantities between samples/batches, and tests the effects of down-sampling perturbations on integration and downstream analysis results (Figure 1A). Datasets utilized were annotated by experts in their respective studies, with the exception of the PDAC data which was re-annotated to better identify malignant cells (Online Methods). We tested five state-of-the-art scRNA-seq integration methods, including BBKNN [21], Harmony [22], Scanorama [23], scVI [24] and Seurat [25]. A uniform integration pipeline embedded within *Iniquitate* was utilized to make comparisons between methods and across datasets comparable, with some noted exceptions (Online Methods). We measured cell-type heterogeneity conservation and batch effect correction for each technique across datasets and perturbations using the Adjusted Rand Index (ARI) [26], Adjusted Mutual Information (AMI) [27], Homogeneity Score [28], and Completeness Score [28] (Online Methods).

**Figure 1:**
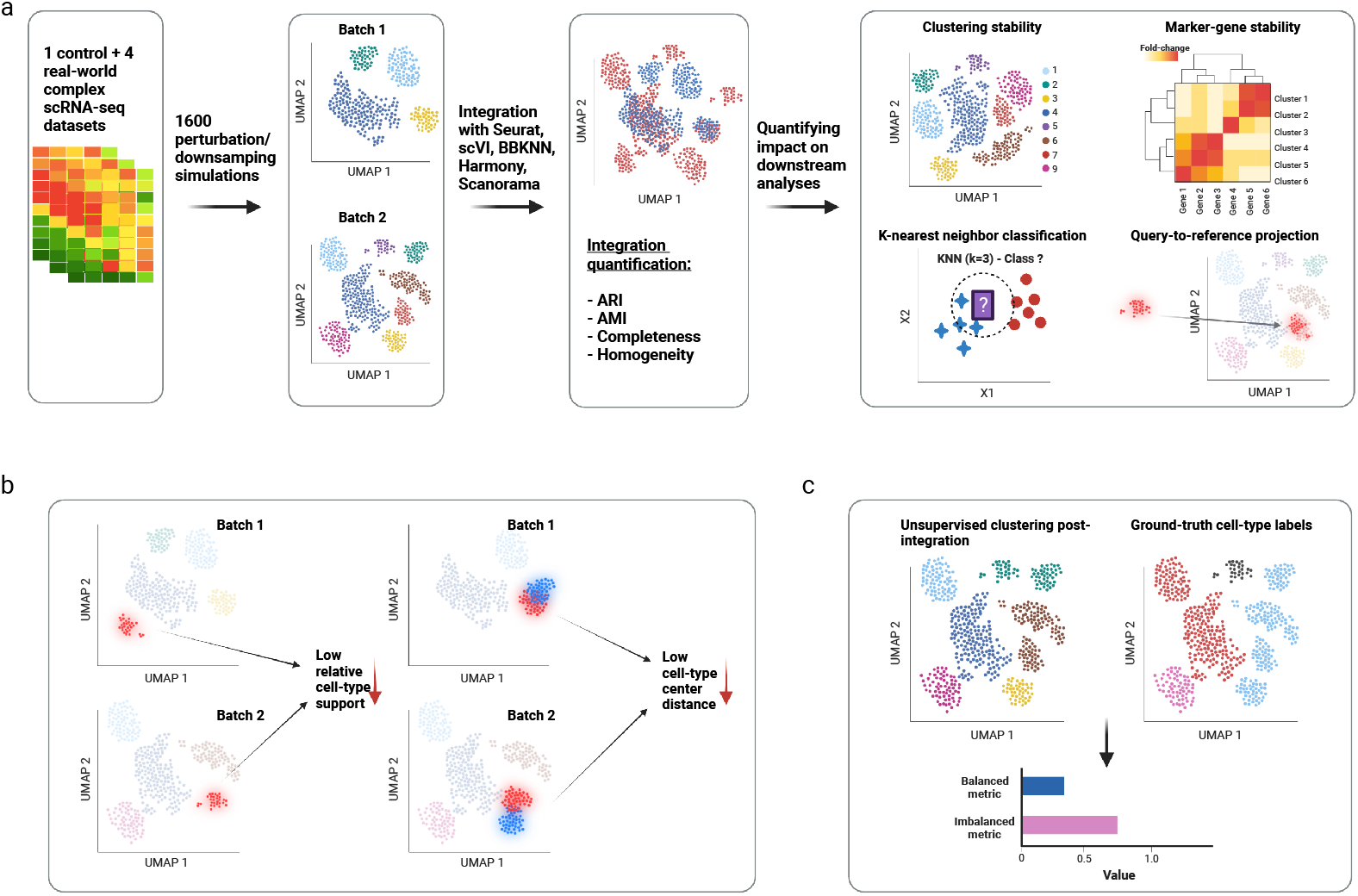
Overview of the Iniquitate pipeline and analysis results. (a) To determine the effects of dataset imbalance in scRNA-seq integration, 1 controlled balanced PBMC dataset and 4 complex datasets with imbalance already present were integrated using current state-of-the-art scRNA-seq integration techniques. A total of 1600 perturbation experiments involving downsampling on the controlled dataset were performed and the effects of imbalance on integration results as well as downstream analyses (clustering, differential gene expression, cell-type classification, query-to-reference prediction) were quantified. (b) In complex datasets, results in the controlled setting were verified, and two key data characteristics were found to contribute to altered downstream results in imbalanced settings - relative celltype support and minimum cell-type center distance. (c) To account for imbalanced scRNA-seq integration scenarios in evaluation and benchmarking, typically utilized metrics and scores were reformulated to reweigh disproportionate cell-types, which includes the Balanced Adjusted Rand Index (bARI), Balanced Adjusted Mutual Information (bAMI), Balanced Homogeneity Score, and Balanced Completeness Score.

To determine the impacts of dataset imbalance on downstream analyses, we analyzed post-integration impacts on unsupervised clustering [8, 29], cell-type classification [16], differential gene expression [8] [30], and query-to-reference annotation [17] results (Figure 1A). Clustering impacts were assessed based on changes in the number of clusters post-integration using unsupervised clustering (Figure 1A). To assess the impacts on cell-type classification, a nearest-neighbor cell-type classifier was trained on the post-integration embeddings and tested on a holdout set (Figure 1A). Differential gene expression results and variation was assessed using a global importance metric of the ranking changes of marker genes specific to each cell-type analyzed, before and after perturbing the dataset balance (Figure 1A). Query-to-reference annotation was done using the Seurat 4.0 method [31], which projects each batch to be integrated onto a reference scRNA-seq dataset, and accuracy of annotation was utilized as an endpoint (Figure 1A). The details of the evaluations utilized for all of the downstream analyses and parameters of the integration pipeline are outlined in Online Methods.

Through perturbation and cell-type-specific analysis of both balanced and complex imbalanced scRNA-seq datasets, we determined that cell-type imbalance affects the scores of typical integration metrics in a cell-type and method-specific manner. Further, we discovered that cell-type imbalance in datasets to be integrated can lead to significant deviations in the results of downstream analyses. After investigating factors of imbalance that can quantifiably lead to distinct downstream results, we found that the distance between cell-types in the embedding space (*minimum cell-type center distance*) and imbalance between cell-types (*relative cell-type support*) to be the most relevant and predictive in this regard (Figure 1B). Finally, we determined that typical clustering metrics utilized in benchmarking single-cell integration techniques, such as ARI and AMI, are inadequate in imbalanced scenarios as they weigh the more prevalent cell-types disproportionality compared to rare cell-types. Therefore, we develop and introduce novel balanced clustering metrics, including the *Balanced Adjusted Rand Index* (bARI), *Balanced Adjusted Mutual Information* (bAMI), *Balanced Homogeneity Score*, and *Balanced Completeness Score* (Figure 1C). The balanced metrics reweigh the base scores such that each ground-truth cell-type’s contribution to the score is considered equally.

### I. Perturbation-induced imbalance in a PBMC cohort indicates cell-type-specific effects on integration results

The ideal test case for assessing impacts of dataset imbalance should begin with a balanced dataset as a baseline, and thus we analyzed a peripheral blood mononuclear (PBMC) cohort of two batches/samples processed independently from two different healthy donors [9, 14, 15]. We downsampled each batch to have 6 major cell-types and an equal number of cells within each cell-type (400 cells for each cell-type) (Figure 2A) (Online Methods). The cell-types were selected such that they are equivalent between the batches. Therefore, the cell-types present, number of cells per cell-type, and cell-type proportions between the batches are equal and the integration scenario is balanced (Figure 2A, Figure 2B). A batch effect is prevalent between the samples (Figure 2B), which is expected as they were processed at different centers using different technologies (10x 3’ vs 5’ protocols - Online Methods) [14, 15]. In this balanced setup, we aimed to assess how the integration results of the two PBMC batches, from a typical integration metric standpoint as well as their impacts on downstream analyses, varied between the control balanced data and perturbation-induced imbalanced data. For each perturbation, we randomly selected one of the two batches and one cell-type within the selected batch to either downsample to 10% of the original population or ablate/remove completely from the selected batch (Figure 2C). These perturbations were repeated 400 times for both downsampling and ablation of a random batch/cell-type. Control experiments with no perturbations to the balanced data were repeated 800 times, resulting in 1600 integration experiments where each integration technique was tested (Online Methods).

**Figure 2:**
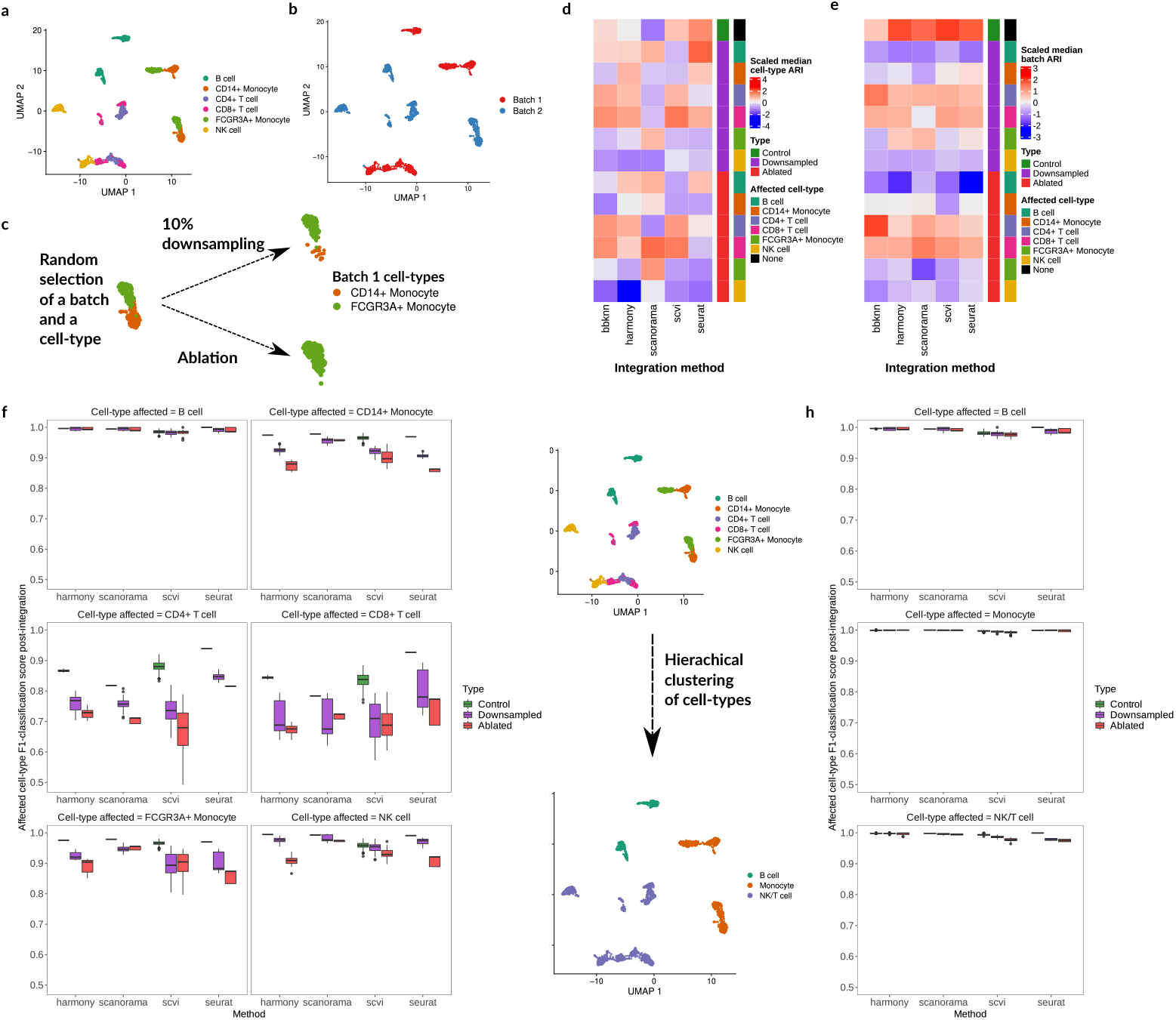
Perturbation analysis of controlled PBMC dataset and effects on cell-type-specific integration. (a), (b) The cell-type and batch representations of the balanced two-batch PBMC dataset.(c) The perturbation setup for the balanced PBMC data - in each iteration, one batch and one cell-type is randomly selected, and the cell-type is randomly either downsampled to 10% of its original number or ablated. Control experiments are also performed where no downsampling occurs. (d), (e) Z-score normalized median ARI_*cell–type*_ (cell-type integration accuracy) (d) and median (1-ARI_*batch*_) (batch mixing) (e) results across experiment type (control, cell-type downsampling, cell-type ablation), specific-cell-type downsampled, and integration method utilized. (f) KNN-classification within the integrated embedding space in control, downsampling and ablation replicates and across methods. The F1-scores are indicated for the same cell-type that was downsampled. (g) Hierarchical clustering of similar cell-types in the balanced two-batch PBMC data. (h) Cell-type-specific integration results using a KNN-classifier after hierarchical clustering across perturbation experiments with the same setup as (f). The cell-types here are based on the label after hierarchical clustering from (g).

To determine how cell-type-specific changes in dataset balance affected typical integration metrics, we examined the ARI_*cell–type*_ and (1 - ARI_*batch*_) scores [9] for each run and method independently. ARI_*cell–type*_ represents conserved heterogeneity of annotated cell-types post-integration, and (1 - ARI_*batch*_) represents the degree to which the two batches being integrated overlap post-integration [9]. As variation between methods was not the main objective of the analysis, the two scores were Z-score normalized for each method across the perturbation experiments and the median value was utilized due to the presence of replicates (Online Methods). Neither the scaled median ARI_*cell–type*_ or scaled median (1 - ARI_*batch*_) indicated distinct patterns for the perturbation experiments (Figures 2D, 2E). In fact, there seemed to be a high degree of method-specific variation in these results, making the interpretation challenging. In terms of the median (1 - ARI_*batch*_) scores, for 4 out of 5 methods the top score occurred in the control setup which shows that imbalance leads to worsening performance in terms of batch-mixing (Figure 2E). The results for cell-type heterogeneity were even less clear and indicated differences based on both the method utilized and cell-type downsampled/ablated. Overall, the results did not contain clear patterns and point to the fact that global clustering metrics do not account for dataset balance and may not be adequate for assessing performance in scenarios with imbalanced datasets and rare cell-types.

To overcome this limitation of global metrics, we examined integration performance at a cell type-specific level through a k-nearest-neighbor (KNN) classifier [16, 32] that was trained on 70% of the post-integration embeddings from each method independently and the remaining 30% was used as a test-set for cell-type classification. The train/test split was stratified by cell-type label, such that an equal proportion of cell-types occurred in both subsets, allowing for comparison of classification at the cell-type level (Online Methods). Overall, the classification results provide evidence for cell-type-specific effects of dataset imbalance, as downsampling a specific cell-type led to a statistically significant decrease in the KNN classification F1-score [33] for the same cell-type post-integration, based on an analysis-of-variance (ANOVA) model [34] (ANOVA *p*-value << 0.05, F-statistic = 1304.96, Supplementary Figure S1) (Figure 2F). This result is method agnostic as the ANOVA test factored in method utilized and cell-type downsampled (Online Methods). The only cell-type that exhibited stability were B cells (Figure 2F - standard deviation of median F1-score across methods and experiment types < 0.01). Comparing the cell-type-specific results with the global ARI metrics, we found weak correlation across all methods (Supplementary Figures S2, S3, Spearman’s *ρ* ≤ 0.4 across methods and metrics). The uniformly worsening F1 classification scores for the majority of cell-types being perturbed, when compared with the global ARI metrics, shows that global metrics may not adequately capture the integration performance in imbalanced settings. Instead, cell-type-specific metrics such as the KNN-classification score can capture more granular information.

We hypothesized that the integration performance for B cells was unaffected in the perturbation experiments because they are highly distinct from the other cell-types. The two monocyte subsets (CD14+ Monocytes and FCG3RA+ Monocytes) are transcriptionally similar, and the two T-cell subsets (CD4+ T cells and CD8+ T cells) and NK cells are also very similar (Figure 2A, Supplementary Figure S13). As a test, we performed hierarchical clustering of the cell-types into three higher-level subsets - B cells, Monocytes, and NK/T cells (Figure 2G) (Online Methods). As expected, downsampling these subsets did not result in worsening performance to the same degree as the base cell-types (Figure 2F, Figure 2H) (ANOVA F-statistic = 374.46 (hierarchical) << 1304.96 (base), Supplementary Figure S1) (Online Methods). This initial result on the balanced PBMC cohort indicates that the relative transcriptomic similarity of cell-types can drive cell-type-specific performance of integration techniques when considering differing levels of dataset imbalance.

### II. Biological interpretation of integration results is contingent on relative cell-type proportions between batches

To further analyze the impact of the perturbation experiments on the balanced PBMC cohort, we quantified the effects of imbalance on downstream analyses typically performed after integration, including unsupervised clustering, differential gene expression/marker gene selection, and query-to-reference annotation (Figure 1A). As we observed significant impacts on KNN-based cell-type classification in the same setting, it is likely that the impacts of imbalance on integration may also affect other aspects of single-cell analysis. Therefore, we utilized the same perturbation setup and downsampling experiments in the balanced PBMC cohort to analyze these effects.

#### Stability of unsupervised clustering of samples post-integration

We observed a significant variation in the inferred number of clusters after integration across all tested methods due to perturbation of cell-type balance (ANOVA *p* << 0.05, F-statistic = 990.79) (Figure 3A). After integration in both balanced and perturbed simulations, clustering was performed using the Leiden clustering algorithm with a fixed resolution (Online Methods). Although all methods indicated at least some degree of variation in the number of clusters between control and downsampled/ablation experiments, there were also method-dependent effects present (Figure 3A). For instance, while Harmony exhibited variation in the number of clusters regardless of cell-type downsampled, ablation of CD14+ Monocytes specifically led to a much smaller number of clusters overall post-integration (Figure 3A). A similar effect was observed for Seurat and BBKKN, while Scanorama’s post-integration clusters diverged most from the control experiments when ablating CD4+ and CD8+ T cells (Figure 3A). scVI’s post-integration clustering results were relatively more stable after perturbation (Figure 3A). There was variation observed for the control experiments across methods as well, but clear deviation after perturbation was present in all tested methods. This result indicates that differing levels of imbalance can cause significant deviations in cluster number, even though the number of clusters should be stable as the number of cell-types across all batches remains the same in both perturbed and unperturbed experiments. As cell-types are typically annotated using unsupervised clustering and subsequent marker gene analysis [8, 17], varying degrees of dataset imbalance can lead to distinct biological conclusions.

**Figure 3:**
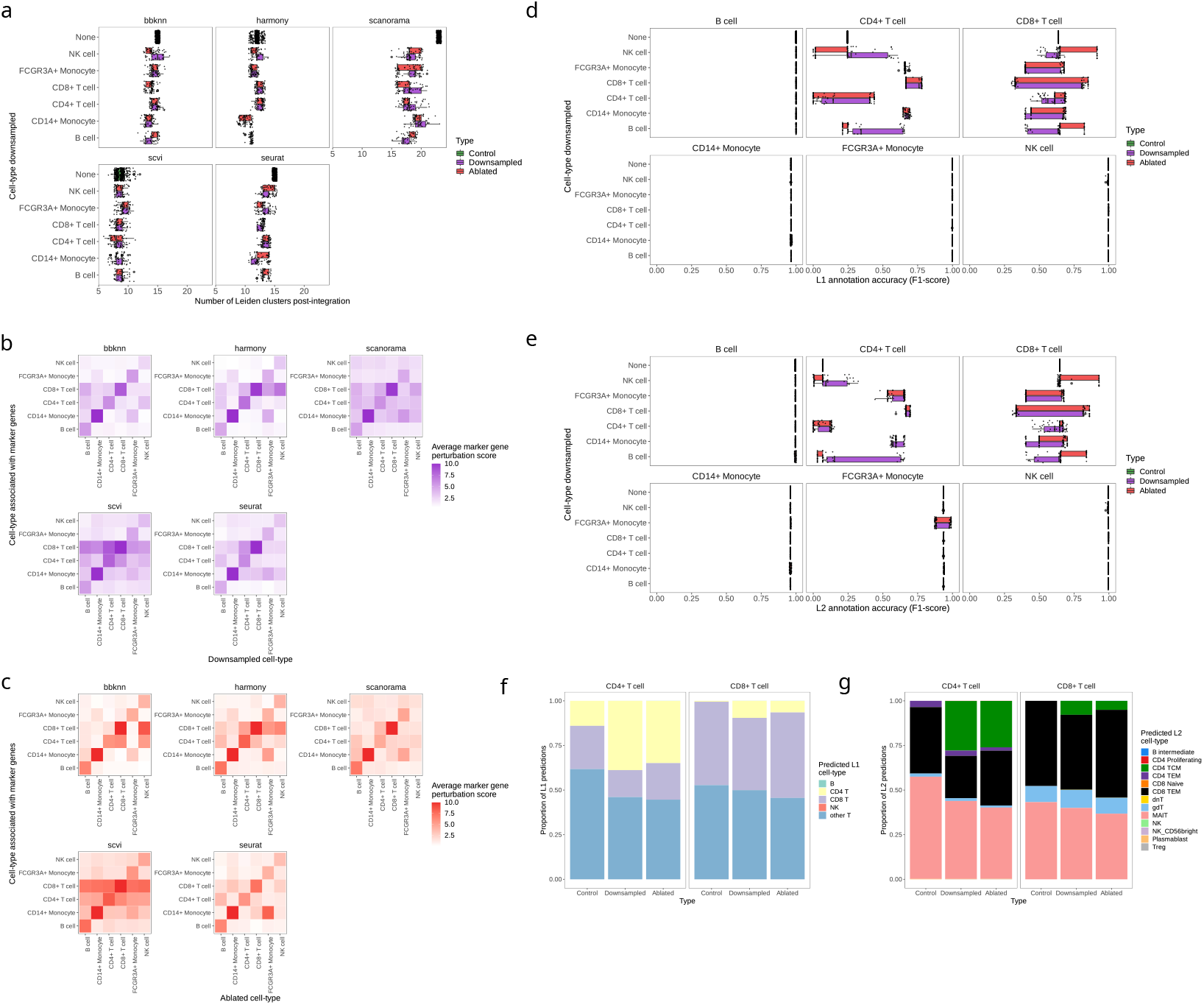
Quantification of the effects of perturbation-induced dataset imbalance on downstream analyses. (a) After integration of the PBMC balanced dataset in different perturbation scenarios (type) and based on the cell-type downsampled, the number of unsupervised clusters from the results of each method based on Leiden clustering across replicates. (b) The average marker gene ranking change in differential gene expression (average marker gene perturbation score) for cell-types downsampled and marker gene sets of specific cell-types, across methods. The rankings are averaged across replicates for the ‘downsampled’ experiment type. (c) The average change in marker gene ranking in differential gene expression averaged across replicates for the ‘ablation’ experiment type. (d), (e) The cell-type-specific L1 annotation (coarse-grained) (d) and L2 annotation (fine-grained) (e) accuracy scores across replicates for query-to-reference results for individual batches based on experiment type (control, downsampling, ablation) and cell-type downsampled. (f), (g) The L1 predictions (f) and L2 predictions (g) by proportion across experiment types and replicates for CD4+ T cells and CD8+ T cells.

An important caveat to this result is that a reduced number of total cells (through downsampling or ablation) might lead to less clusters in general at a fixed resolution due to less overall heterogeneity in the data. We argue that this is not a major limitation due to two factors - (1) This case is still reflective of the effects of perturbing the cell-type balance, even if the effects are uniform across cell-types downsampled, (2) We did not observe a uniform reduction in cluster number based on the integration methods utilized. For example, scVI’s results for cluster number were fairly stable after downsampling or ablation, while the results from Scanorama indicated a drastic reduction after perturbation (Figure 3A). Moreover, within each method, the results for reduction in cluster number were not uniform based on the cell-type that was downsampled (Figure 3A). Therefore, although we are limited in evaluation at a fixed clustering resolution, the results nevertheless show that the perturbation setup can lead to cell-type and method-specific effects that can potentially alter further analyses.

#### Differential gene expression and marker gene stability

Frequently, the next step after integration and unsupervised clustering in a scRNA-seq analysis workflow is differential gene expression analysis [8, 35]. Typically, a series of one-versus-all differential expression experiments, using statistical tests such as the non-parametric Wilcoxon Rank-Sum Test or more RNA-seq specific techniques such as DESeq2, is done for each cluster to determine the top ranking “marker genes” specific to all clusters [8, 17, 35]. These marker genes are indicative of cell-type identity for each cluster and are used to annotate clusters into putative cell-types [8, 17, 35]. One way to assess marker gene stability before and after perturbation is to constrain the number of clusters to be equivalent across simulations, but this would be unrealistic as variation in cluster number in both control and perturbed experiments was observed across methods (Figure 3A). As the *ranking* of marker genes is typically utilized to annotate clusters from scRNA-seq data [8, 35], we considered deviation in ranking for genes with known cell-type associations to be an important end-point. Using the unintegrated data separately for each batch, we determined the top 10 marker genes for each cell-type, and assessed the stability of their ranking before and after perturbation (Online Methods). Changes in ranking for marker genes across replicates for a given subset of experiments were defined as the *marker gene perturbation score*, indicating the standard deviation of the rank (Online Methods). In the case of examining all marker genes for a given cell-type, the standard deviation of ranking of all of the marker genes was averaged, and this is indicated as the *average marker gene perturbation score* (Online Methods).

For the majority of marker genes, we observed deviations in ranking after downsampling and ablation, with many diverging as much as 10 ranks (Supplementary Figure S4 - Marker gene perturbation score) which could lead to significant changes in biological interpretation of results if the top 10 marker genes are used as a heuristic for annotation. An ANOVA test factoring in the specific marker gene, method, and downsampled cell-types indicated that perturbation led to statistically significant changes in ranking (ANOVA *p* << 0.05, F-statistic = 48.99 - highest of all factors) (Online Methods). There was strong correlation in marker gene perturbation across methods, with the exception of scVI, which exhibited stronger deviations in rankings for some marker genes (Supplementary Figure S4).

Next, we examined whether downsampling or ablation of a specific cell-type will change the ranking of marker genes for the same cell-type, and we observed that this was the case across all methods (Figure 3B, Figure 3C). The strongest ranking change of marker genes occurred after downsampling or ablation of CD8+ T cells and CD14+ Monocytes (Figure 3B, Figure 3C). As these two cell-types are highly similar to CD4+ T cells and FCGR3A+ Monocytes respectively, downsampling likely induces a collapse of cells in the downsampled cell-types into clusters corresponding to their neighboring cell-types. This contributes to significant deviations in marker gene ranking and possible changes in biological interpretation in both the downsampling and ablation experiments. Significant changes in marker gene ranks were also observed for cell-types that were not downsampled or ablated, such as in NK cells, which were pronounced for Harmony and scVI results (Figure 3B, Figure 3C). Once again, this is likely due to mixing of cell-types within clusters after an imbalance is introduced, as NK cells are very transcriptionally similar to CD4+ and CD8+ T cell subsets. Similarity of cell-types and effects in integration are investigated further in Results III. and IV.

To more definitively determine whether or not these perturbations in marker gene rankings could change the biological conclusions of an analysis, we performed a case-study with clusters that contained a majority of CD4+ and CD8+ T cells after integration using Seurat (Supplementary Figure S16). Considering a permissive list of the top 50 marker genes and using canonical markers for CD4+ and CD8+ T cells, we observed that the fraction of clusters annotated as either CD4+ or CD8+ T do change after downsampling/ablation induced imbalance is introduced (Supplementary Figure S16).

#### Query-to-reference projection and cell-type annotation

With the increasing availability of public scRNA-seq datasets with high quality annotations, query-to-reference annotation has become a major application for scRNA-seq data integration [17]. However, the accuracy of annotation depends on the quality of the integrated space. To examine the effects of imbalance in this setting, we utilized the Seurat 4.0 query-to-reference annotation pipeline and a large-scale multi-modal PBMC dataset of 211 000 cells as a reference [31]. In the Seurat 4.0 pipeline, each batch (query) is projected to the reference dataset, such that the integration is performed individually for each batch [31] (Online Methods). In this setup, the effects of inter-batch imbalances are not relevant, but only the imbalance relative to each query batch and the reference dataset. In this setting, the perturbations were done for the query batches (balanced PBMC 2 batch data), and the reference was static (Online Methods). We assessed the accuracy of query-to-reference projection through a “fuzzy-matching” of cell-type labels between the balanced PBMC batches and multi-modal PBMC reference from Seurat [9, 31] (Online Methods).

The majority of cell-types were stable across control and downsampling/ablation experiments with near perfect scores. However, the two T-cell subsets had varying performance to a high degree, regardless of which cell-type was downsampled or ablated (Figure 3D, Figure 3E). This result is indicative of the fact that the imbalance between the projected batch (which was perturbed) and the reference dataset (held constant across all experiments) is driving variance in integration and subsequent annotation results. This highlights a similar problem concomitant with previous results, in that perturbing the degree of balance for transcriptionally similar cell-types can lead to biologically distinct results compared to the balanced scenario. In this case, the CD4+ T cell and CD8+ T cell populations are transcriptionally similar, and a trade-off in their annotation performance can be observed in the control unperturbed data (Figure 3D, Figure 3E). After perturbing the degree of balance within a given batch, the trade-off point is moved in favor of either subset (Figure 3D, Figure 3E). Further, the result highlights that perturbation of dataset balance can affect downstream results even when there is a degree of imbalance already present between integrated datasets, which was the case between the query and reference data in these experiments (Supplementary Table 3, Supplementary Table 9).

Examining the cell-type annotations more closely at two levels of resolution, we observed that both the CD4+ and CD8+ T cells were largely mis-annotated as Mucosal Associated Invariant T-Cells (MAIT) (Figure 3F, Figure 3G). After downsampling or ablation of a given cell-type and subsequent analysis of annotation accuracy of the same cell-type, we find that CD4+ T cells were annotated more accurately, while CD8+ T cells were further mis-annotated, compared to their respective control scores (Figure 3F). The transcriptional similarity between not just the CD4+/CD8+ subsets, but the many subsets that fall under “other T”, is a challenging problem for integration and subsequent label-transfer [36]. This challenge is potentially exacerbated when imbalance is present, as indicated by the perturbation experiments and their effects on the annotation results.

Overall, cell-type imbalance affected all three major aspects of downstream analysis that were tested, and we observed strong evidence of impact on biological interpretation of the results. This observation is likely even more relevant in complex datasets, as the balanced PBMC co-hort is not representative of the ever-increasing throughput of current scRNA-seq protocols [37]. The limitations of the reference dataset utilized may also be a major source of variation in the query-to-reference integration results. It may be the case that a more suitable reference may not lead to high variance in the results of the two T-cell subsets, however assessment and selection of reference datasets is outside the scope of this study and the multi-modal PBMC reference used is one of the most comprehensive single-cell references to date.

### III. Analysis of imbalanced complex datasets reveals key metrics for stability of integration results

While perturbation experiments of the balanced two batch PBMC cohort revealed the effects of dataset imbalance on integration and downstream analyses in a controlled setting, current scRNA-seq datasets typically involve a much larger number of cells and cell-types captured [37]. Therefore, we examined the effects of dataset imbalance when integrating complex datasets with multiple samples that are not perturbed, but already have inherent cell-type imbalance between samples. To this end, we analyzed an imbalanced 2 batch PBMC cohort [18], imbalanced 4 batch PBMC cohort [18], imbalanced cohort of 6 batches of mouse hindbrain developmental data [19], and an imbalanced heterogeneous tumor cohort of 8 batches of pancreatic ductal adenocarcinoma (PDAC) data [20] (Online Methods). No downsampling was done in these experiments and we aimed to analyze the effects of integration on cell-types that are imbalanced with respect to others, both within and across batches.

As determined in the analysis using the balanced PBMC data and perturbations, transcriptomic similarity (relatedness) of cell-types and cell-type imbalance are two important factors that impact downstream results. We sought to observe if these properties also led to differences in integration performance per-cell-type in more complex datasets without perturbations. We formalized these two properties as the *relative cell-type support* and *minimum cell-type center distance*, quantifying the degree of imbalance and relatedness to other cell-types (Online Methods). The *relative cell-type support* is defined as the number of cells specific to a cell-type present across all batches, and the *minimum cell-type center distance* considers the average distance across all batches between cell-types in a principal component analysis (PCA) dimensionality reduction representation space and selects the distance of the closest neighboring cell-type for each cell-type (Online Methods).

To correlate these properties with integration performance, we used the same KNN-classification setup as before to determine performance on a per-cell-type basis (Online Methods). We started by analyzing the cell-type center distances on the imbalanced PBMC 2 and 4 batch datasets. Examining the average cell-type center distances - which were averaged across all batches if the cell-types were present in more than one batch - there is a clear pattern evident between cell-types that were observed before in the 2 batch balanced PBMC data (Figure 4A, Figure 4B). NK cells have small relative distance between the T cell subsets, while dendritic cells share transcriptional similarity with the monocyte subsets (Figure 4A, Figure 4B). B cells and megakaryocytes have the greatest distance between the rest of the cell-type centers (Figure 4A, Figure 4B), and thus we expect these cell-types to have strong performance in integration which was previously observed for B cells. This pattern did hold when examining integration performance through KNN-classification for both datasets on a per cell-type basis compared to their *minimum cell-type center distance* across batches, as Megakaryocytes and B cells had strong performance regardless of integration technique utilized (Figure 4C, Figure 4D).

**Figure 4:**
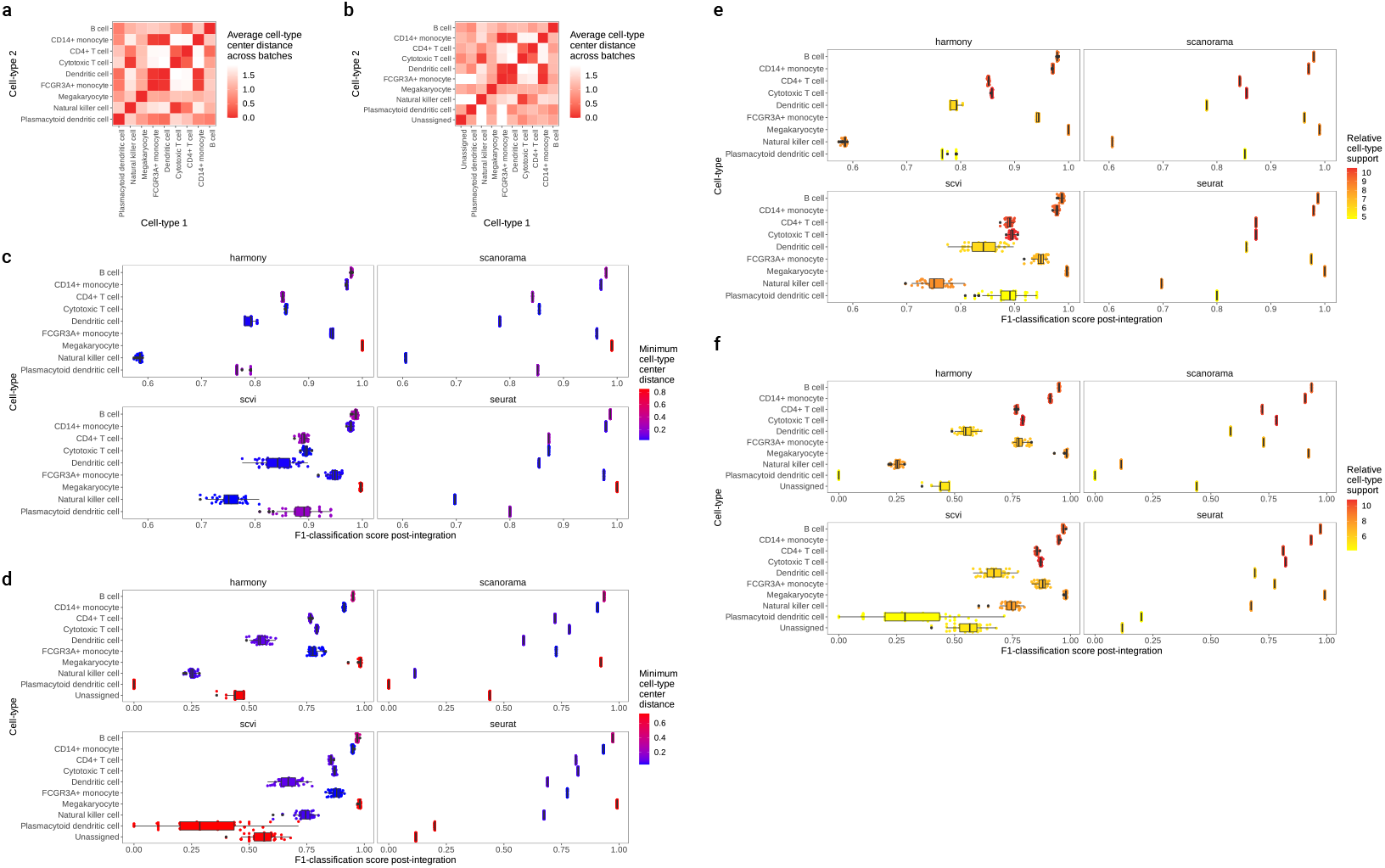
Factors in imbalanced complex datasets predictive of altered integration and downstream results. (a) The average cell-type center distance across cell-types in the imbalanced PBMC 2 batch dataset. For each batch, the distance from the centers of cell-type clusters in principal component analysis (PCA) reduction space are calculated, and the relative distances between cell-types are determined and averaged across batches. (b) The average cell-type center distance across cell-types in the imbalanced PBMC 4 batch dataset. (c), (d) Comparison of F1-classification accuracy of each cell-type in the imbalanced PBMC 2 batch dataset (c) and imbalanced PBMC 4 batch dataset (d), specific to method and across replicates, compared with the *minimum cell-type center distance* value. (e), (f) Comparison of F1-classification accuracy of each cell-type in the imbalanced PBMC 2 batch dataset (e) and imbalanced PBMC 4 batch dataset (f), across methods and replicates, compared with the *relative cell-type support* value. The *relative cell-type support* is based on the number of cells in the integrated embedding space present for each cell-type.

However, the results were not straightforward for other cell-types. Examining the plasmacytoid dendritic cells, we expected strong performance due to their high relative distance between other cell-types, but this was not the case for both the 2 and 4 batch datasets (Figure 4C, Figure 4D). Although these cells have a large relative distance between other cell-types, they occur in a much smaller number compared to others (Figure 4E, Figure 4F). We quantified *relative cell-type support* as the log-transformation of the total number of cells for each cell-type across batches (Online Methods), and plasmacytoid dendritic cells have the lowest value across cell-types within these two datasets (Figure 4E, Figure 4F). This result indicates that *minimum cell-type center distance* is necessary but not sufficient to explain variation in integration results on a per-cell-type basis. Overall, higher *relative cell-type support* does seem to lead to higher performance in integration for some cell-types, such as the CD14+ and CD16+ Monocyte subsets, but is also not sufficient for higher performance, as NK cells perform poorly across integration techniques due to having a low *minimum cell-type center distance* and overlap with the T-cell subsets despite not having low *relative cell-type support*. Examining the results across both metrics and performing an ANOVA test to determine variance in scores explained by the metric, we did find statistically significant associations for both *minimum cell-type center distance* and *relative cell-type support* [ANOVA *p*-value << 0.05 for both metrics across all datasets, with the exception of the mouse hindbrain 6 batch dataset, for which *relative cell-type support* is non-significant (Supplementary Table 1) (Online Methods)].

Analysis of the 6 batch mouse hindbrain developmental data and 8 batch PDAC datasets indicated similar results, albeit much less easily interpretable due to the presence of a very large number of cell-types and more batches (Supplementary Figures S5–S8). Cell-types in close proximity within an embedding space have an interpretable explanation for poor integration performance, as they may collapse and become merged with their overlapping counterpart in the integration step. This was observed in Results sections I and III. Low cell-type support leads to less data for a given cell-type that an integration method/model can utilize, and therefore models may not be able to learn the correct embedding for these cell-types.

### IV. Perturbation analysis in PDAC samples reveals tumor compartment-specific effects of dataset imbalance

To further analyze the effects of dataset imbalance in realistic scenarios, we considered the pancreatic ductal adenocarcinoma (PDAC) dataset of 8 batches comprising tumor samples across 8 different biopsies [20]. One major challenge in the analysis of PDAC data is accurate annotation of tumor cells, and being able to separate these from normal non-cancerous epithelial cells [38, 39]. As both acinar and ductal epithelial cells have been proposed as cell of origin candidates in PDAC across numerous studies [40, 41], reliably classifying tumor cells from these normal epithelial cell-types in scRNA-seq data remains a major computational challenge. Given this difficulty, we sought to determine if different levels of imbalance between epithelial normal and epithelial tumor compartments can influence the accuracy of PDAC tumor tissue integration and subsequent classification of tumor cells. As scRNA-seq data from tumor tissue is often integrated across multiple biopsy sites, patients, and cohorts [42], the ability to reliably quantify tumor cells is imperative to the biological validity of subsequent downstream analyses. We preprocessed and annotated tumor cells in the PDAC samples through integration, unsupervised clustering, and marker gene-based annotation (Online Methods). In setting up the perturbation experiments, we grouped epithelial normal cells (acinar and ductal) into an “epithelial normal” compartment, tumor cells into “epithelial tumor” compartment and the remaining microenvironment cells into the “microenvironment” compartment (Figure 5A) (Online Methods). Overall, the microenvironment heavily outnumbered the epithelial tumor and epithelial normal populations (Figure 5B), which is reflective of the low tumor purity typical of PDAC biopsy samples [20]. Perturbation experiments included downsampling or ablation of a randomly selected compartment within 4 randomly selected batches out of 8 (Figure 5A, Figure 5B). We also performed replicates for control, downsampling, and ablation, for a total of 200 simulations for integration of all 8 batches (Online Methods).

**Figure 5:**
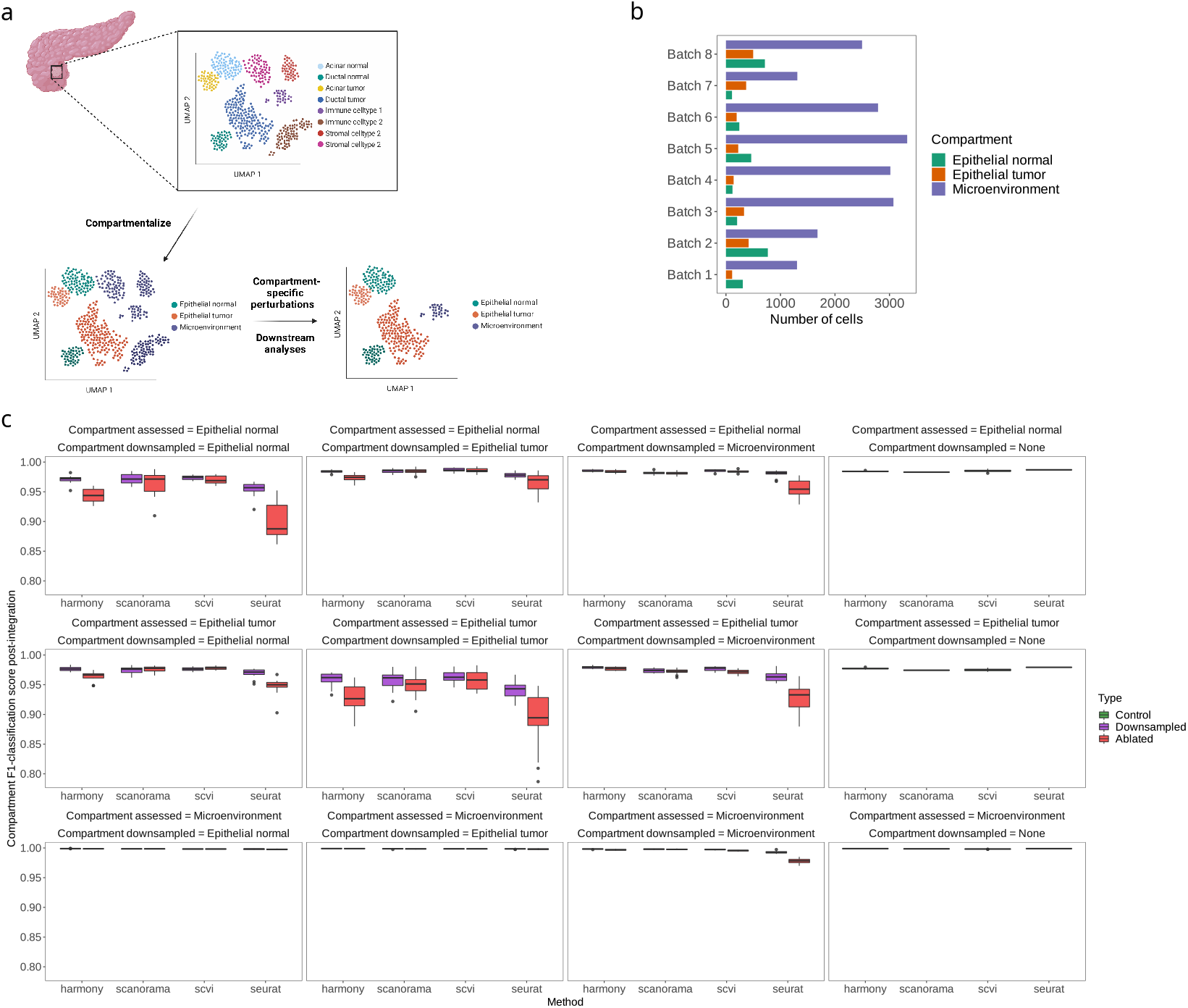
Compartment-wise perturbation experiments for 8 batches of PDAC samples. (a) Overview of the experimental setup. To determine the effects of dataset imbalance across tumor compartments, various microenvironment tumor were collapsed into the ‘microenvironment’ compartment, normal ductal and acinar cells into the ‘epithelial normal’ compartment, and malignant ductal and acinar cells into the ‘epithelial tumor’ compartment. The perturbation experiments involved the sample downsampling (10% of a compartment) and ablation (complete removal of a compartment) setup for 4/8 randomly selected batches. Note that all batches are integrated at once using each method. (b) Number of cells in each compartment after cell-type collapse, across batches/biopsy samples in the PDAC data. (c) F1-classification score for KNN classification post-integration, specific to each compartment when compared with the compartment that was downsampled or ablated, across replicates and methods utilized for integration.

Examining the KNN-classification scores on a per-compartment assessed, per-compartment downsampled basis indicated that downsampling or ablating the microenvironment compartment leads to stable compartment classification across all methods, with a slight decrease in performance observed for Seurat in the epithelial normal and tumor compartments (Figure 5C). This result is concordant with previous analysis indicating that proximity is a key factor that dictates the degree to which perturbations in cell-type balance can affect integration results. The *minimum cell-type center distance* in the PDAC data shows that acinar and ductal cells, which comprise the epithelial normal and epithelial tumor populations, are two of the most distant cell-types from others in the data (Supplementary Figure S8, Supplementary Figure S15)). Similar to the discrepancy between ARI_*cell–type*_ and (1 - ARI_*batch*_) observed in the integration of the balanced and imbalanced PBMC datasets, we observed that higher ARI_*compartment*_ based on downsampling of the microenvironment did not lead to higher batch mixing scores (Supplementary Figure S9, Supplementary Figure S10). In fact, we observed the opposite effect almost uniformly across all methods, as downsampling the epithelial compartments decreased the ARI_*compartment*_ and increased (1 - ARI_*batch*_) (Supplementary Figure S9, Supplementary Figure S10). Downsampling the microenvironment had the opposite effect. As the microenvironment is quite large, downsampling likely leads to decreases in batch mixing scores because these metrics are driven by more prevalent compartments/cell-types. Epithelial cells and their tumor/normal dichotomy is of strong interest in analyzing PDAC data, and therefore batch mixing is likely a poor quantifier of integration performance and the biological validity and utility of the results. This result also reiterates the limitation of global clustering metrics that do not take into account less prevalent cell-types and their overall difficulty in interpretation, as the increased performance in ARI_*compartment*_ after downsampling the microenvironment was not concordant with the KNN-classification results (Supplementary Figure S9, Figure 5C).

Tumor and normal epithelial compartment KNN-classification scores worsened as either compartment was downsampled (Figure 5C). More specifically, downsampling either the epithelial normal or epithelial tumor compartments led to the greatest decrease in the integration performance of the same compartment through the KNN-classification setup (Figure 5C) (ANOVA F-statistic_*Normal epithelial*_, F-statistic_*Tumor epithelial*_ > F-statistic_*Microenvironment*_ - Supplementary Figure S11) (Online Methods). This indicates that relative proportions of tumor to normal cells can lead to differing results in integration performance in this setting, which is reflective of the earlier observations in the balanced PBMC dataset with highly similar populations, such as the NK and T cell subsets. Overall, these results demonstrate that the degree of imbalance between the similar compartments across tumor tissue cohorts can significantly affect the downstream results and possibly subsequent analyses. This result is not specific to PDAC data, as tumor samples across cancer types share typical characteristics, but may not have the same compartments or compartment proportions [43]. We formalize recommendations for the integration of highly imbalanced datasets in Section VI.

### V. Balanced clustering metrics accurately benchmark imbalanced integration

Through extensive analysis of both simulated balanced and real-world imbalanced scRNA-seq datasets, we have shown that clustering metrics commonly used in quantifying scRNA-seq integration may be insufficient in imbalanced contexts. Metrics such as the AMI and ARI are agnostic to information on label proportions [26, 27] and are thus inadequate for assessing integration performance in imbalanced datasets, which is a common case in single-cell integration. To overcome limitations of routinely used metrics, we developed balanced versions of these scores, including the *Balanced Adjusted Rand Index* (bARI), *Balanced Adjusted Mutual Information* (bAMI), *Balanced Homogeneity*, and *Balanced Completeness*. Combining *Balanced Homogeneity* and *Balanced Completeness* also allows us to attain the *Balanced V-measure* [28]. These metrics are robust to dataset imbalance and allow for more nuanced comparisons of integration results in the aforementioned cases, as they weigh each cell-type present equally and are not driven by cell-types present in high proportions (Online Methods).

We first demonstrated the utility of the proposed balanced clustering metrics on simulated data. In the first scenario, we examined a dataset with 3 classes that are incorrectly clustered into 2 instances using K-means clustering [32] (Figure 6A) (Online Methods). This scenario can occur in single-cell settings when a cell-type is highly related to a neighboring cell-type and unsupervised clustering leads to a collapse of both into the same cluster. As expected, the base/imbalanced metrics all overestimated the clustering accuracy as they do not weigh the smaller class (B) as much as the larger classes when assessing the incorrect assignment (Figure 6A). However, the balanced metrics account for the incorrect clustering of class B, and indicated worse performance with the exception of the Balanced Completeness measure. This is an expected result as Completeness only measures whether all members of a given class are members of the same cluster [28].

**Figure 6:**
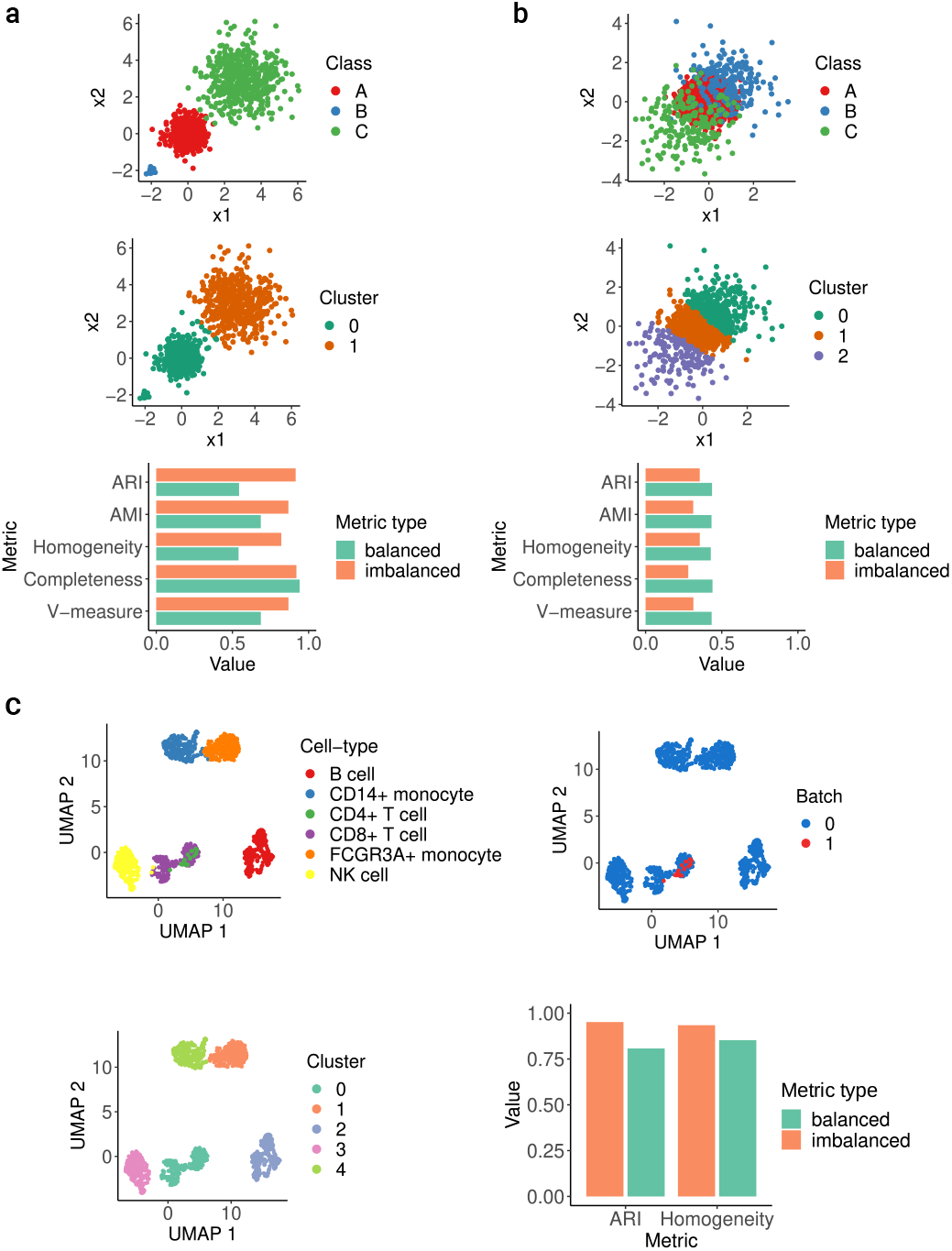
Demonstration of balanced clustering metrics on simulated data and scenarios. (a) Simulated data of 3 well separated imbalanced isotropic Gaussian classes with imbalance that are incorrectly clustered into two clusters that collapses the smaller class (B) with another. The concordance of the class labels with the clustering result for the base (imbalanced) and balanced ARI, AMI, Homogeneity, Completeness and V-measure for this result are indicated. (b) Simulated data of 3 imbalanced and overlapping isotropic Gaussian classes that are clustered into 3 clusters that mix the larger class (middle - A) with the smaller classes (B, C) and concordance of the class labels with the clustering result for the base (imbalanced) and balanced metrics. (c) Constructed scenario with balanced two batch PBMC single-cell data where a very small subset of CD4+ T cells (10% of original proportion) present in only one batch are incorrectly clustered with CD8+ T cells after integration. In this scenario, the concordance of the unsupervised clustering labels and the groundtruth cell-type labels are indicated for both the base (imbalanced) and balanced ARI and Homogeneity scores.

In a second scenario, we sought to check whether the balanced metrics can return higher performance than their base counterparts in the appropriate scenario, and to do so we simulated a dataset where a larger class (A) partially overlaps two smaller classes (B, C) (Figure 6B) (Online Methods). K-means clustering with a preset cluster number of 3 slices the larger class in a manner that the two smaller classes are mostly assigned to the correct cluster/label while the larger class is split between the 3 clusters (Figure 6B) (Online Methods). In this setting, the base metrics penalized the results based on the prevalence of the larger class (A) and the associated mis-clustering, but the balanced metrics took into account the strong performance on the smaller classes (B, C) and returned higher scores.

These two simulated scenarios demonstrate that balanced metrics can reveal information not present in typical global clustering scores and benchmark results in a manner that takes into account class imbalance.

We further assessed the applicability of the balanced metrics in single-cell data by considering the balanced PBMC cohort of two batches (Results I.). The first test assessed whether or not the first simulated case holds in single-cell data, as we downsampled CD4+ T cells in one batch in a manner where they overlapped with CD8+ T cells after clustering (Figure 6C) (Online Methods). Comparing the balanced and base ARI and Homogeneity scores, we found that the balanced scores did in fact decrease by a significant margin (Figure 6C). This is because the balanced metrics are considering the mis-clustering of the CD4+ T cells in a manner that is weighted equally to the correct clustering of the other cell-types, even though these cells are present in a much smaller number overall. To determine if the balanced metrics can change the results of a benchmarking analysis, we downsampled CD4+ T cells and FCGR3A+ monocytes from one batch in the balanced 2 batch PBMC dataset and performed integration using BBKNN, Harmony, Scanorama, and scVI (Figure 7A, Figure 7B) (Online Methods). After integration, Leiden graph-based unsupervised clustering [29] was performed and the results of clustering were compared with ground-truth labels using the base and balanced metrics by considering an average of their scores across ARI, AMI, Homogeneity, Completeness, and V-measure values (Figure 7C) (Online Methods). Examining the integration results, most methods mixed the CD4+ T cells with the CD8+ T Cells and the FCGR3A+ Monocytes with the CD14+ Monocytes to differing extents, while also having varying success in integrating the two batches overall (Figure 7A, Figure 7C). When using the base metrics and their averaged scores, scVI ranked the highest and BBKNN ranked the worst. Surprisingly, the base metric scores for Scanorama and BBKNN, which ranked the worst in this subset, were almost the same as using the unintegrated embedding (Figure 7D). Scanorama and BBKNN have shown strong performance with low variance results for all of our previous analyses and performed well in comprehensive benchmarking studies [9, 13], which is not in concordance with this result. When analyzing the result with the balanced metrics, the rankings changed significantly, as Harmony became the top performer (switched with scVI) and BBKNN now performed better than Scanorama (Figure 7E). Of particular note is the fact that there is a larger separation in scores between the unintegrated embedding and the results of BBKNN and Scanorama using the balanced metrics, and this result is more valid as we expect the integration methods to perform significantly better than an unintegrated baseline. This ranking shift occurred while the magnitude of the overall scores did not diverge significantly.

**Figure 7:**
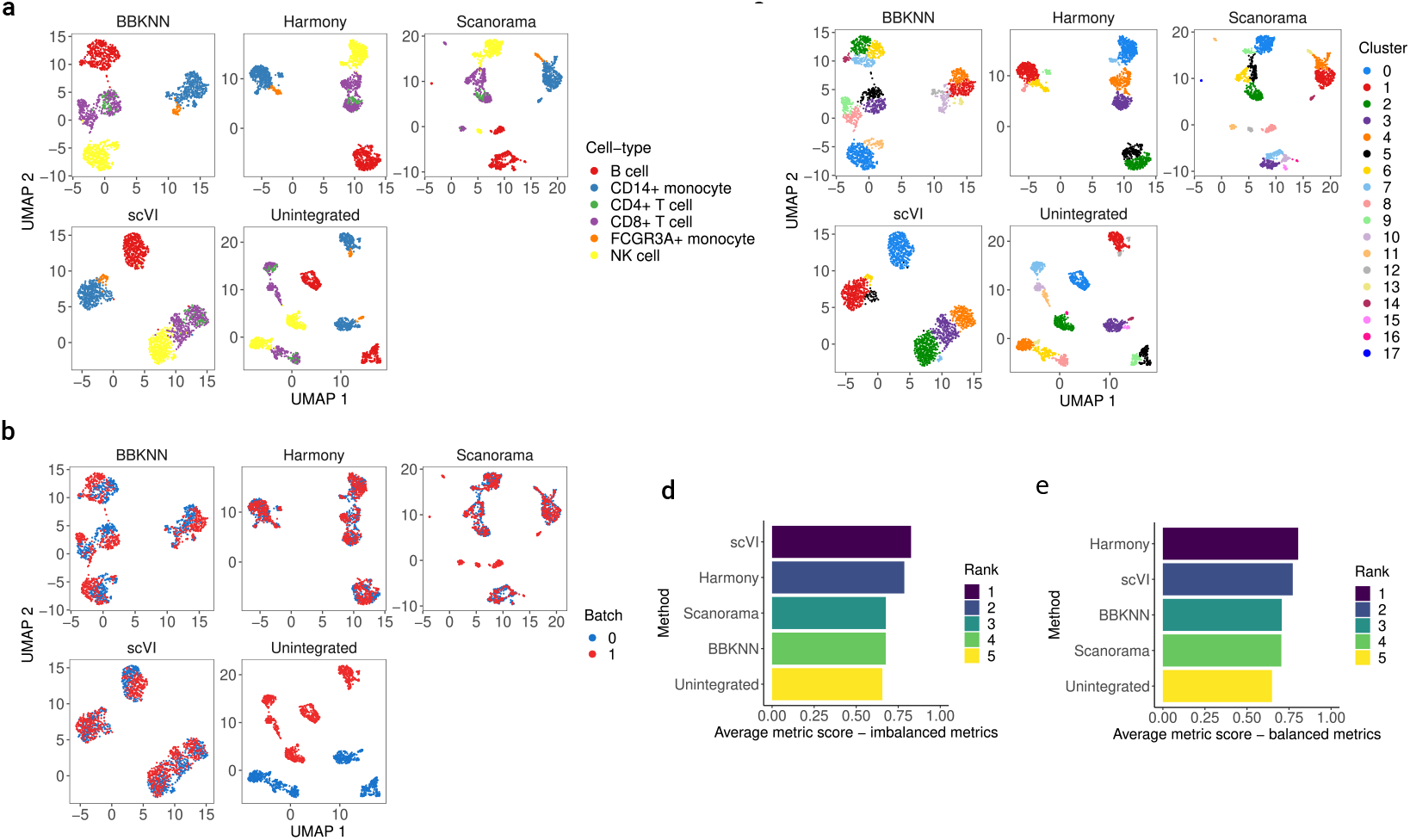
Benchmarking single-cell data integration using balanced clustering metrics. (a) Cell-type values for the balanced two batch PBMC data with FCG3RA+ Monocytes and CD4+ T cells downsampled to 10% of their original proportion in one batch, after integration with the tested methods as well as an unintegrated representation. (b) Batch values for the integrated and unintegrated downsampled two batch PBMC data. (c) Unsupervised clustering results for Leiden clustering in the embedding space of the integrated and unintegrated results for the downsampled two batch PBMC data. (d), (e) Scoring and ranking of integration results, when considering concordance of the unsupervised clustering labels and ground-truth cell-type labels for each integration method and the unintegrated subset, using the average results of the base (imbalanced) clustering metrics (d) (ARI, AMI, Completeness, Homogeneity) and average of the balanced clustering metrics (e) (bARI, bAMI, Balanced Completeness, Balanced Homogeneity).

Lastly, we reexamined the initial uninformative results obtained using the base ARI_*cell–type*_ scores for the perturbation experiments on the balanced 2 batch PBMC dataset (Figure 2D). Utilizing the balanced ARI (bARI) instead of the base metric for calculating ARI_*cell–type*_, the results indicated more clear/distinct patterns that reflected both the *relative cell-type support* and *minimum cell-type center distance* properties (Supplementary Figure S12). Specifically, downsampling/ablating B cells did not lead to decreases in the balanced ARI_*cell–type*_ across all methods, which is in line with the cell-type center distance property as the B cells are distant from all other cell-types in this dataset (Supplementary Figure S13). Further, there is a clear pattern of worsening performance in the balanced ARI_*cell–type*_ scores when the CD4+ or CD8+ T cells are downsampled or ablated, which is concordance with both the expected results in terms of *minimum cell-type center distance* (Supplementary Figure S13) and the KNN-classification results (Figure 2F). Similar results hold for downsampling or ablation of the monocyte subsets, although the scores are more method-specific and performance decreases are less pronounced (Supplementary Figure S12). Overall, the balanced clustering metrics can capture nuances in the data related to cell-type imbalance in a manner that the base metrics cannot. The balanced metrics are also more concordant with cell-type specific results, such as the KNN F1-score, as they weigh classes equally when considering performance.

### VI. Guidelines for imbalanced single-cell data integration

To aid in the integration of imbalanced datasets, we introduce general guidelines for users of integration techniques (Figure 8, Supplementary Table 2). We note that these guidelines are not meant to be strict rules, but rather suggestive in nature, as scRNA-seq and multi-modal single-cell data from different samples can have very different properties even after taking imbalance into account [4]. The guidelines are method agnostic, as our analysis revealed that all frequently utilized techniques in scRNA-seq integration are susceptible to the outlined effects of dataset imbalance (Results sections I-IV). An important aspect to consider when utilizing these guidelines is prior knowledge on potential disparity in the datasets can help guide the degree of desired batch mixing. For instance, in analyzing heterogeneous tumor samples from distinct patients with disparate cell-types and proportions, biological heterogeneity conservation is likely to be poor if batch-mixing is prioritized in integration [8]. However, this may be a desired result if the end analysis goal is only to assess common variation between the tumor samples and perform downstream analyses such as differential abundance of shared cell-types [8]. Judging the degree of desired batch mixing is often very difficult in practice [8]. As such, we emphasize an iterative process where imbalance, degree of batch-correction, and conservation of biological heterogeneity are assessed at multiple steps in the scRNA-seq integration pipeline (Figure 8).

**Figure 8:**
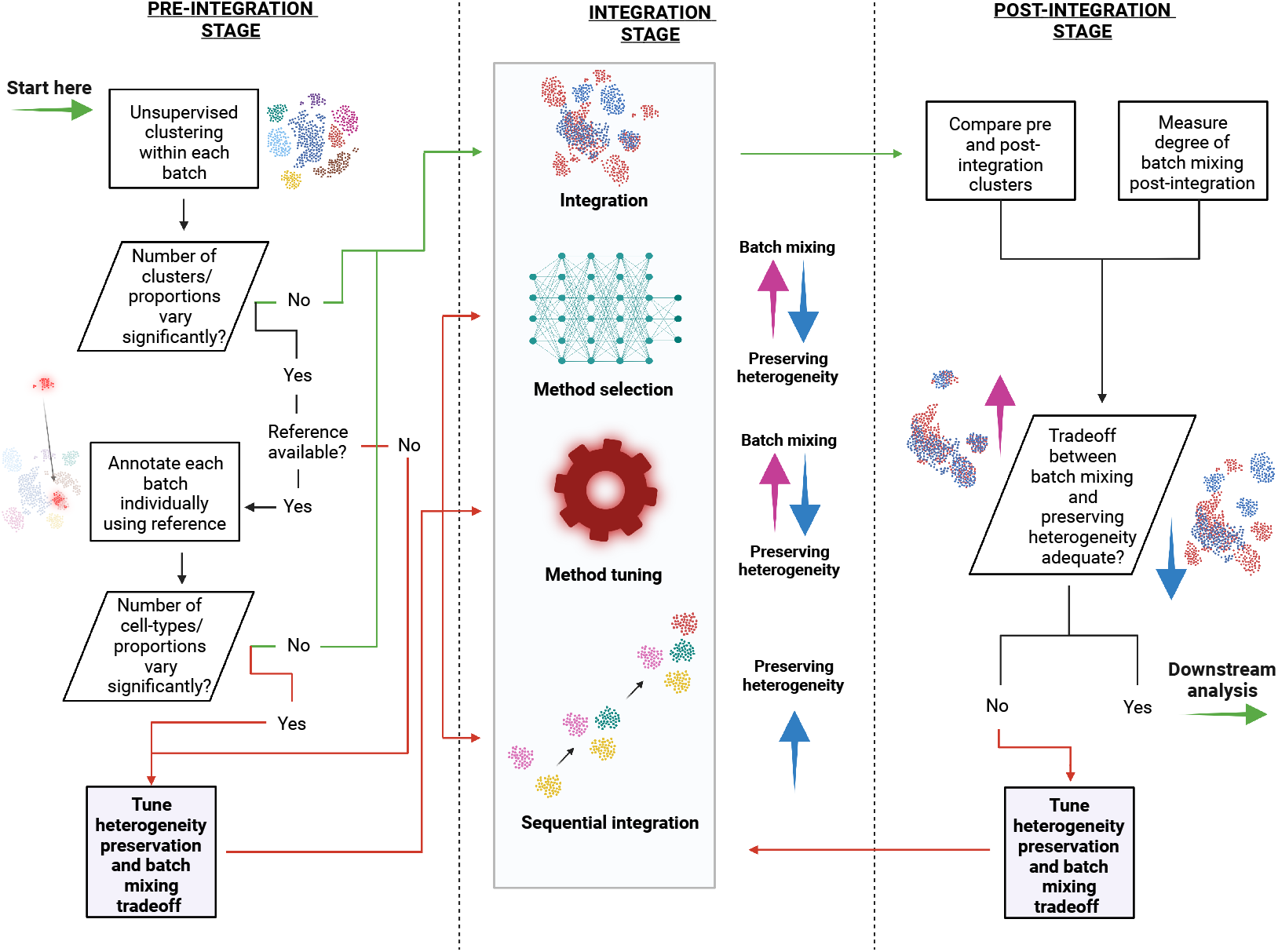
Guidelines for single-cell integration in imbalanced settings. A stepwise procedure is outlined, starting with diagnostic tests in the pre-integration stage that dictate whether or not to tune integration methods at the integration stage or perform further steps in the pre-integration stage. After integration, the trade off between batch-mixing and conservation of biological heterogeneity can also be diagnosed, and if determined inadequate, further tuning at the pre-integration and integration stages can be done. Complete details as well as examples of implementations for each recommendation are given in Supplementary Table 2.

Overall, potential imbalance within datasets to be integrated can be assessed based on pre-integration tests using unsupervised clustering and/or query-to-reference annotation (Figure 8, Supplementary Table 2). The latter would yield a more accurate representation of potential imbalance, but can only be used when a reference dataset is available for the given tissue samples. Unsupervised clustering can be used in any situation as it does not require cell-type labels but can be a noisy readout as clustering is highly sensitive to the technique, parameters, and the underlying data distribution [44]. If either of these outlined pre-integration tests reveals disparity in the datasets, the integration step itself can be altered by: (i) picking an integration method that is suitable for preserving biological heterogeneity - Luecken et al. [13] provide an extensive overview of scRNA-seq and multi-modal integration techniques in this respect and provide selection criteria, (ii) tuning the integration method itself to better preserve biological heterogeneity across datasets - the availability of such parameters will vary based on method [13], and (iii) performing sequential integration if the datasets are known or suspected to have temporal structure [45] (e.g. developmental data) (Figure 8). There is also the possibility of integrating only shared putative cell-types between batches if a reference dataset is available, as this would better ensure imbalance is minimized in the integration step (Supplementary Table 2). After the integration step, post-integration techniques to assess preservation of biological heterogeneity and degree of batch mixing can be used to determine the current balance between the two desired outcomes [8], and integration and pre-integration steps can be further tuned to strike the desired balance (Figure 8). A complete description with specifics and code implementations, in the R and python programming languages, of the outlined recommendations are indicated in Supplementary Table 2.

## Discussion

In this work, we thoroughly analyzed the effects of dataset imbalance in scRNA-seq integration scenarios, and its impacts on downstream analyses and overall biological conclusions. When the level of imbalance between batches was perturbed, we observed varying degrees of effects on unsupervised clustering, neighbor-based cell-type annotation, differential gene expression analysis, and query-to-reference annotation. More importantly, these effects were not method-specific, and thus have implications for single-cell data integration overall, where biological conclusions plausible under one scenario may not be concordant if the pre-integration data distribution is different due to many possible underlying factors of variation. These results have significant ramifications for single-cell data integration, as most datasets being integrated will likely not have a high degree of shared variation with the increasing complexity of the tissues being analyzed and higher throughput of current scRNA-seq and multi-modal sequencing protocols [37, 46]. We further examined these results on more complex data and concluded that the potential of dataset imbalance to affect integration results can be summarized by two key metrics - *relative cell-type support* and *minimum cell-type center distance*. To aid the integration and subsequent downstream analyses in scenarios with imbalanced datasets, we introduce several guidelines pre-integration, at the integration step, and post-integration, as well as balanced clustering metrics for more accurate assessment and bench-marking in such cases.

Although single-cell data integration is ubiquitous in current computational analysis pipelines for both scRNA-seq and multi-modal single-cell sequencing data, analyzing the nuanced properties and behavior of integration techniques on different datasets has lagged in lieu of performance-based studies. Extensive benchmarking studies have been performed for scRNA-seq integration, but these analyses have largely focused on performance in specific settings, as determined by batch-mixing and conservation of biological heterogeneity [9, 47]. Some studies have raised specific concerns towards the impacts dataset imbalance can have on integration and a few methods have been developed specifically to address this challenge [10, 36, 48–50], but an extensive analysis on downstream effects had yet to be performed. Further understanding of the properties of both the pre-integration and post-integration representation spaces will likely shed light on gaps in performance between different techniques and the situational trade offs between batch mixing and conservation of biological heterogeneity. For example, although anchor-based techniques are used to link both scRNA-seq and multi-modal datasets in integration [45], the conditions that lead to false-positive and false-negative (missing) anchors between batches and multi-modal samples have not been extensively characterized. Such analyses will further the understanding of the limiting conditions of single-cell data integration, and lead to better tools, guidelines, and a sounder foundation for downstream analyses and inference of biological phenomena.

In benchmarking single-cell integration techniques, often standardized datasets do not contain high degrees of cell-type imbalance across batches that may be encountered in common real-world scenarios such as temporal integration [9, 13, 45]. Therefore, a more principled approach to benchmarking may involve a stronger focus on these cases and non-trivial datasets such as tumor samples from multiple-patients and cohorts. The trade-off between batch mixing and biological heterogeneity conservation is an important research direction, as conserving biological signal can be much more complex than what is indicated by clustering metrics post-integration, particularly if integration is being done on the entire count matrix and not within an embedding space [13]. In our analysis, we introduced four novel balanced clustering metrics that can be utilized to better benchmark integration techniques in imbalanced scenarios. These metrics are used to analyze clustering results post-integration, but more salient scores for preserving biological signal such as the conservation of highly-variable genes introduced by Lueken et al. [13] will also allow for a complete picture of the potential downstream impacts of integration. As our understanding of the limitations of current integration methods evolves, we envision more comprehensive guidelines that incorporate our analysis on situational integration setups and utilization of different methods in the correct contexts, as opposed to a single method for every integration scenario. Our findings and guidelines can be extended to multi-modal analysis of disjoint samples (e.g. scRNA-seq and scATAC-seq of similar but distinct tissue samples), but the finer details of the impacts of imbalance on both joint and separately profiled multi-modal integration and subsequent analysis remains unknown. An important future research direction in multi-modal integration is better understanding of integration results at both the technique and data level, as comprehensive benchmarks specifically focused on multi-modal data have yet to be completed.

Our analysis is limited by the extent of datasets analyzed and methods tested. We sought to identify the effects of downsampling in a highly controlled scenario where imbalance was not already present, which was the balanced PBMC 2 batch dataset, but extrapolating the results of the downsampling experiments to more complex cases was not straightforward. Thus, only the cell-type-specific effects in already imbalanced datasets with no perturbations were examined for complex cases, although we did analyze perturbation of complex and imbalanced PDAC samples. Further, although we included frequently utilized and best performing scRNA-seq integration techniques based on previous benchmarking studies [9, 13], we did not include recent methods that focus specifically on preserving biological heterogeneity when differing cell populations are present between samples, such as CIDER [49]. There have also been strides in this direction in the multi-omic integration space, with techniques such as SCOTv2 [50]. As the aim of this analysis was not to determine the best performing method, but to analyze the impacts of imbalance on integration results with frequently utilized methods, we deemed this omission to be acceptable. However, future method-based benchmarking studies should feature techniques that have sought to explicitly address the issue of dataset imbalance and several datasets with a high degree of imbalance present. Lastly, this analysis focused on scRNA-seq integration and did not incorporate multi-modal datasets and techniques, and although extrapolation may be possible, this must be confirmed by future work addressing integration when imbalance across jointly and separately profiled multi-modal datasets is present.

## Online Methods

### 1 Dataset preprocessing

#### 1.1 Preprocessing and normalization

All datasets utilized in the study were preprocessed using a uniform pipeline. Datasets were only further processed if it was clear that no filtering was done on the raw scRNA-seq data, such as removal of low quality cells and genes [8]. If it was indicated that no filtering was done, quality-control (QC) metrics were calculated using the *Scuttle* R package (version 1.4.0) [51], including cells with the most genes having low counts, cells with a high percentage of mitochondrial genome content, and cells with a low library size (total number of reads overall). An approach recommended by Amezquita et al. [8] was taken, and the cells and features with values more than 3 median absolute deviations (MADs) for two out of three criteria were filtered out. As normalization and log-transformation need to be tuned specific to the method being utilized and are done in the integration pipeline where necessary, these were not done in the preprocessing steps.

Datasets were split and saved as individual batches in *h5ad* format, and the *scanpy* library (version 1.8.2) [52] was used for all further downstream processing within the integration pipeline, including total-count per cell normalization, log1p transformation, and highly variable gene selection [52]. These steps were carried out uniformly for each method tested, with the exception of scVI, as the technique must utilize the raw counts [24]. Therefore, total-count normalization and log1p transformation were not done for scVI.

All scRNA-seq datasets utilized were preprocessed in this manner, including the balanced 2 batch PBMC dataset [15][14][9], imbalanced 2 batch PBMC dataset [18], 4 batch PBMC dataset [18], 6 batch mouse hindbrain development dataset [19] and 8 batch pancreatic ductal adenocarcinoma (PDAC) dataset [20].

The **ground-truth annotations for cell-types** in each dataset across batches were determined specific to the annotation protocol followed by each original study, with the exception of the 8 batch PDAC data, which was re-annotated (see 1.3).

#### 1.2 Setting up the PBMC control dataset

For testing in a scenario where the cell-types and cell-type proportions are perfectly balanced between batches, and subsequent perturbation experiments, a dataset that was preprocessed by Tran et al.[9] comprising of two batches of peripheral blood mononuclear cells (PBMCs) sequenced using two variants of 10x genomics protocols - 5’ versus 3’ end. As these technologies capture different regions of mRNA, there is an expected batch effect present. To create a balanced dataset, the two batches were downsampled for cell-types that had at least 200 cells in each batch - leaving B cells, CD14+ Monocytes, CD4+ T cells, CD8+ T cells, FCGR3A+ Monocytes, and Natural Killer (NK) cells. Within each batch, these remaining cell-types were randomly downsampled to 200 cells, leading to a perfectly balanced control setup for perturbation experiments.

#### 1.3 Setting up the pancreatic ductal adenocarcinoma (PDAC) dataset

The pancreatic ductal adenocarcinoma dataset was taken from the Peng et al. [20] multi-patient study, which comprised of 23 samples. For this data, custom annotations of tumor cells were done in the following manner:

Cells from different samples were integrated with Harmony [22] and clustered with Seurat [25]. A cell type label was then assigned to each Seurat cluster, based on the expression of specific marker genes for each cell type (Supplementary Table 8). To identify tumour cells, all epithelial cells including those from normal tissue were clustered again using Seurat. All cells that clustered with the normal samples were assigned as ’Epithelial normal’, while all others were assigned as ’Epithelial tumor’.

After annotation of the epithelial normal and tumor cells, the rest of the cells were collapsed into the ’Microenvironment’ compartment. Ductal and acinar cells that did not fall into the epithelial normal or epithelial tumor populations were removed, as these were likely mis-annotated. Batches/samples were filtered based on the presence of at least 50 cells in each of the three compartments (Epithelial normal, Epithelial tumor, Microenvironment). This left 8 batches/samples, which were utilized in subsequent experiments.

### 2 scRNA-seq integration methods and parameters

Five state of the art scRNA-seq methods were utilized, based on their performance in previous benchmarking papers [9] [13], including BBKNN (version 1.5.1) [21], Harmony (python implementation - version 0.0.5) [22], scVI (scvi-tools version 0.14.4) [24], Scanorama (version 1.7.1) [23], and Seurat (version 4.0.6) [25]. LIGER (version 0.5.0) [53] was also originally tested, but did not indicate strong performance and resulted in a high degree of variability due to the removal of seeding in different steps. Therefore, the results from LIGER were omitted from the main findings, as the high variance of results even within the control experiments did not allow for a statistically sound comparison between control and perturbation groups.

Because the perturbation experiments were carried out in replicates, to get a more clear sense of variability within replicates, seeding mechanisms within each method were removed. This included removing any calls in the method source-code to R-based seeding for Seurat, and any calls to seeding from the following libraries for BBKNN, Harmony, Scanorama, and scVI: random, numpy, torch. This led to a more true estimation of the variability in performance of each method, as well as a more reliable estimation of the effects of perturbation because the variability can no longer be simply attributed to variability in the method which has been accounted for.

With the exception of scVI, each method utilized the same processing pipeline for the data, where scanpy’s functions [52] were utilized in the following manner:

1. Count normalization for each cell to total value of 1 * 10^4^
2. Transformation of counts using the log(1 + *x*) function
3. Highly variable gene selection using the ’seurat’ method for 2500 genes
4. **Integration at this step for Seurat - returns corrected HVG counts**
5. Principal component analysis (PCA) reduction to top 50 principal components that explain the highest variance (PCs) for HVG counts
6. **Integration at this step for Harmony, Scanorama - return corrected PCs**
7. Creating neighborhood graph using the embedding with 20 dimensions (highest explained variance) and 15 nearest-neighbors - **integration at this step for BBKNN (replaces neighborhood graph step in scanpy pipeline)**
8. Leiden clustering on the neighborhood graph using scanpy’s default parameters
9. Uniform Manifold Approximation and Projection (UMAP) on the neighborhood graph using scanpy’s default parameters

The only exception to this setup was scVI, which requires raw scRNA-seq expression counts [24], and utilized the entire set of genes for each dataset and the raw counts. For scVI, steps 1-6 outlined are omitted, and it simply integrates the raw data and returns a 10 dimensional embedding, which replaces the reduced dimensions of the other methods (**input embedding for step 7**). 10 dimensions were utilized in this case instead of 20, as this was the indicated default setting for scVI. The rest of the steps (7-9) are the same.

BBKNN performs integration on the embedding neighborhood representation [21], and as a result, many of the downstream analyses that required embeddings did not have data for BBKNN as it was untestable. Default parameters were utilized for all methods to ensure fairness in across-method comparisons, as well as comparisons before and after perturbations.

### 3 Perturbation experiments

Perturbation experiments were carried out in two settings - the balanced 2 batch PBMC data, and the pancreatic ductal adenocarcinoma data. In both instances, batches are randomly selected to be perturbed, as well as given cell-types/compartments. There are three types of perturbation experiments performed - control, downsampling, and ablation. Control experiments don’t downsample any data but allow for replicates of integration runs across methods to get a sense of intra and inter-method variance on the data without perturbation. Downsampling experiments involve randomly selecting cells of a selected cell-type across the indicated number of batches, and downsampling to 10% of the original cell-type population. Ablation experiments involve completely removing selected cell-types from the indicated number of batches. Randomness of selection for the batches, cell-types, and cells within indicated cell-type are ensured through randomly generated numbers for each perturbation simulation/run. To determine the effects of perturbation, results from the control experiments are compared with results from downsampling and ablation experiments, across all methods and selected datasets. The code for the experimental setup, as well as the Iniquitate pipeline, are available at https://github.com/hsmaan/Iniquitate.

#### 3.1 Balanced 2 batch PBMC data

Within the balanced 2 batch PBMC dataset, perturbation experiments were performed for one of two batches (randomly selected) at a time, and for one cell-type at a time. 400 replicates were done for the control experiments, and 200 replicates were done for the downsampling and ablation experiments, ensuring that both batches (n=2) and each cell-type (n=6) is sampled repeatedly and method performance variance within control experiments is taken into account adequately. Within the **hierarchical setup**, where similar cell-types were hierarchically clustered into 3 groups (B cell, Monocyte, NK/T cell), the same number of replicates were done for for the control, downsampling and ablation experiments.

#### 3.2 8 batch PDAC data

For the 8 batch pancreatic ductal adenocarcinoma dataset, where cells were grouped into three major compartments, 4 batches were randomly selected for downsampling or ablation, and one compartment was downsampled or ablated within one replicate. In total, 100 control replicates were performed, and 50 downsampling and ablation replicates, as the number of compartments is small (n=3) and there will be adequate sampling and repetition within 50 runs.

### 4 Benchmarking integration performance - PBMC 2 batch control dataset

Performance in integration and downstream tasks was assessed using the integrated embeddings and Leiden clustering [29] results from each integration technique. After integration at either the embedding or neighborhood calculation stage through the scanpy library, Leiden clustering was used. Default values were used for the embedding and clustering steps in the scanpy library [52]. Only BBKNN did not result in embeddings to be utilized as it performs integration at the neighborhood, and therefore was not included in the *K-nearest neighbors classification* experiments as these relied on integrated embeddings.

#### 4.1 Quantifying cell-type conservation and batch-mixing with clustering metrics

Four metrics were calculated for all integration experiments, including perturbations and replicates, - Adjusted Rand Index (ARI) [26], Adjusted Mutual Information (AMI) [27], Completeness [28], and Homogeneity [28]. The *sklearn* (version ≥ 1.0.1) implementation of these metrics was utilized. Details of these metrics can be found in the scikit learn documentation [32]. These values were calculated by comparing the known annotated labels with the cluster labels obtained after integration for each technique. Both cell-type and (1 - batch) values were calculated for each metric, where cell-type metrics compared the known cell-type annotations with the cluster labels to determine how well the integrated embeddings corresponded to known cell-type labels, and the (1 - batch) values used the batch annotations and the cluster labels to determine how well the different batches co-aggregate in the embeddings. The assumption of the latter is that integration should lead to strong batch mixing, and the shadow of the value is used (1 - batch) to reflect this desired property. The **median** value across all replicates for a given combination of {method, experiment type, downsampled cell-type} was determined. For the main analysis, the cell-type and (1 - ARI_*batch*_) was utilized, but values for all metrics were calculated (Supplementary table 9).

##### 4.1.1 Z-score normalization of ARI metrics

As the focus of the analysis was not to assess inter-method variation, but determine intra-method variation based on the properties of perturbations versus the control experiments, the median values for cell-type and (1 - ARI_*batch*_), for all combinations of {method, experiment type, downsampled cell-type}, were Z-score normalized. E.g. for cell-type ARI values for a specific subset:

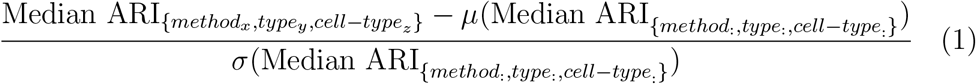

The exact same procedure is followed for the (1 - ARI_*batch*_) values.

#### 4.2 Downstream analysis - unsupervised clustering

In this downstream analysis test post-integration, the number of unsupervised Leiden clusters are determined and compared between the different perturbation experiments and control groups. The same indicated setup is used, and the number of clusters are determined using the default parameters for Leiden clustering in the scanpy library [52]. As each method resulted in different Leiden clusters, these were analyzed independently and intra-method and experiment-type comparisons were performed.

#### 4.3 Downstream analysis - k-nearest neighbor (KNN) classification

The goal of this downstream analysis test was to determine the performance of integration techniques at a per-cell-type level before and after perturbation. After obtaining the integrated embeddings for all methods, with the exception of BBKNN, a KNN classifier is trained on a 70/30 training/test split of the integrated embeddings to predict the cell-type labels of the test data. Stratified sampling was used for the split to ensure that all classes were represented in the same proportions between train and test sets. The *sklearn* (version ≥ 1.0.1) library was used for the data preparation, test/train split, stratified sampling, KNN-classifier training and prediction [32]. The explicit formulation for prediction of a class on a test data point *x_i_* is:

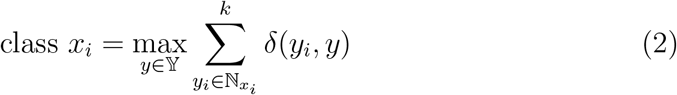

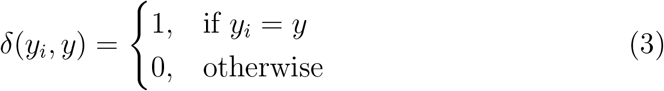

Where *k* indicates the number of neighbors used in the classifier, which was set to 15 for all runs. As different runs could possible lead to different test/train splits of the integrated embedding, a seed was used to ensure that the same split occurs across all experiments. This also ensured that each method was tested on the same split of the data. Using the results of the predictions, the cell-type-specific precision, recall, and F1 scores were determined, and the F1-score specific to each cell-type was used as key metric. These metrics was calculated Primarily, cases were examined where a specific cell-type was downsampled or ablated, and the effects of performance on the same cell-type based on the KNN-classification F1-score was analyzed. The form of the score is given by [33]:

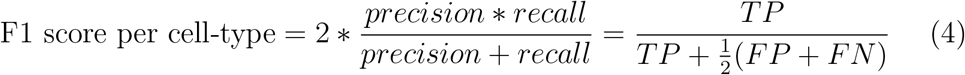

Where *TP* is the number of true positive calls, *FP* is the number of false positive calls, and *FN* is the number of false negative calls, all on a per-cell-type basis.

#### 4.4 Downstream analysis - marker gene ranking

Differential gene expression (DGE) analysis is typically performed after clustering (or clustering an integrated representation of many batches/samples) to determine marker genes specific to each cluster that are then used to annotate cells within those clusters [8]. The goal of this analysis was to determine to what extent can the results of DGE be altered after perturbation of balanced data.

First, the top 10 marker genes corresponding to each cell-type were determined in each batch for each dataset (e.g. 2 batch PBMC dataset) using the Wilcoxon Rank-Sum Test and the scanpy package (sc.tl.rank_genes_groups) [52]. To ensure selection of relevant markers, ribosomal and mitochondrial genes were removed from the pool of tested genes. After this, to obtain a consensus on the marker genes across batches, the union of markers for each cell-type across batches was determined and duplicate gene calls across batches were dropped. This will lead to an uneven number of markers for some cell-types if completely distinct sets are called from different batches, but leads to a more complete set as the integrated space across batches is being analyzed. This set of markers for each cell-type in a dataset was deemed the **master marker list**.

From here, after each the integration step using each method in each control or perturbation simulation for a given dataset, unsupervised clustering using the Leiden clustering algorithm with the scanpy default parameters was done for the integrated embedding and DGE using the Wilcoxon Rank-Sum Test was performed for each of the obtained unsupervised clusters [52]. A challenge here is that *there is no correspondence between the unsupervised clusters obtained in this integrated embedding and the cell-types used for determining the master marker list*. However, a way to get around this is to do DGE for each cluster and check the **maximum ranking** of a given marker gene across all clusters. Ranking is defined by how significant the DGE *p*-value is for a given gene, where the highest rank is the most statistically significant differentially expressed gene for a given cluster. If a cluster still corresponds to a given cell-type (which is the central assumption in unsupervised integration), then that cluster should return a high ranking for a given marker gene corresponding to that cell-type in DGE. Therefore, for the markers in the master marker list, we can analyze their **maximum ranking** across unsupervised clusters in the integrated space - to see if biological information specific to that cell-type and its markers is still being retained after integration.

This is precisely the operation carried out, and the change in ranking for all of the marker genes corresponding to the different known cell-types in datasets quantified by their standard deviation in a given subset of experiments (control, downsampling, or ablation) **change in maximum ranking**. This change in maximum ranking within an experiment group (e.g. the control group) was indicated as the **marker gene perturbation score**:

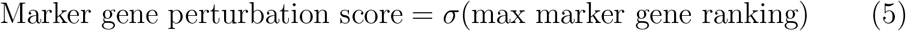

If this value is being averaged over many genes (e.g. for a cell-type), this is indicated as the **average marker gene perturbation score**. If there are *m* marker genes for a given cell-type:

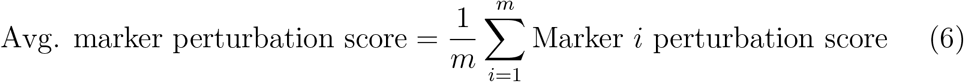

##### 4.4.1 Case study - CD4/CD8 T cell assignment based on marker genes

To determine if the changes in marker gene ranking that were observed could realistically influence the results of a single-cell analysis, the same marker gene perturbation set-up was utilized. In this case however, each of the unsupervised clusters after integration were annotated as specific cell-types based on a majority of their cells present (e.g. Cluster 1 -¿ Majority CD4+ T cells -¿ CD4+ T). In this setup, only clusters that contained a majority of either CD4+ or CD8+ T cells were kept. For simplifying the case study, only integration results from the Seurat method were utilized.

After integration and selection of clusters with a majority of CD4+ and CD8+ T cells, differential expression analysis was performed as previously indicated, and a permissive threshold of the top 50 marker genes was used to select markers for the CD4+ and CD8+ majority clusters. From here, each of the CD4+ and CD8+ T cell majority clusters were predicted to be either CD4+ or CD8+ based on the presence of canonical marker genes: IL7R for CD4+ T cells and CD8A for CD8+ T cells [25].

Examining the top 50 marker genes for each cluster, the rules for predicting the cell-types each of the CD4+ and CD8+ T majority clusters comprised of were the following:

~~~
  **if** IL7R **and** CD8A present **then**
     **if** Rank(IL7R) > Rank (CD8A) **then**
        Annotate as CD4+ T
     **else if** Rank(CD8A) > Rank (IL7R) **then**
         Annotate as CD8+ T
     **end if**
  **else if** IL7R present **then**
     Annotate as CD4+ T
  **else if** CD8A present **then**
     Annotate as CD8+ T
  **else**
     Annotate as Undefined
  **end if**
~~~

From here, the fraction of unsupervised clusters that contained a majority of CD4+ and CD8+ T cells were predicted for their cell-types based on differential expression in control and perturbation (downsampling and ablation) experiments, including replicates. Only downsampling and ablation experiments that affected CD4+ and CD8+ T cells were analyzed, as downsampling/ablating these were found to most likely affect the marker gene rankings of either cell-type.

#### 4.5 Downstream analysis - query-to-reference annotation

To test the robustness of query-to-reference annotation techniques across varying degrees of unshared variation, the Seurat 4.0 multi-modal projection technique was utilized [31]. Although the control PBMC 2 batch dataset has only scRNA-seq information, a multi-modal reference can still be utilized as is, as only the RNA-seq modality will be integrated. The reference dataset utilized is from Hao et al., and the same parameters indicated in the vignette were utilized [31].

It’s important to note that **integration was not performed before projection** using the Seurat 4.0 method. Instead, **each batch/sample is individually projected/integrated to the reference dataset and annotated**, as per the guidelines for Seurat 4.0 [31]. Therefore, there are no method-specific comparisons to be made in this analysis.

As the annotations in the reference will not exactly match the annotations from the PBMC 2 batch data (mostly due to a higher degree of granularity and different naming conventions) [9] [31], a scoring guide was created to determine if the annotation correctly matches the ground-truth cell-type label by using ”fuzzy-matching” of ground-truth cell-type labels from the PBMC 2 batch dataset and the labels in the reference data. The following table summarizes the guide for the PBMC 2 batch data, and acceptable annotations for L1 (coarse-grained label from Hao et al. [31]) and L2 (fine-grained label from Hao et al. [31]):

| Ground-truth label | Acceptable L1 reference | Acceptable L2 reference |
| --- | --- | --- |
| CD4 T cell | CD4 T | CD4 TCM, CD4 Naive, CD4 CTL, CD4 Proliferating, CD4 TEM |
| CD8 T cell | CD8 T | CD8 Naive, CD8 TEM, CD8 TCM, CD8 Proliferating |
| Monocyte_CD14 | Mono | CD14 Mono |
| Monocyte_FCGR3A | Mono | CD16 Mono |
| NK cell | NK | NK, NK Proliferating, NK_CD56bright |

Using this annotation guide, the annotation accuracy as determined by the F1 score (4.3) was determined for each experiment and experiment type (control, downsampling, ablation). This value was calculated using both the L1 and L2 annotations, and the annotations for each cell in each experiment were saved.

### 5 Complex imbalanced dataset analysis

After quantifying the effects of unshared variation in the control 2 batch PBMC dataset through perturbation experiments, complex datasets that are multi-batch and already imbalanced were analyzed, including: imbalanced 2 batch PBMC dataset, batch PBMC dataset, 6 batch mouse hindbrain development dataset, and 8 batch pancreatic ductal adenocarcinoma dataset.

#### 5.1 Cell-type center distance

To determine the distance between cell-types in the embedding space utilized for integration, across all batches to be integrated, the following preprocessing steps were performed on the raw data for each batch in a dataset:

1. Count normalization for each cell to total value of 1 × 10^4^
2. Transformation of counts using the log(1 + *x*) function
3. Highly variable gene selection using the ’seurat’ method for 2500 genes
4. PCA to top 20 (PCs) for on the counts data

After obtaining the PCs for a given dataset, the ground-truth cell-type labels are used to determine the cell-type center distance between all cell-types in the data in a pairwise manner. The cell-type center distance is defined as the weighted cosine distance between the center (average) of the PCA representation for each cell-type in a given batch.

For each batch *b*, and cell-type *a* with *n* cells and a PCA reduction of the data:

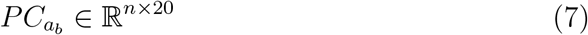

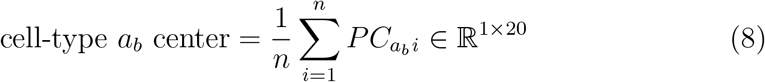

Then for quantifying the distance between cell-types *a* and *c* in batch *b*:

Let 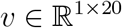 be the variance explained by each of the top 20 PCs

Let *CC_a_b__* be the cell-type center for cell-type *a* in batch *b*
Let *CC_c_b__* be the cell-type center for cell-type *c* in batch *b*

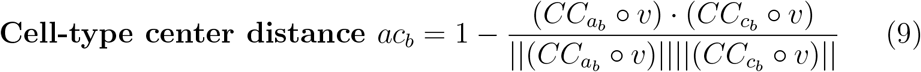

Where *CC_ab_* ∘ *v* is the element-wise rescaling of the cell-type center of *a* based on the variance explained by the PCs.

The rationale behind a reweighted cosine distance is that the distance itself between cell-types should be scaled according to the variance explained by each PC because the distance is being calculated in the joint PCA reduction of all cells, and not every PC axis will have equal contribution for the variance explained.

We can take the average of this cell-type center distance across *p* batches:

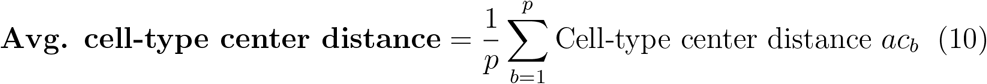

From here, the minimum cell-type center distance, or the distance corresponding to the cell-type closest to cell-type *a* is simply the minimum value across all batches (*p* total). Assume there are *k* total cell-types across all batches and missing cell-type pairs (e.g. cell-type present in batch 1 and not batch 2) have an imputed maximum cosine distance of 1. Values between the same cell-types are also imputed as 1. Then using the tensor of cell-type center distances across batches *D:*

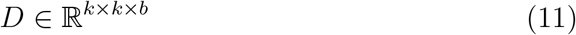

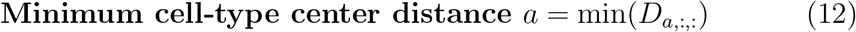

Where *D_a_* is the subset of the first axis for cell-type *a*. The cell-type and batch corresponding to this value can also be found through the *argmin*.

The minimum distance in any batch is taken instead of averaging distances across all batches because this minimum distance will correspond to the most ’haphazard’ scenario for a given batch being integrated. There are two scenarios possible here:

1. The similarity between cell-types is largely similar across batches, and the minimum value will correspond roughly to the average
2. The similarity between cell-types can be very different across batches, due to scenarios/factors such as developmental data or treatment-effects

The first case is most readily applicable to the PBMC datasets, but the second scenario may be more applicable to the PDAC and hindbrain developmental data. However, even in these cases, taking the minimum may lead to a better approximation of proximity affecting integration results because it will factor in the worst possible scenario (across batches) for a given cell-type.

#### 5.2 Cell-type support

The cell-type support (or relative cell-type support) was simply the log2-transformation of the number of cells for each cell-type *a* across all batches *b*:

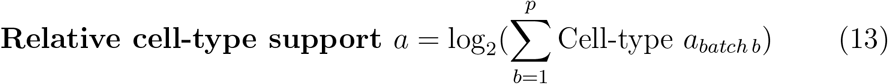

### 6 Statistical testing

#### 6.1 One-way ANOVA tests

To determine statistical significance for the effects of perturbations, the following generic one-way analysis-of-variance (ANOVA) setup was utilized [34]:

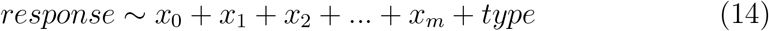

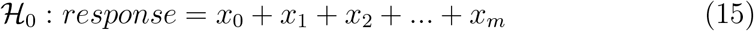

Where 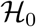 is the null hypothesis, *response* can be an endpoint of interest in the analysis (e.g. number of clusters post integration), *x*_0_ is a constant (intercept/bias), *x*_1_, …, *x_m_* are factors we’d like to control before testing significant with respect to perturbations (e.g. method, cell-type that was downsampled), and *type* is a binary covariate indicating the experiment type that was done:

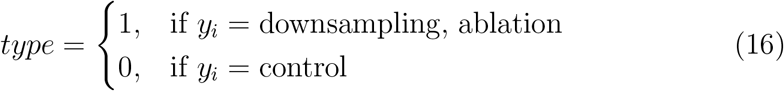

After accounting for the various factors we’d like to control (*x*_1_, …, *x_m_*), we can assess the statistical significance of perturbation of unshared variation (*type*) with respect to the *response* covariate through the **ANOVA F-statistic** and *p*-**value** associated with the *type* covariate.

In situations where significance is achieved across various groups due to factors such as intra- and inter-method variance, the magnitude of the F-statistic is compared.

#### 6.2 Control PBMC 2 batch dataset

##### 6.2.1 KNN classification per cell-type

For assessing the effects of perturbation on the F1 classification scores post-integration on a per-cell-type level, the following ANOVA (6.1) setup was utilized:

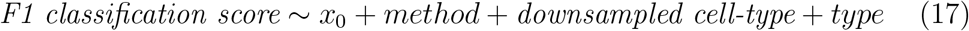

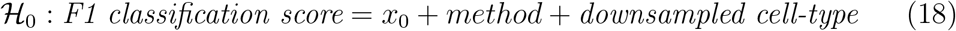

The F1-classification scores here are across all cell-types in the integrated dataset. **The cell-type being analyzed (for the F1 classification score in each instance) is equivalent to the downsampled cell-type in each sample included in the test.**

##### 6.2.2 Unsupervised clustering

For comparing the significance of perturbation on the number of unsupervised clusters obtained post-integration using Leiden clustering, the following ANOVA (6.1) setup was utilized:

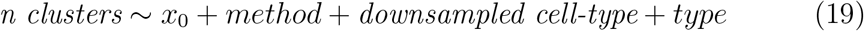

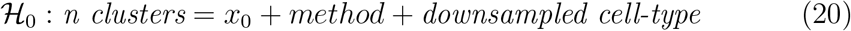

##### 6.2.3 marker gene ranking

To test the statistical significance of perturbations for each marker gene analyzed (4.4), the following ANOVA setup was used for each marker gene *g*:

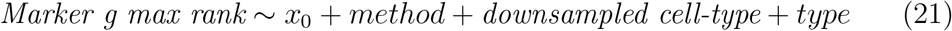

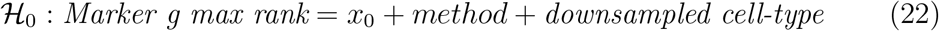

Then, to test the overall effects on marker gene ranking, considering all marker genes at once, the following test was done:

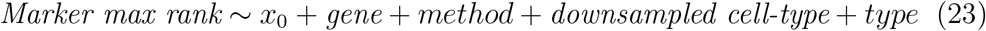

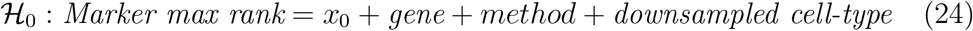

#### 6.3 Complex imbalanced datasets

##### 6.3.1 Cell-type support and cell-type center distance

To determine if the two key metrics that were determined in the complex dataset analysis - **relative cell-type support** (5.2) and **cell-type center distance** (5.1) - are in fact predictive of integration performance, the following ANOVA setups were used where the F1-classification score for each experiment, cell and associated ground-truth cell-type was tested:

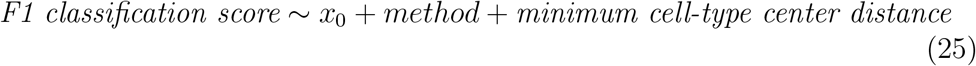

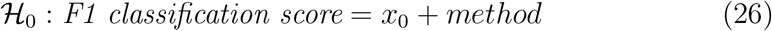

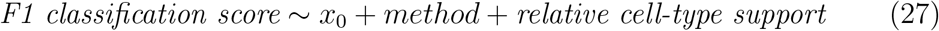

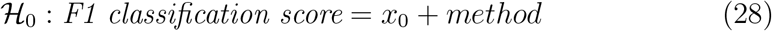

Where:

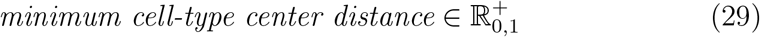

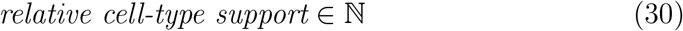

The *cell-type analyzed* was not included as a factor to control, because the *minimum cell-type center distance* and *relative cell-type support* metrics were calculated on a **per cell-type basis**. Therefore, these metrics are perfectly collinear with cell-type, and this would absorb the residuals that would be picked up by the key metrics.

##### 6.3.2 PDAC perturbation analysis

Perturbations were performed for the compartmentalized PDAC data (1.3 and 3.2) to determine the effects of downsampling/ablation on the classification scores of all compartments. Here, the following ANOVA setup was used to determine the effects on F1-scores **for a specific compartment** based on **downsampling of the same compartment** for each compartment *c*:

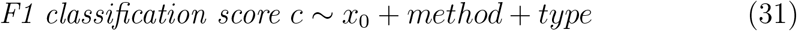

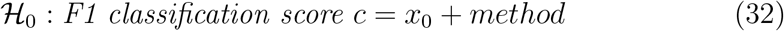

These results were analyzed independently and jointly for all compartments downsampled, where joint-comparison included comparison of F-values for perturbation in each setup.

### 7 Balanced clustering scores

None of the utilized clustering metrics, which in this analysis and other integration benchmarking/methods papers are used to compare the concordance of ground-truth cell-type labels and unsupervised clusters attained in an embedding, factor in class balance. The metrics utilized include: the Adjusted Rand Index (ARI), Adjusted Mutual Information (AMI), Homogeneity Score, and Completeness Score. The implementation details of these metrics can be found in the scikit learn documentation [32].

Strictly speaking, the **Homogeneity Score and Completeness Scores are not metrics, because they are not symmetric**. However, this symmetry is not necessary in the case of single-cell benchmarking, and the general case of comparing clustering labels with ground-truth annotations, because one set of labels is known to be ground-truth. In fact, balancing the ARI and AMI will break their symmetry as well.

To introduce the procedure behind reweighing these metrics, we’ll begin with the balanced ARI. Then we’ll extrapolate this procedure to the entropy-based metrics/scores (AMI, Homogeneity, and Completeness), as this extrapolation only involves a slight modification to the ARI procedure for these scores.

Code notebooks on implementing the balanced clustering scores with usage demonstrations and relevant examples are available at https://github.com/hsmaan/balanced-clustering/tree/main/notebooks.

#### 7.1 The *Balanced Adjusted Rand Index*

##### 7.1.1 The Rand Index and Adjusted Rand Index

For a set of *n* objects, *S* = {*O*_1_, *O*_2_, *O*_3_, …, *O_n_*}, the goal of clustering is to partition these objects into meaningful subsets, which we can call partitioning *V*. Assuming we have access to either ground-truth labels or clusters from another technique, which we can denote partitioning *U*. Both *U* and *V* contain subsets, which we call either classes or clusters: *U* = {*u*_1_, *u*_2_, …, *u_R_*} and *V* = {*v*_1_, *v*_2_, …, *v_C_*}. These clustering results are subject to the following constraints to be valid for calculating the Rand Index:

1. All *n* objects within the set *S* must be within sets *U* and *V*:

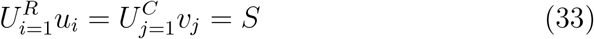
2. No element from set *S* can belong in to more than one subset in either *U* or *V*

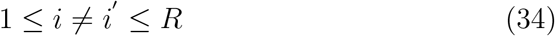

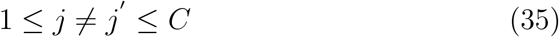

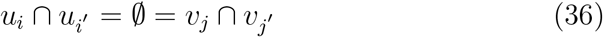

To quantify the overlap between the partitions *U* and *V* (e.g. in determining overlap between a set of ground-truth labels and the results of clustering), we can start by creating a contingency table which indicates the overlap:

| | $v_1$ | $v_2$ | $v_3$ | … | $v_c$ | |
| --- | --- | --- | --- | --- | --- | --- |
| $\mathbf{u_1}$ | $t_{11}$ | $t_{12}$ | $t_{13}$ | … | $t_{1C}$ | $\mathbf{t_{1.}}$ |
| $\mathbf{u_2}$ | $t_{21}$ | $t_{22}$ | … | … | … | … |
| $\mathbf{u_3}$ | $t_{31}$ | … | … | … | … | … |
| … | … | … | … | … | … | … |
| $\mathbf{u_R}$ | $t_{R1}$ | … | … | … | $t_{RC}$ | $\mathbf{t_{R.}}$ |
| | $\mathbf{t_{.1}}$ | … | … | … | $\mathbf{t_{.C}}$ | $t_{..}$ |

Each element of this table indicates overlapping elements. E.g. *t*_11_ indicates the number of samples that have the label *v*_1_, in *V* and *u*_1_, in *U*. The total number of values in the matrix is 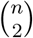 if *n* objects/samples are present. As we now have a table/matrix that represents the overlap of assignments to subsets in *U* and *V* for *n* objects, we can determine the concordance of partitions *U* and *V* for these objects using the Rand Index:

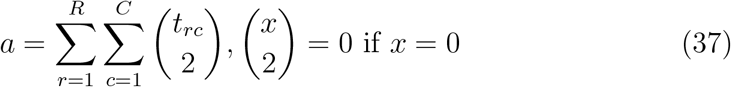

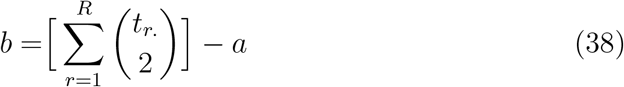

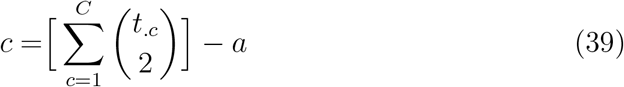

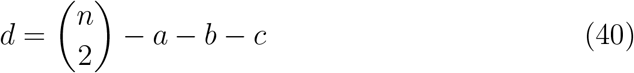

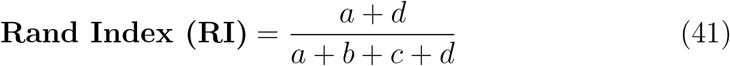

Intuitively, the Rand Index aims to calculate how many pairs are concordantly in the same subsets in *V* and *U* (*a*), how many pairs are concordantly in different subsets in *V* and *U*, and how many are discordant (in the same group in one partition and otherwise in the other). It’s important to note that **pairs here refer to all combinations of two different objects, not the same object being considered in the two partitions**.

Although the Rand Index is normalized (lower bound = 0, upper bound = 1), it is not adjusted for chance clustering. A correction can be made [26] to the RI formula that takes into account the **expected value of the RI for two partitions of the objects** *U* **and** *V*, denoted by the Adjusted Rand Index (ARI) [26]:

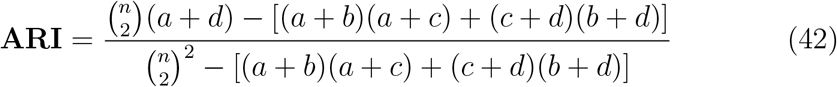

With this correction, the ARI is a metric (symmetric, positive-definite, and the triangle inequality) [27] that has the properties of **normalization** and **expectation** [26].

##### 7.1.2 Balancing the ARI

Rebalancing the ARI (as well as the other entropy-based scores/metrics) will amount to **rescaling the total number of values in the subsets of the partition we consider ground-truth**. In this case, **assume** *U* **is the partition with the ground-truth information**. We want each subset from *U* to have an equal contribution to the ARI value - this is concomitant with each class from the ground-truth data (which we have assumed to be *U*) having an equal weighing in the calculating of the score. This can be done in the following step-wise manner:

1. Determine contribution of each subset of *U* (*t*_1_, *t*_2_, …, *t_R_*) to score through mean of marginals from contingency table:

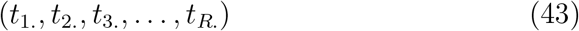
2. Get the mean contributions of all subsets:

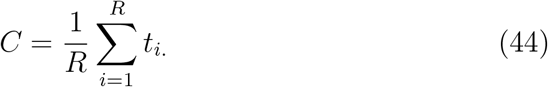
3. For each subset (*t_i_*) of *U*, normalize the contribution to be equal to the mean using a scaling factor:

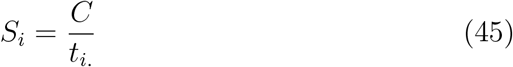

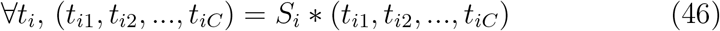

After these steps, we’ve essentially rescaled the contingency table such that the contribution from each subset in *U* will be considered equally in calculations using the table results. To calculated the *Balanced Adjusted Rand Index*, we can apply the 7.1.2 normalization procedure and use the same ARI formula as before (42):

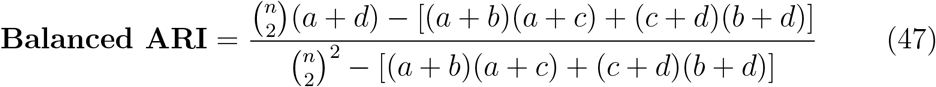

Examining the values needed to calculate the RI and ARI (37), we can see that this normalization procedure will effectively rescale the calculations for *a*, *b*, and *c*. This procedure does this while still retaining the total counts (*n*), such that the calculation for *d* will be unaffected. Because the RI and ARI calculations simply depend on these values in the contingency table that can be calculated independently, the application of this normalization procedure is straightforward and does not require any further steps.

#### 7.2 Balancing entropy-based scores

##### 7.2.1 Mutual information

Central to the Adjusted Mutual Information (AMI), Homogeneity, and Completeness scores is the calculation of mutual information between partitions *U* and *V* [27] [28]. For the contingency table previously defined in 47, the mutual information between these two partitions is equal to the following [27]:

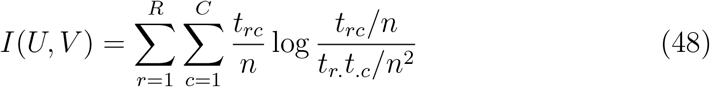

We’ll follow the same normalization procedure that we did in 7.1.2, as we are starting from the same contingency table of overlapping objects in subsets of partitions *U* and *V*. From here, we can calculate the mutual information value and proceed with the rest of the calculations for the entropy-based scores.

##### 7.2.2 Entropy

Aside from mutual information, the other important factor that is used by all of the entropy-based scores is the calculation of the entropy of the labelling - i.e. how ordered/disordered are the objects in partitions *U* and *V*. This can also be calculated from the contingency table from 47 in the following manner [27]:

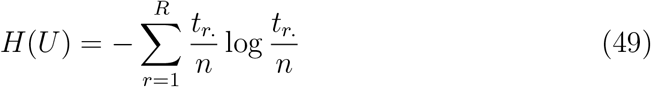

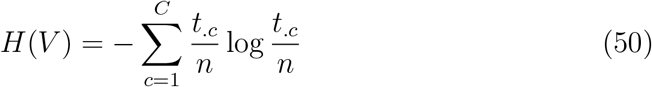

The Homogeneity and Completeness scores also require the conditional entropy formulation [28] [27]:

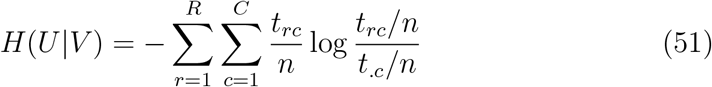

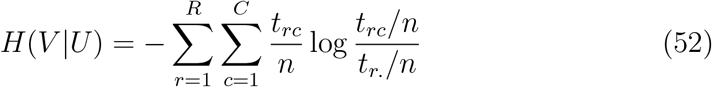

##### 7.2.3 Balanced entropy-based scores

The calculation of the entropy and mutual information can proceed as-is after the normalization procedure from 7.1.2, and this will balance the contributions from a presumed ground-truth partition *U* in calculating the entropy and mutual information. From here the Balanced Adjusted Mutual Information, Balanced Homogeneity, and Balanced Completeness scores can be calculated using these two values, the rescaled contingency matrix after 7.1.2, and the base formulas for these scores [27] [28]:

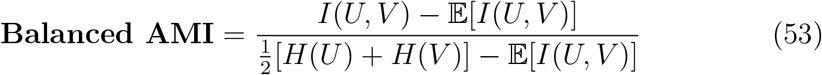

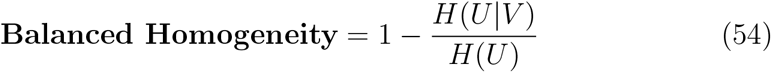

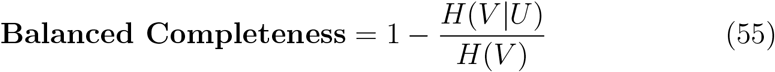

The V-measure and Balanced V-measure are simply the harmonic mean of the Completeness and Homogeneity scores [28]:

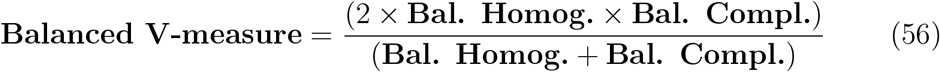

#### 7.3 Balanced clustering evaluations

The following section details the evaluations that were utilized for the balanced clustering metric analysis. Seeding was set for all of these cases to ensure reproducibility of the simulations, downsampling, and integration methods (where possible).

##### 7.3.1 3 imbalanced well-separated classes, 2 clusters

In this scenario, 3 well separated but imbalanced classes were utilized and a mis-clustering of the smaller class was done with k-means clustering with k=2. This data was simulated using 2D Gaussian densities with the following values for each class:

- Class A ~ *N*(0, 0.5) - 500 samples
- Class B ~ *N*(−2, 0.1) - 20 samples
- Class C ~ *N*(3, 1) - 500 samples

K-means clustering with k=2 led to class B overlapping with class A in the clustering result.

The balanced and imbalanced metrics were compared when calculating the concordance of the ground-truth labels (class labels) and k-means clustering labels.

##### 7.3.2 3 imbalanced overlapping classes, 3 clusters

In this case, 3 classes that are overlapping and imbalanced (2 smaller classes on edges of larger class) were analyzed, and k-means clustering with k=3 was done and the result correctly clustered most of the samples from the smaller classes, but due to slicing of the larger class present because of overlap, mis-clustered a large number of majority class samples.

This data was simulated using 2D Gaussian densities with the following values for each class:

- Class A ~ *N*(0, 0.5) - 1500 samples (larger class)
- Class B ~ *N*(1, 1) - 200 samples
- Class C ~ *N*(−1, 1) - 200 samples

##### 7.3.3 Balanced 2 batch PBMC - co-clustered CD4+ T cells and CD8+ T cells

The balanced 2 batch PBMC dataset was utilized here (1.2). Batch 1 was kept as is, and batch 2 had all of the cells ablated except for CD4+ T cells, which were downsampled to 10% of their original proportion.

The default Leiden clustering resolution of 1 in the scanpy implementation was changed to 0.1, as this value perfectly clusters all of the cell-types with the exception of the CD4+ T cells, which get collapsed into a cluster with CD8+ T cells, simulating a case where a smaller cell-type is co-clustered with a larger cell-type.

The resultant embedding with no integration was utilized, and the ground-truth cell-type labels and unsupervised clustering labels were used to compare the balanced and imbalanced/vanilla scores - where the ARI and Homogeneity scores were shown.

##### 7.3.4 Balanced 2 batch PBMC data - downsampled CD4+ T cells and FCGR3A+ monocytes

In this evaluation, the 2 batch balanced PBMC dataset was once again utilized. For the two batches, each one had either the CD4+ T cells or FCGR3A+ monocytes downsampled to 10% of their original population, creating an imbalanced scenario specific to these two cell-types.

After this, integration was done using BBKNN, Harmony, Scanorama, and scVI. The same integration pipeline from 2 was utilized. An ’unintegrated’ control subset was used, where the pipeline from 2 was followed without integration with any method.

From here, the average value of the balanced and imbalanced metrics was used for comparison. e.g.:

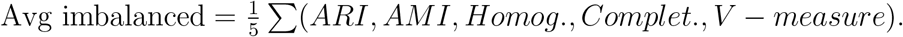

### 8 Code and data availability

The python package for implementing the balanced clustering metrics can be found here:

https://github.com/hsmaan/balanced-clustering

All of the code necessary to reproduce the results of the Iniquitate pipeline are available at:

https://github.com/hsmaan/Iniquitate

The datasets utilized in this study, which are associated with the various configurations used in the Iniquitate GitHub repository, can be all found here:

https://drive.google.com/file/d/102ntQuclUzQILRxMVXo1-yQR43t97Q3r/view?usp=sharing

This directory is in the exact necessary structure needed to run the Iniquitate pipeline, and can be copied into the cloned GitHub repository for Iniquitate under **Iniquitate/resources**. Instructions are also given in the Iniquitate GitHub link.

## Supporting information

Main figures 1-8 full pdfs

Supplementary figures 1-16 full pdfs

## 9 Appendix A: Python and R library/package version numbers

The following two environments (9.1, 9.2) were used in the benchmarking and analysis phases, where all integration experiments and downstream analysis tests were done with the pipeline environment (9.1), and all results analysis and plotting was done with the analysis environment (9.2). The only exception were the balanced metric analyses and tests (7), which utilized the analysis environment (9.2) for generation and testing of the various scenarios outlined.

Configurations for these environments are also available at https://github.com/hsmaan/Iniquitate/tree/main/workflow/envs.

The library for the balanced metrics (7) was developed independently, and all of the information on dependency versions is available at https://github.com/hsmaan/balanced-clustering.

### 9.1 Iniquitate pipeline environment

- python>=3.7,<=3.10
- numpy>=1.19.0
- pandas>=1.2.0
- scipy>=1.5.0
- leidenalg>=0.8.0
- umap-learn>=0.5.0
- mnnpy>=0.1.9
- scikit-learn>=1.0.1
- scanpy=1.8.2
- anndata=0.8.0
- faiss-cpu>=1.7.0
- pytorch=1.10.1
- torchmetrics<=0.6.0
- cudatoolkit=10.2
- scvi-tools=0.14.4
- bbknn=1.5.1
- harmonypy=0.0.5
- scanorama=1.7.1
- r-base>=4.0.0
- r-liger=0.5.0
- r-seurat=4.0.6
- r-seuratdisk>=0.0.9
- r-data.table>=1.14.0
- r-reticulate=1.24
- cython>=0.29.25
- r-rann=2.6.1

### 9.2 Analysis scripts environment

- python>=3.7,<=3.10
- numpy>=1.19.0
- pandas>=1.2.0
- scipy>=1.5.0
- seaborn>=0.11.2
- plotnine>=0.8.0
- leidenalg>=0.8.0
- umap-learn>=0.5.0
- scikit-learn>=1.0.1
- scanpy=1.8.2
- anndata>=0.7.5
- ipykernel>=6.4.0
- jupyterlab>=3.2.9
- notebook>=6.4.2
- scvi-tools=0.14.4
- pytorch=1.10.1
- torchmetrics<=0.6.0
- cudatoolkit=10.2
- bbknn=1.5.1
- harmonypy=0.0.5
- scanorama=1.7.1
- r-base>=4.0.5
- r-seurat>=4.0.5
- r-data.table>=1.14.0
- r-ggplot2>=3.3.0
- r-tidyverse>=1.2.1
- r-reshape2>=1.4.3
- r-data.table>=1.14.0
- r-ggthemes>=4.2.0
- r-ggextra>=0.8.0
- r-dotwhisker>=0.7.4
- r-seuratdisk>=0.0.9019
- r-deldir>=1.0.2
- r-ggpubr>=0.4.0
- r-cowplot>=1.1.1
- r-ggrepel>=0.9.1
- r-rcolorbrewer>=1.1
- r-ggbump>=0.1.0
- bioconductor-complexheatmap<=2.9.0
- r-venndiagram>=1.7.1
- r-multipanelfigure>=2.1.2
- r-gridextra>=2.3
- r-cairo>=1.5
- r-lemon>=0.4.5
- r-networkd3>=0.4
- r-emt>=1.2
- cython>=0.29.25

## Supplementary Figures

**Figure S1:**
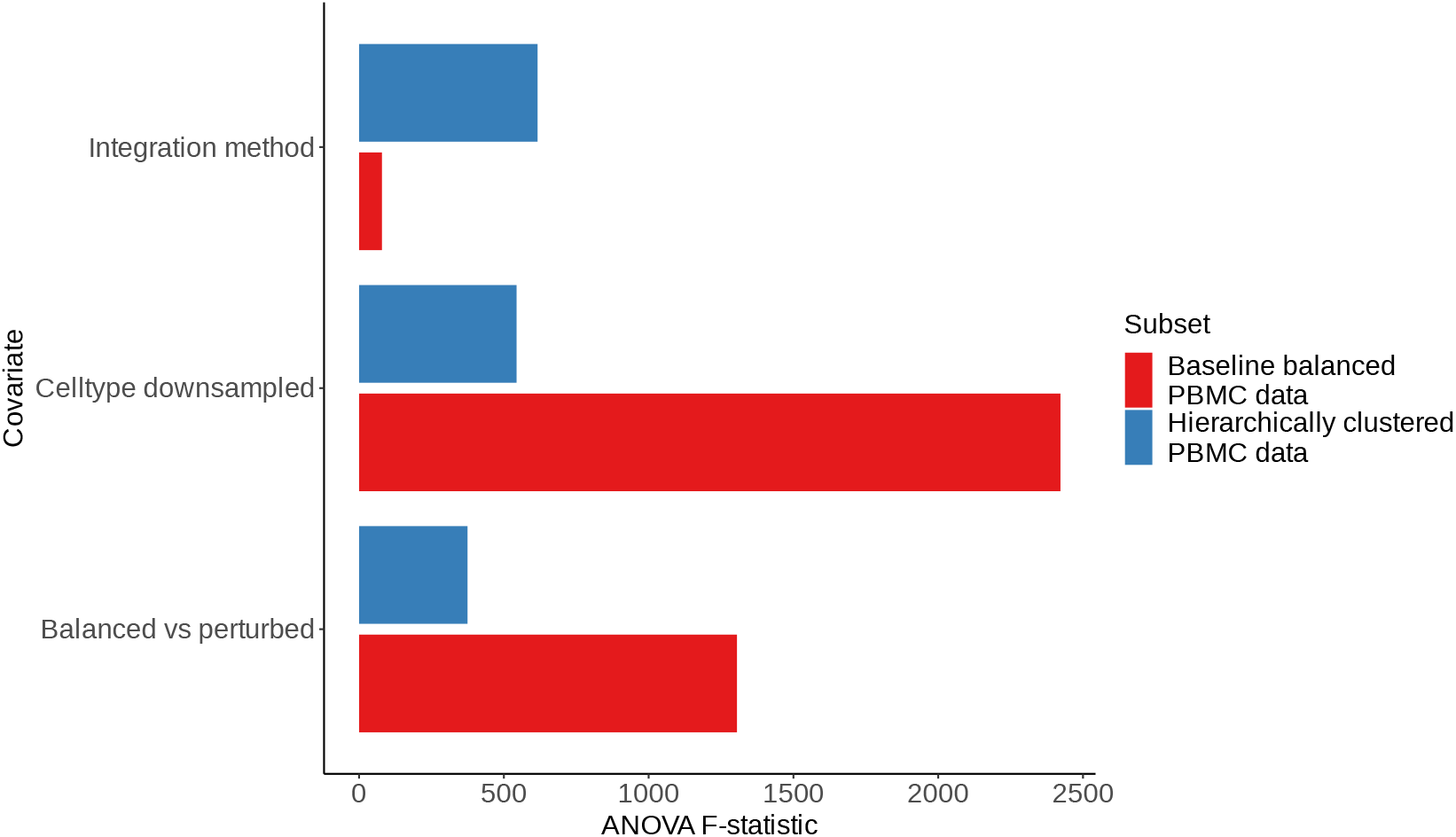
ANOVA F-statistic values for cell-type specific KNN classification in the baseline and hierarchical 2 batch balanced PBMC data. The ANOVA F-statistic values, indicating the ratio of variation between between sample means and variation within the samples themselves, for the covariates used in the KNN-classification task ANOVA for the 2 batch PBMC balanced dataset (Online Methods). F-statistics are shown for integration method (first covariate in model), cell-type that was downsampled (second covariate in model), and which type of experiment was performed (control balanced vs. perturbed - last covariate in model). The F-statistics are compared between the baseline setup (6 cell-types initially utilized) and the hierarchical setup after merging closely related cell-types.

**Figure S2:**
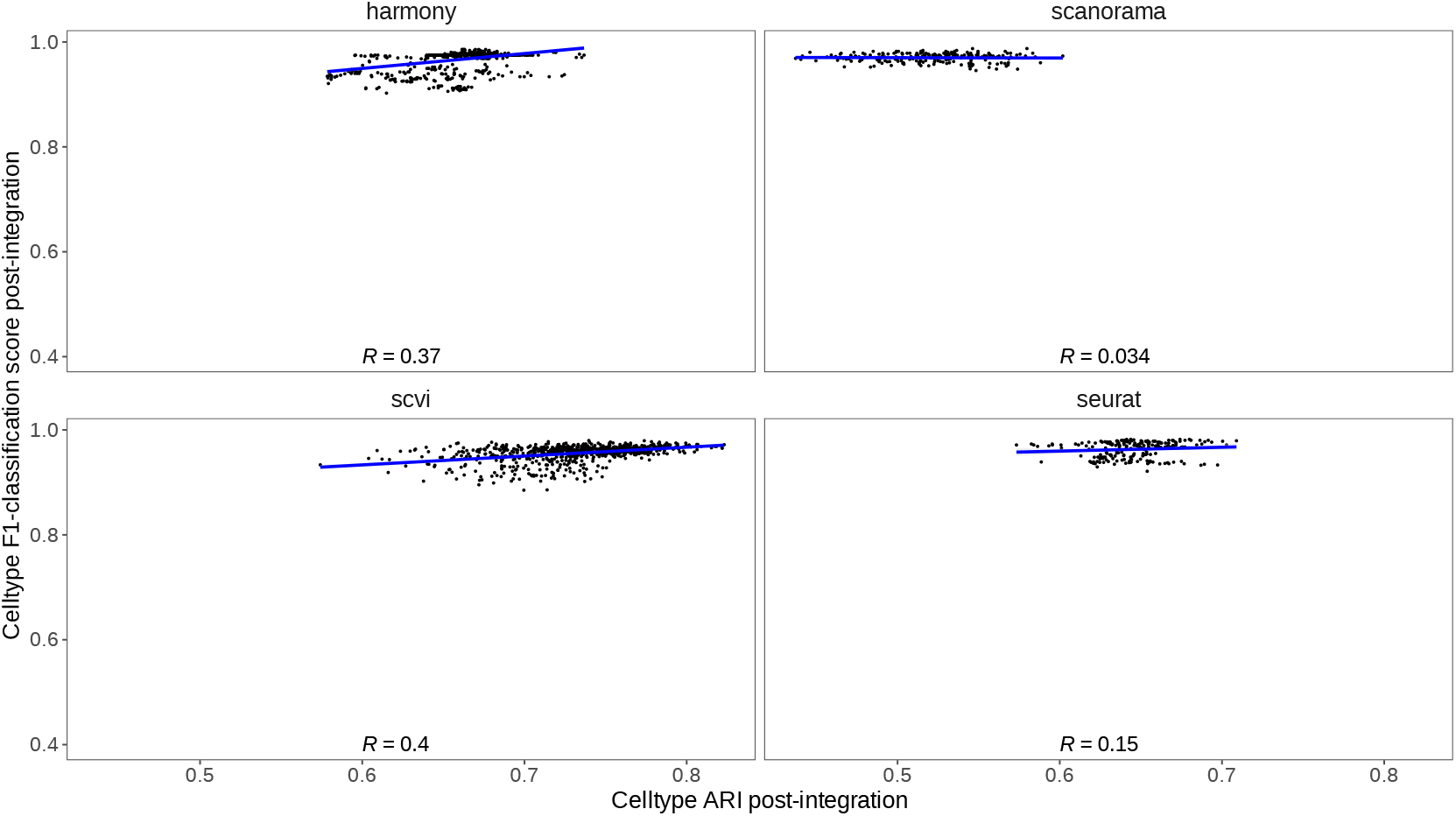
Correlation between cell-type-specific F1-classification scores and cell-type ARI_*cell–type*_ in balanced 2 batch PBMC data. All experiments (control, downsampling, and ablation) are indicated for the baseline 2 batch PBMC data. For perturbation experiments, values are subset for only where the cell-type being classified (F1-classification score) is equivalent to the cell-type that was down-sampled. The median cell-type classification F1-score across all cell-types is shown, grouped by method and experiment type, for direct comparison with the ARI values which are calculated per replicate. The Spearman correlation value between the scores is indicated.

**Figure S3:**
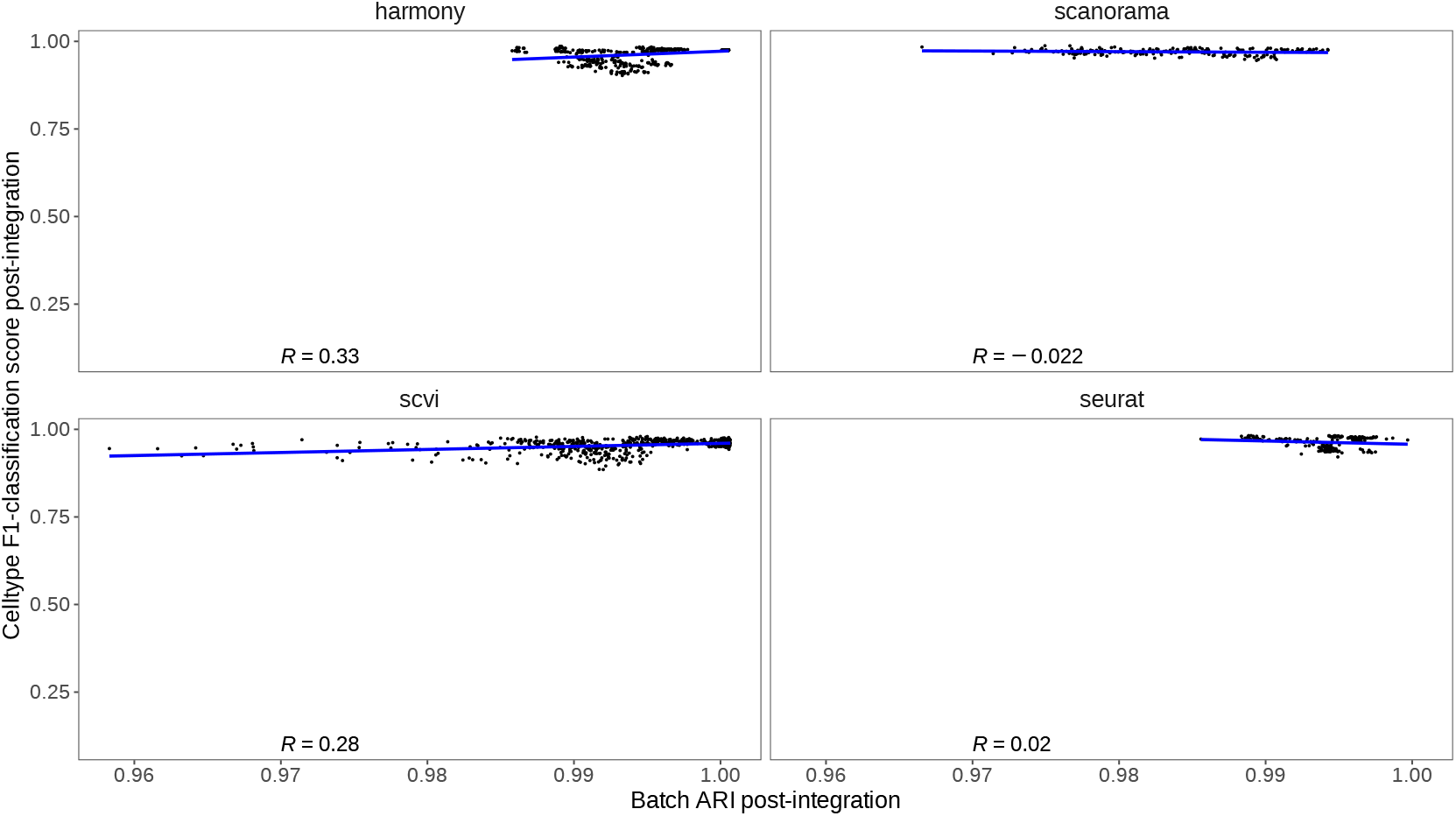
Correlation between cell-type-specific F1-classification scores and (1 - ARI_*batch*_) values in balanced 2 batch PBMC data. All experiments (control, downsampling, and ablation) are indicated for the baseline 2 batch PBMC data. For perturbation experiments, values are subset for only where the cell-type being classified (F1-classification score) is equivalent to the cell-type that was downsampled. The median cell-type classification F1-score across all cell-types is shown, grouped by method and experiment type, for direct comparison with the ARI values which are calculated per replicate. The Spearman correlation value between the scores is indicated.

**Figure S4:**
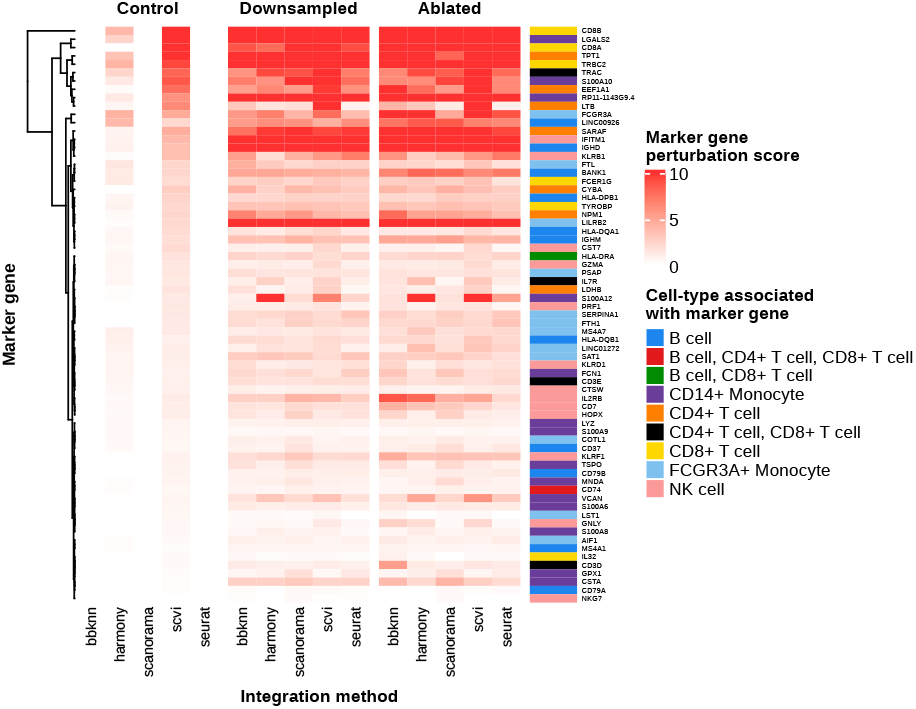
marker gene perturbation scores across all marker genes for the cell-types in the balanced 2 batch PBMC dataset. Marker genes were determined through differential gene expression analysis within each batch (Online Methods), and their perturbation score, indicating change in maximum ranking across unsupervised clusters post-integration are shown across control, downsampling, and ablation experiments (Online Methods). Note that downsampling and ablation (perturbation) experiments are not subset here for the marker gene being analyzed and its associated cell-type (e.g. maximum-rank change for B-cell markers in only runs where B-cells are downsampled).

**Figure S5:**
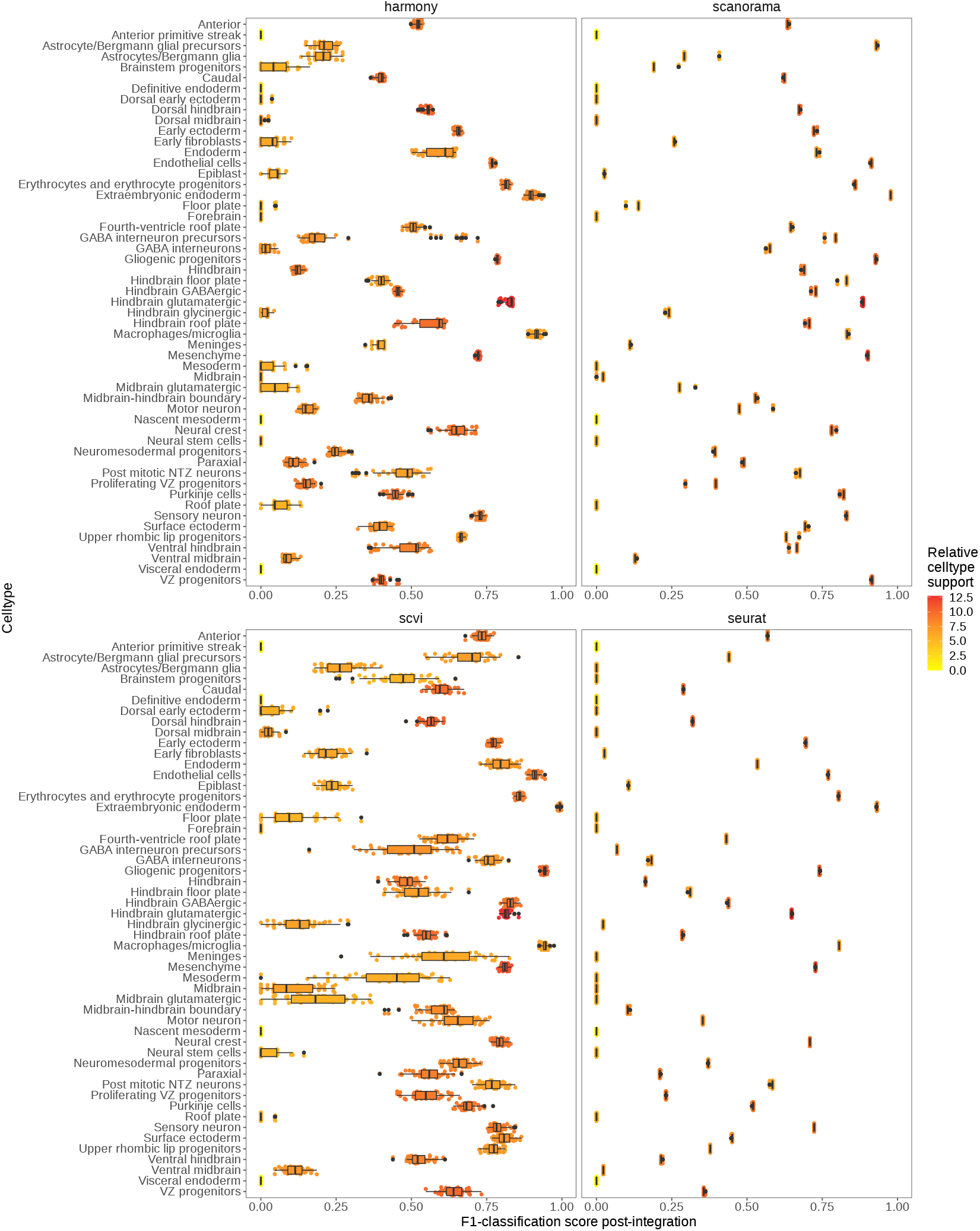
Comparison of F1-classification accuracy and relative cell-type support of each cell-type in the imbalanced 6 batch mouse hindbrain development dataset. The relative cell-type support is based on the number of cells in the integrated embedding space present for each cell-type (Online Methods).

**Figure S6:**
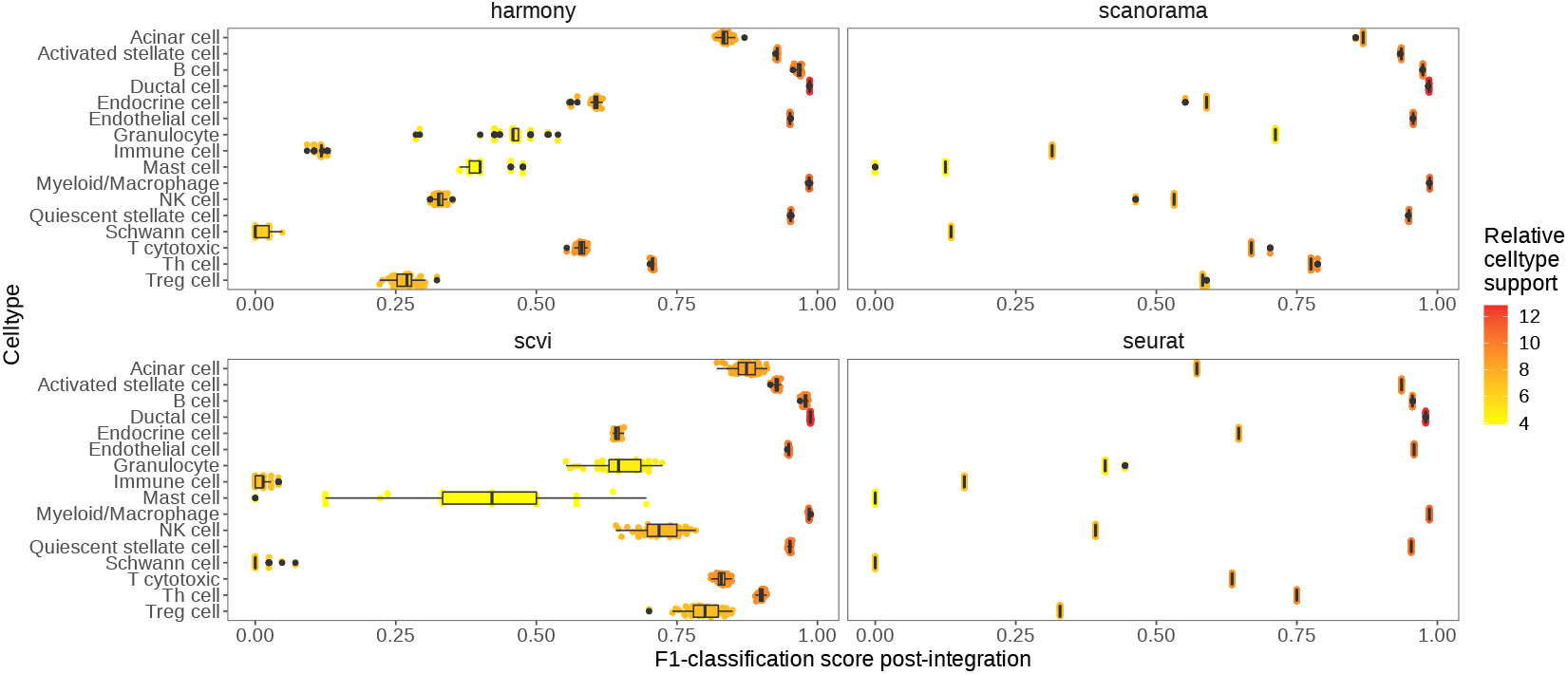
Comparison of F1-classification accuracy and relative cell-type support of each cell-type in the imbalanced 8 batch PDAC dataset. The relative cell-type support is based on the number of cells in the integrated embedding space present for each cell-type (Online Methods).

**Figure S7:**
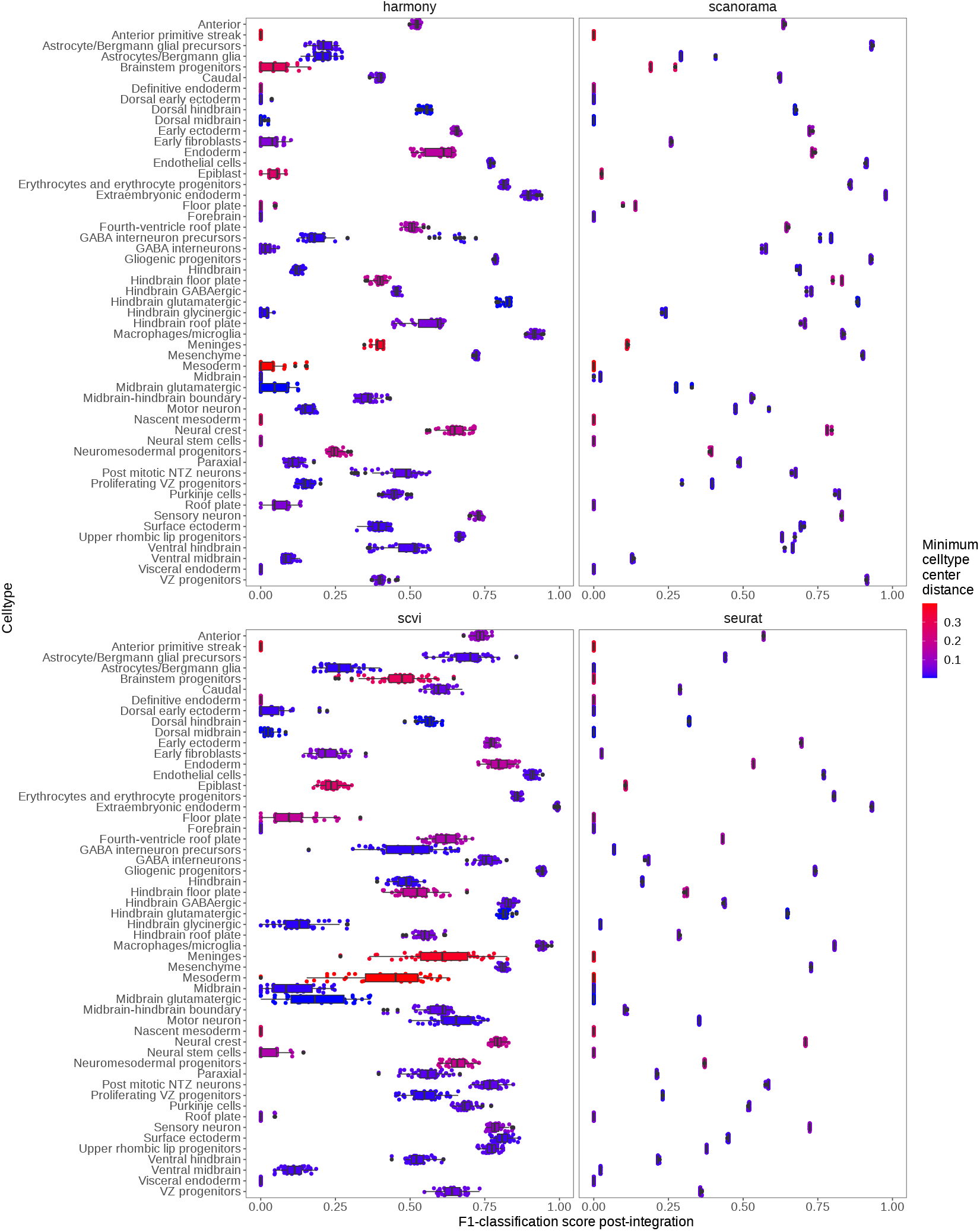
Comparison of F1-classification accuracy and minimum celltype center distance of each cell-type in the imbalanced 6 batch mouse hindbrain development dataset. The minimum cell-type center distance value indicates how close is the closest other cell-type across batches in PCA space (Online Methods).

**Figure S8:**
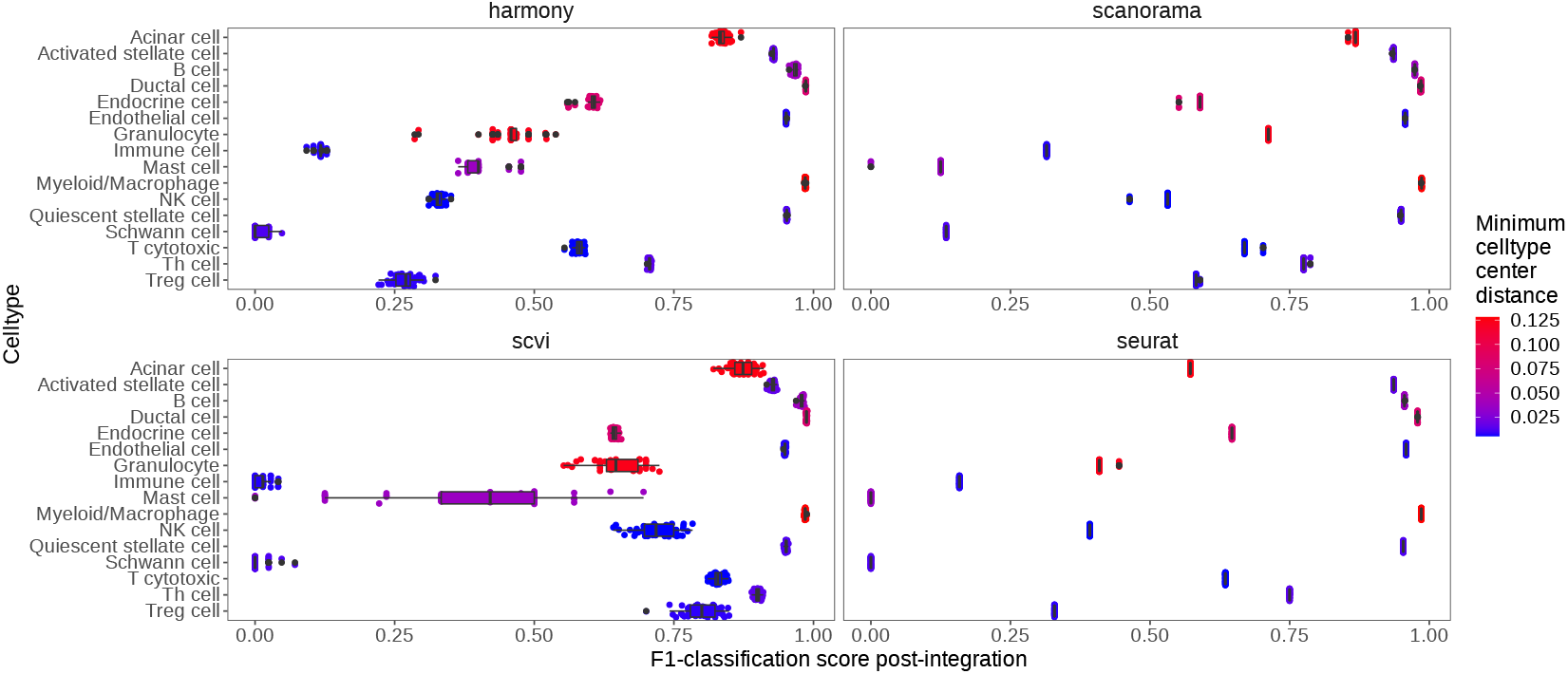
Comparison of F1-classification accuracy and minimum celltype center distance of each cell-type in the imbalanced 8 batch PDAC dataset. The minimum cell-type center distance value indicates how close is the closest other cell-type across batches in PCA space (Online Methods).

**Figure S9:**
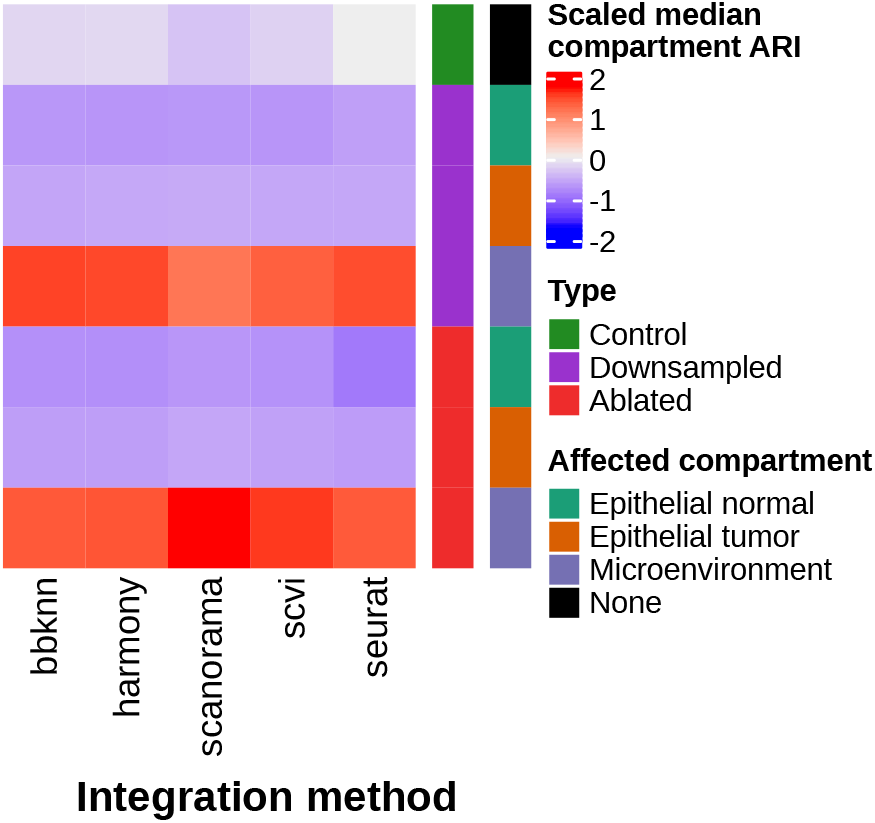
Comparison of compartment heterogeneity conservation ARI results across PDAC data perturbation experiments. Z-score normalized median ARI_*compartment*_ (compartment integration accuracy) results across experiment type (control, compartment downsampling, compartment ablation), specific-compartment downsampled, and integration method utilized.

**Figure S10:**
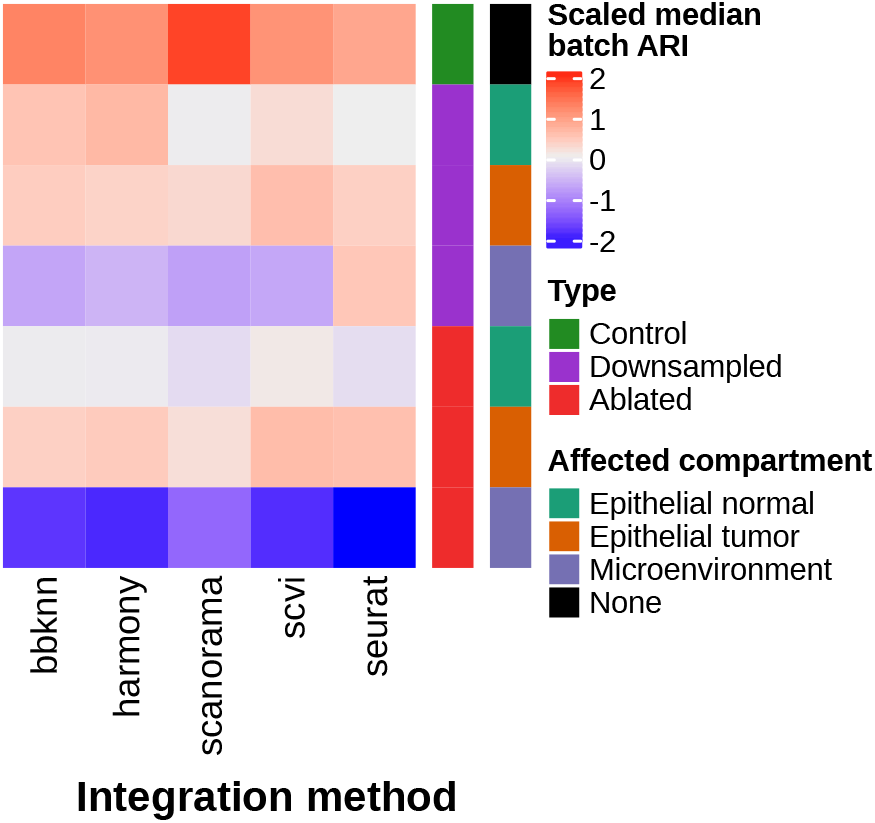
Comparison of batch-mixing ARI results across PDAC data perturbation experiments. Z-score normalized median (1-ARI_*batch*_) (batch mixing) results across experiment type, compartment downsampled, and integration method.

**Figure S11:**
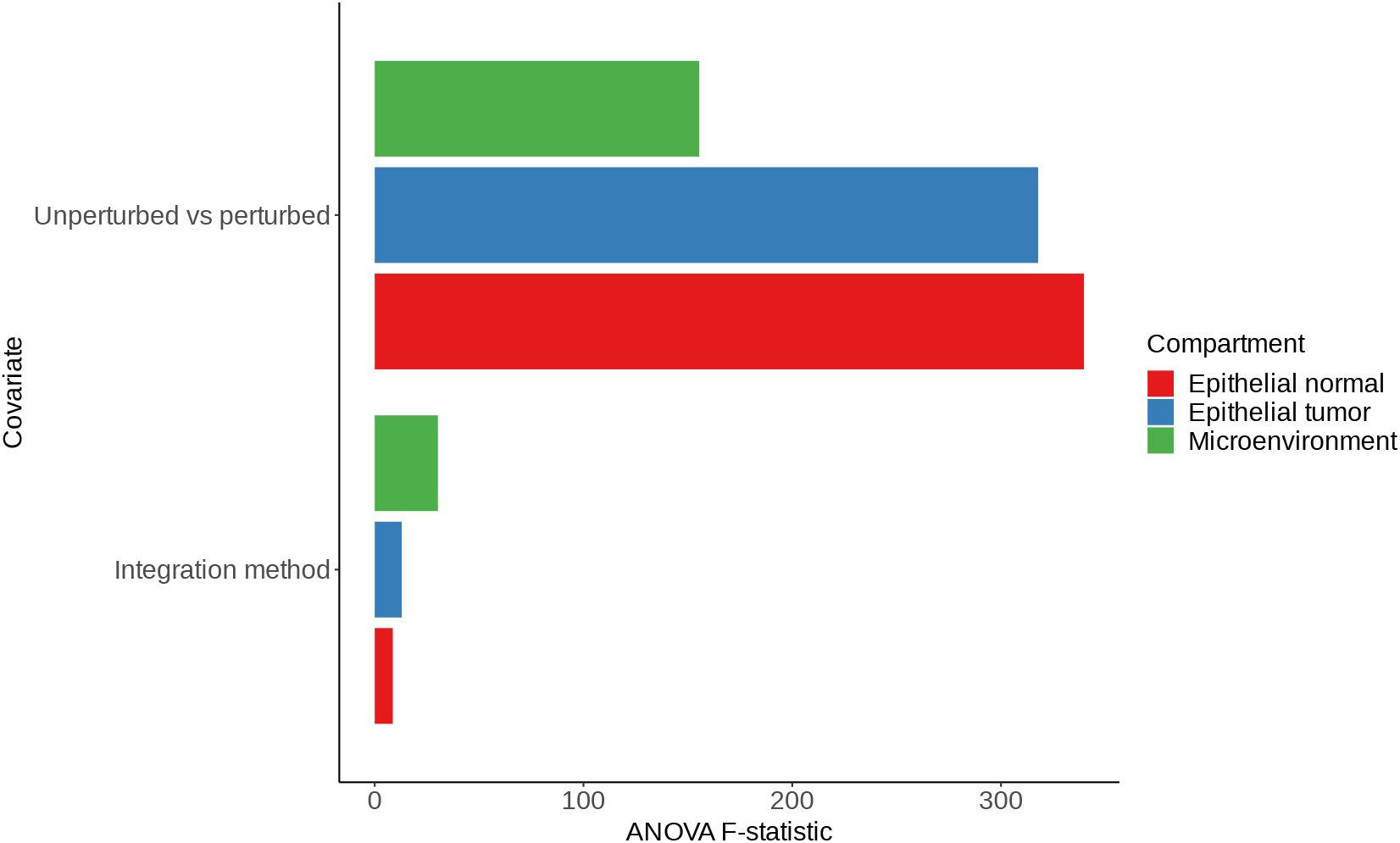
ANOVA F-statistic values for compartment specific KNN-classification in the 8 batch compartmentalized PDAC data. The ANOVA F-statistic values, indicating the ratio of variation between sample means and variation within the samples themselves, for KNN-classification of individual compartments before and after perturbation (Online Methods). F-statistics are shown for integration method (first covariate in model), and which type of experiment was performed (control vs. perturbed - last covariate in model). The ANOVA tests were performed individually for each compartment, and the compartment-specific F-statistics are shown. Note that the perturbations here are specific to the compartment being analyzed (e.g. microenvironment subset will only contain perturbations that targeted the microenvironment) (Online Methods).

**Figure S12:**
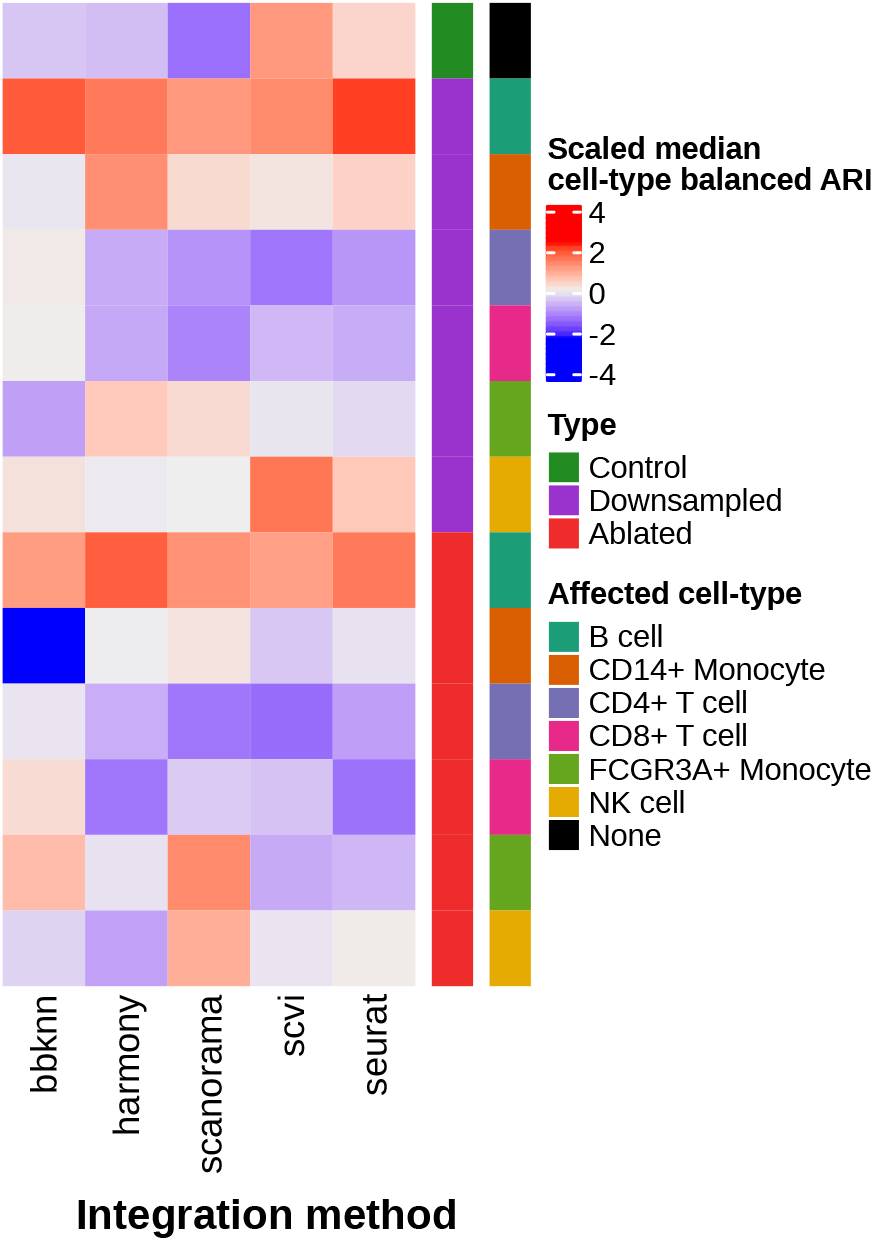
Comparison of cell-type heterogeneity conservation ARI results across the PBMC 2 batch perturbation experiments, using the balanced ARI (bARI) score. Z-score normalized median ARI_*cell–type*_ (cell-type integration accuracy) results across experiment type (control, compartment downsampling, compartment ablation), specific-cell-type downsampled, and integration method utilized, using the bARI instead of the base ARI metric.

**Figure S13:**
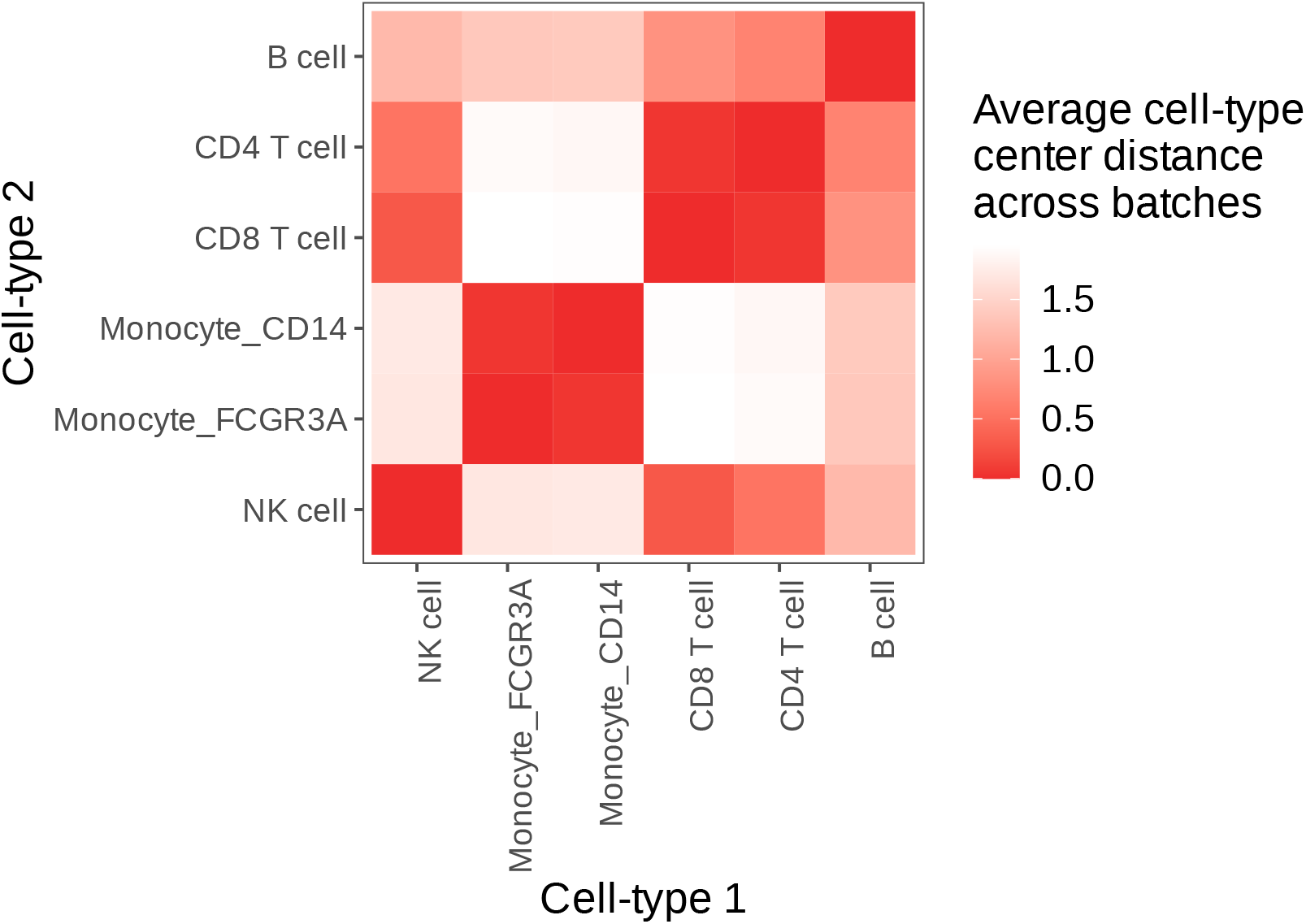
Average cell-type center distance between cell-types in the balanced PBMC 2 batch dataset. For each batch, the distance from the centers of cell-type clusters in principal component analysis (PCA) reduction space are calculated, and the relative distances between cell-types are determined and averaged across batches (Online Methods).

**Figure S14:**
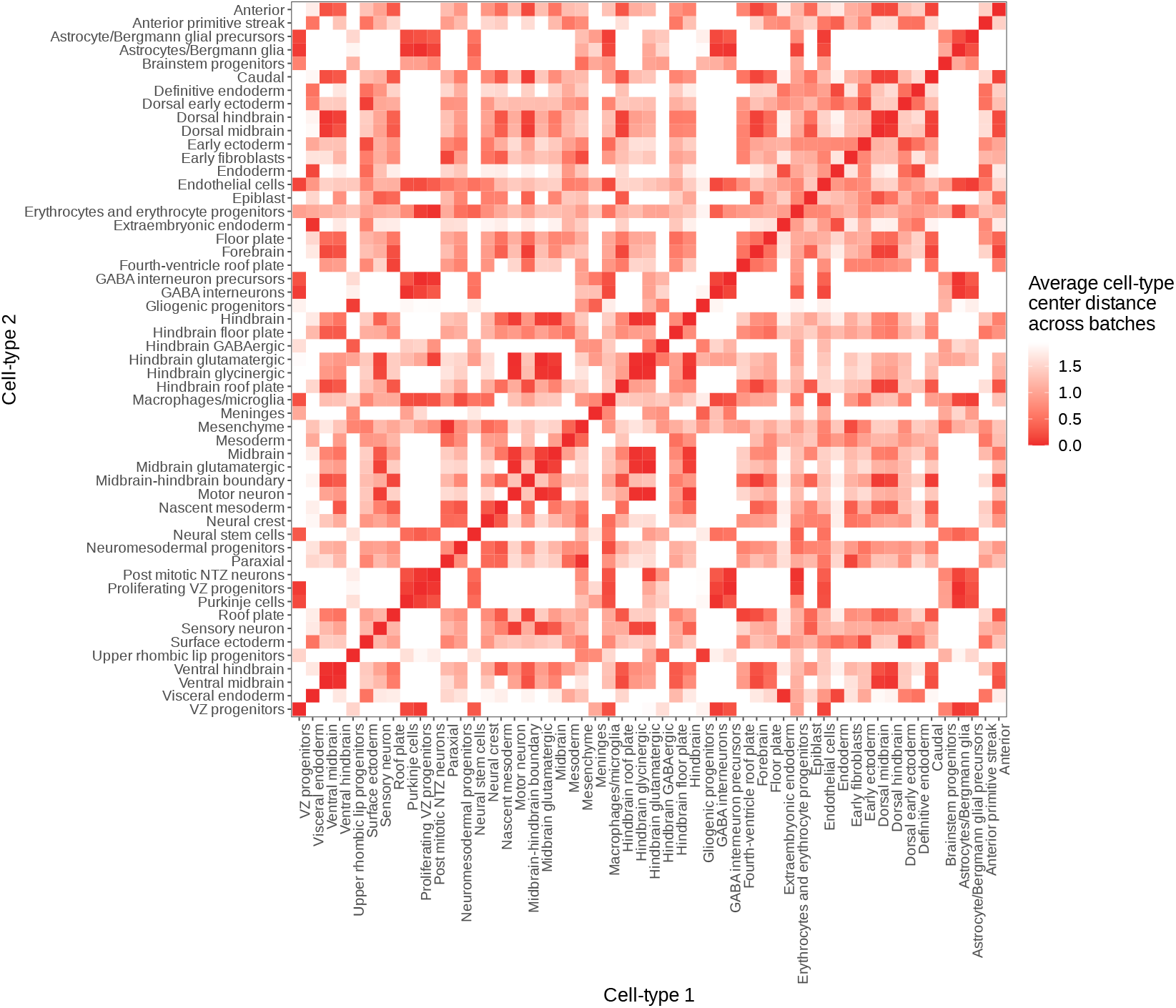
Average cell-type center distance between cell-types in the 6 batch mouse hindbrain development dataset. For each batch, the distance from the centers of cell-type clusters in principal component analysis (PCA) reduction space are calculated, and the relative distances between cell-types are determined and averaged across batches (Online Methods).

**Figure S15:**
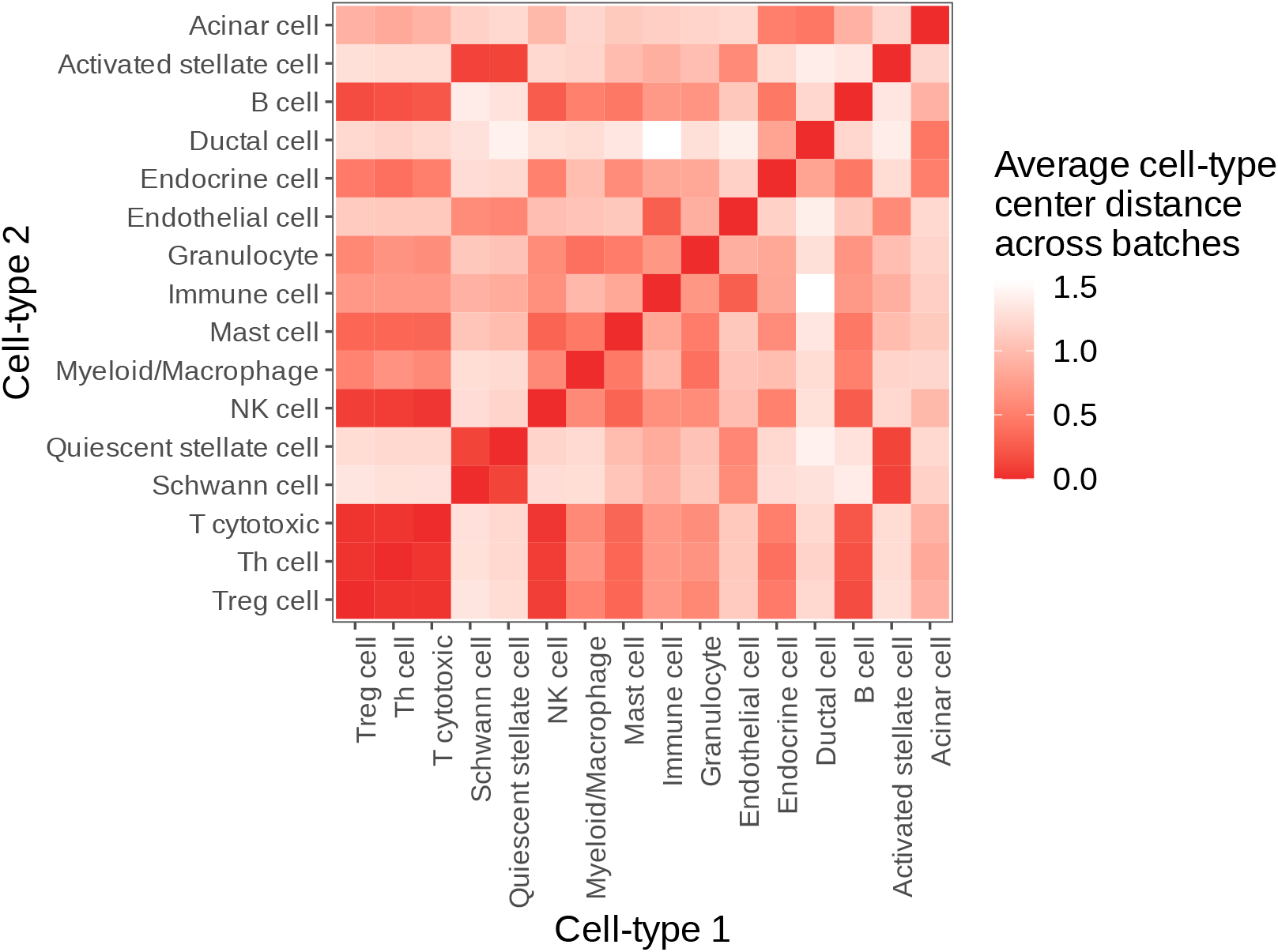
Average cell-type center distance between cell-types in the 8 batch pancreatic ductal adenocarcinoma (PDAC) dataset. For each batch, the distance from the centers of cell-type clusters in principal component analysis (PCA) reduction space are calculated, and the relative distances between cell-types are determined and averaged across batches (Online Methods).

**Figure S16:**
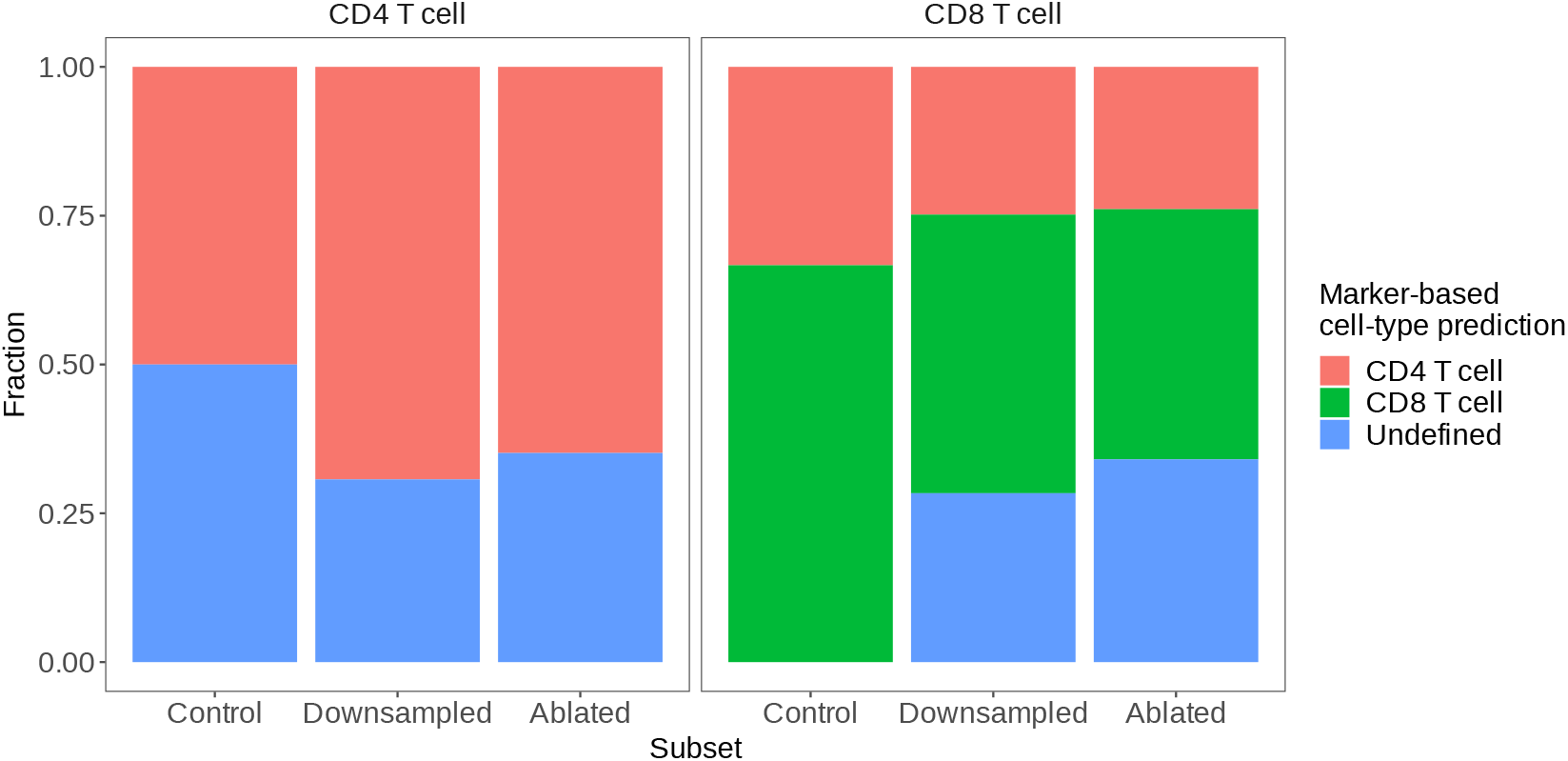
Predicted cell-types for CD4/CD8 T-cell majority clusters in the balanced PBMC 2 batch data, based on marker gene differential expression. In this setup, the top 50 marker genes were analyzed based on differential expression for unsupervised clusters from the Seurat integration method across experimental subsets (Control, Downsampling, Ablation). The Downsampling and Ablation subsets here contain only instances where CD4+ and CD8+ T cells were affected. Only clusters that contained a majority of cells (based on ground-truth annotations) of CD4+ or CD8+ T were kept. The canonical marker genes for CD4+ T cells (IL7R) and CD8+ T cells (CD8A) were used to predict the cell-type for each cluster based on their relative ranking in the top 50 marker genes for the given clusters (Details in Online Methods: Downstream analysis - marker gene ranking - Case study - CD4/CD8 T cell assignment based on marker genes). The fraction of clusters that contain a majority of CD4+ or CD8+ T cells, and their predicted cell-type based on the aforementioned marker gene setup are indicated, across experimental subsets.

