## Supplementary figures and images for "The differential impacts of dataset imbalance in single-cell data integration"

### Figure_1.pdf

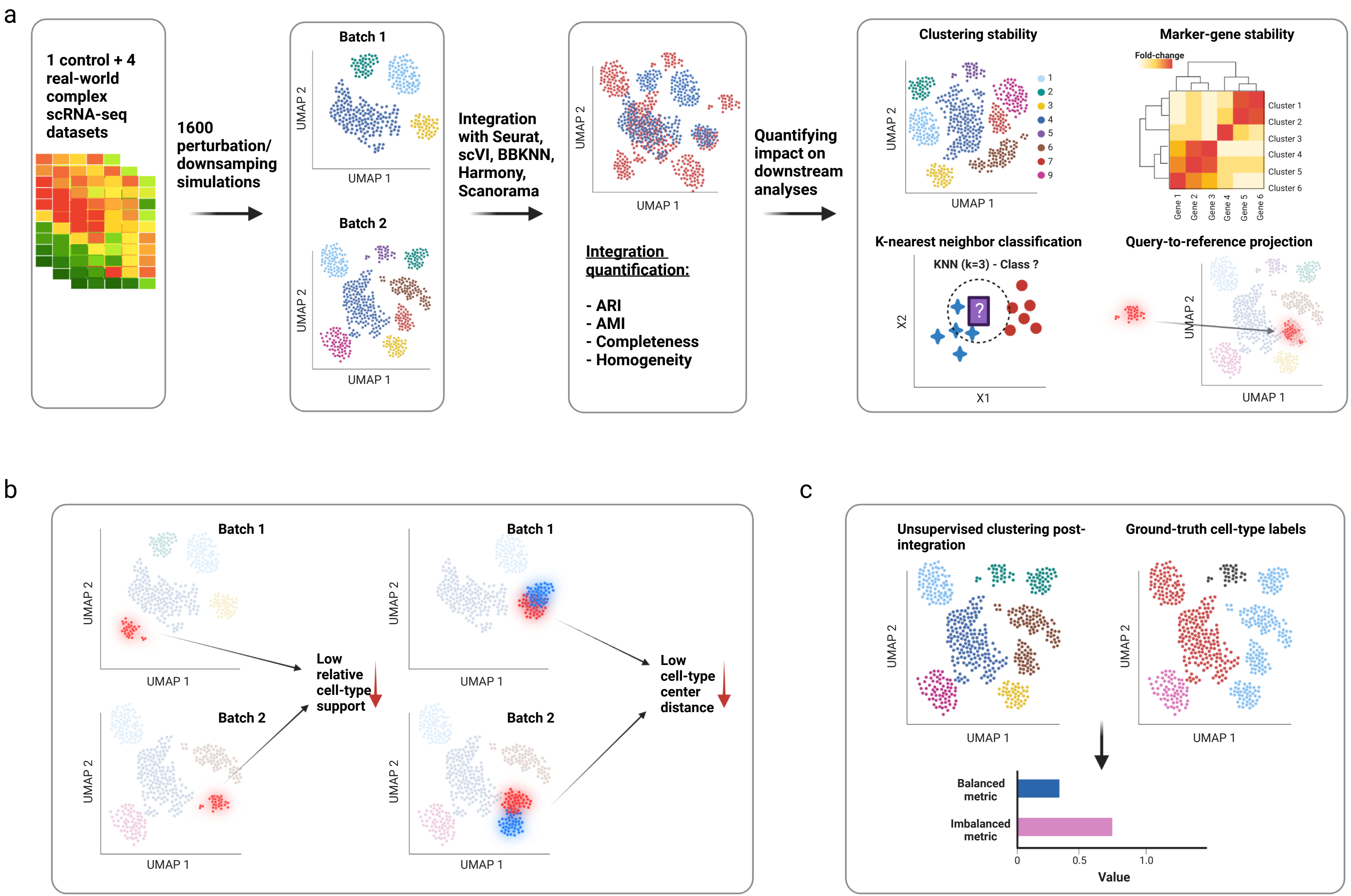

### Figure_2.pdf

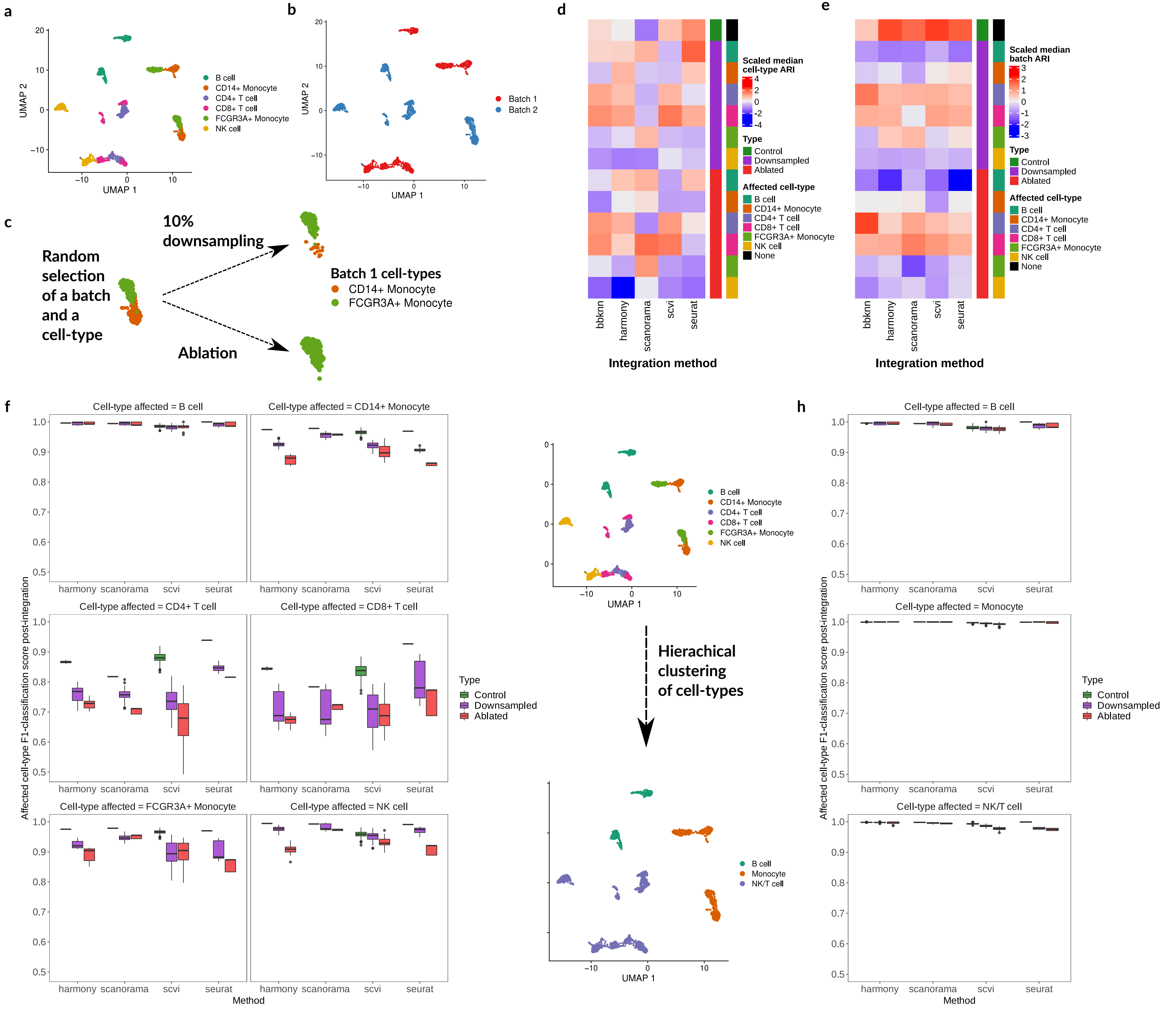

### Figure_3.pdf

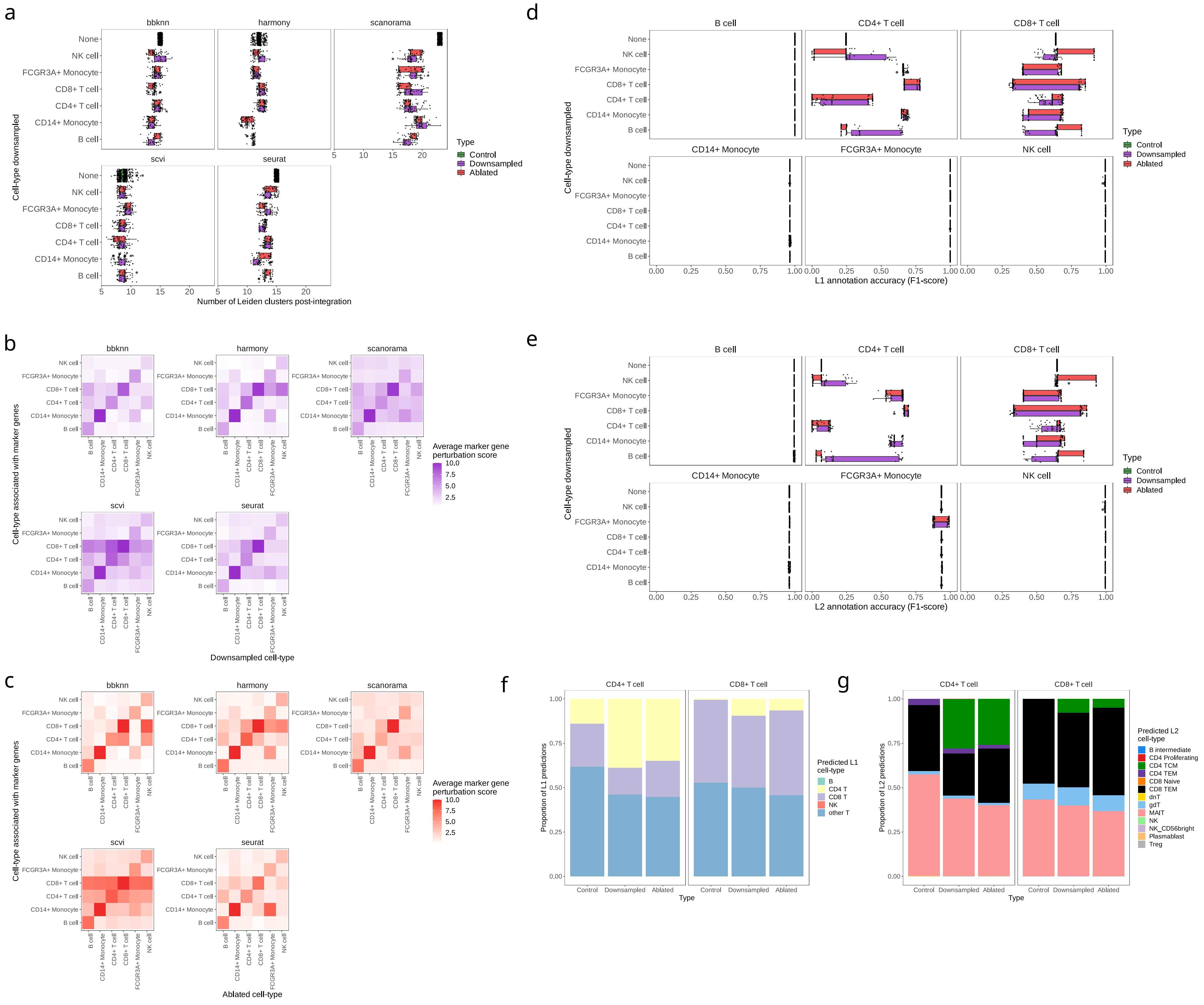

### Figure_4.pdf

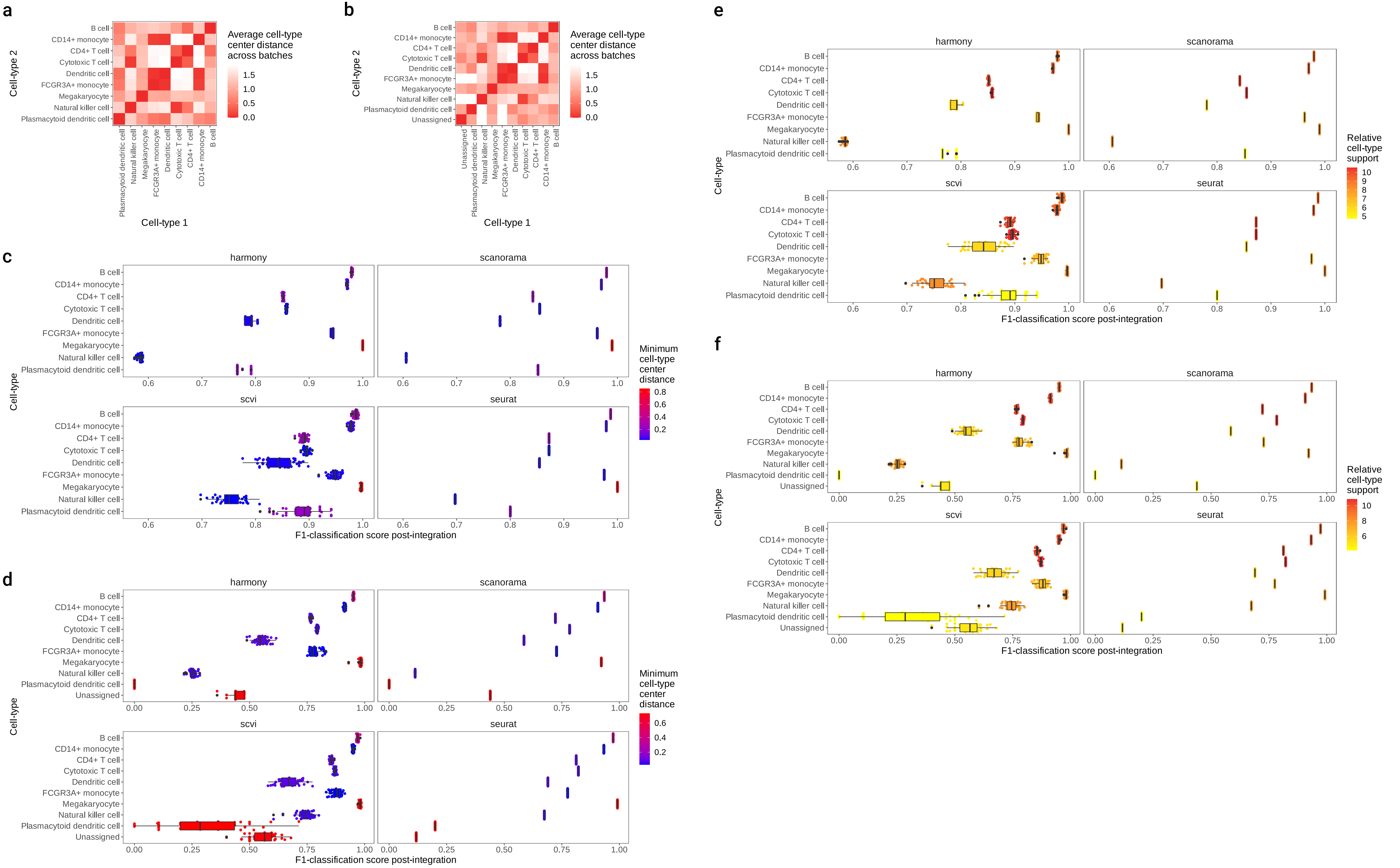

### Figure_5.pdf

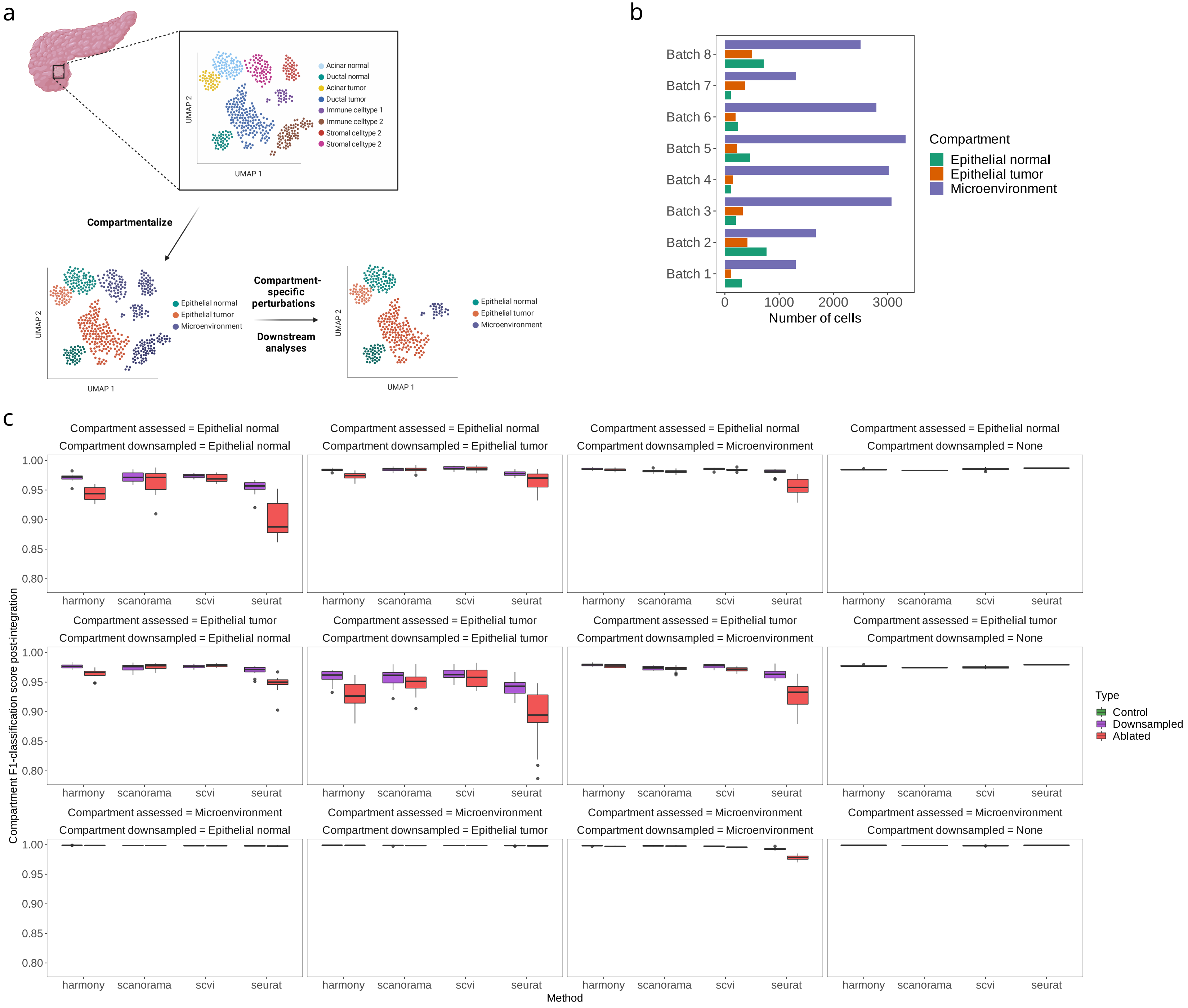

### Figure_6.pdf

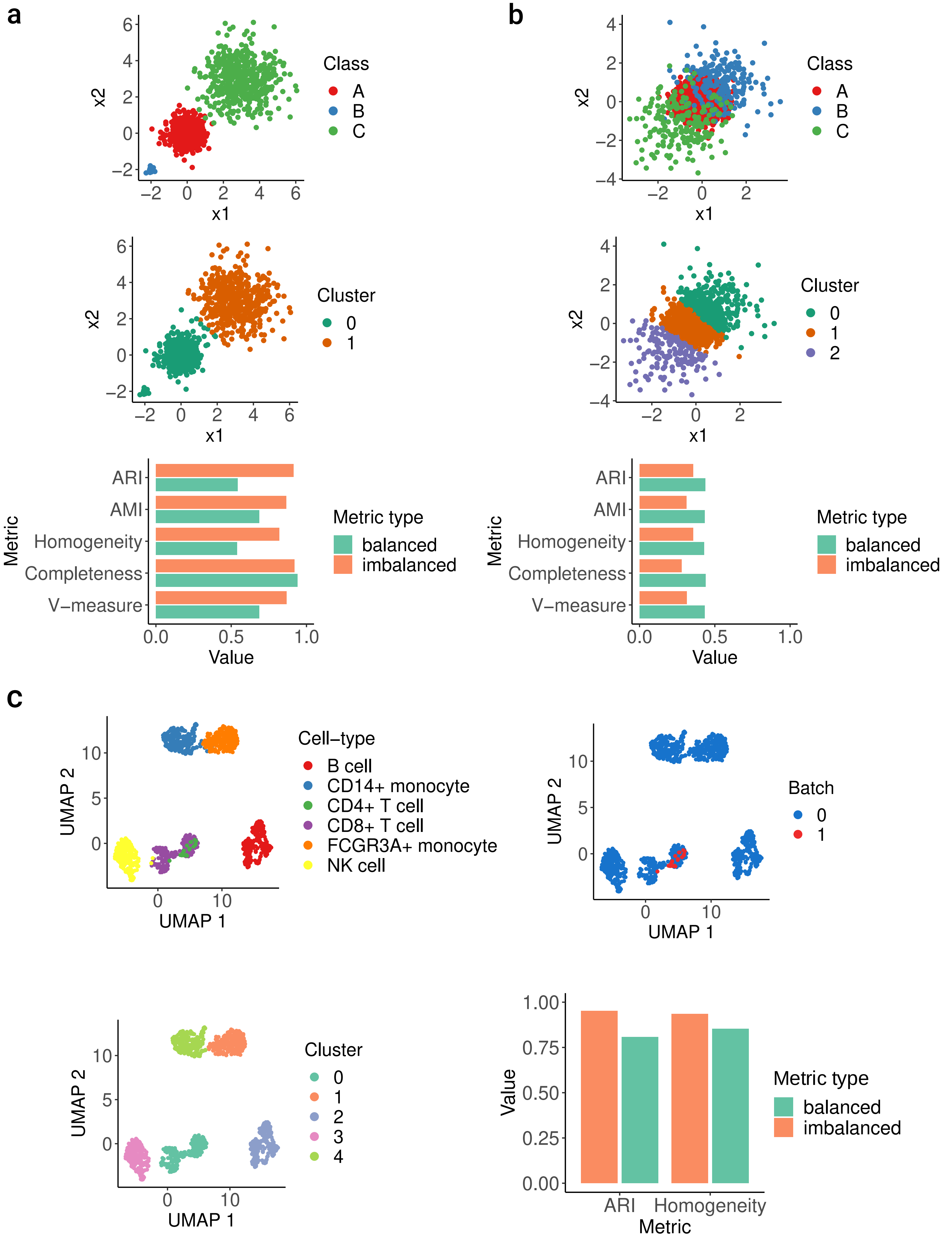

### Figure_7.pdf

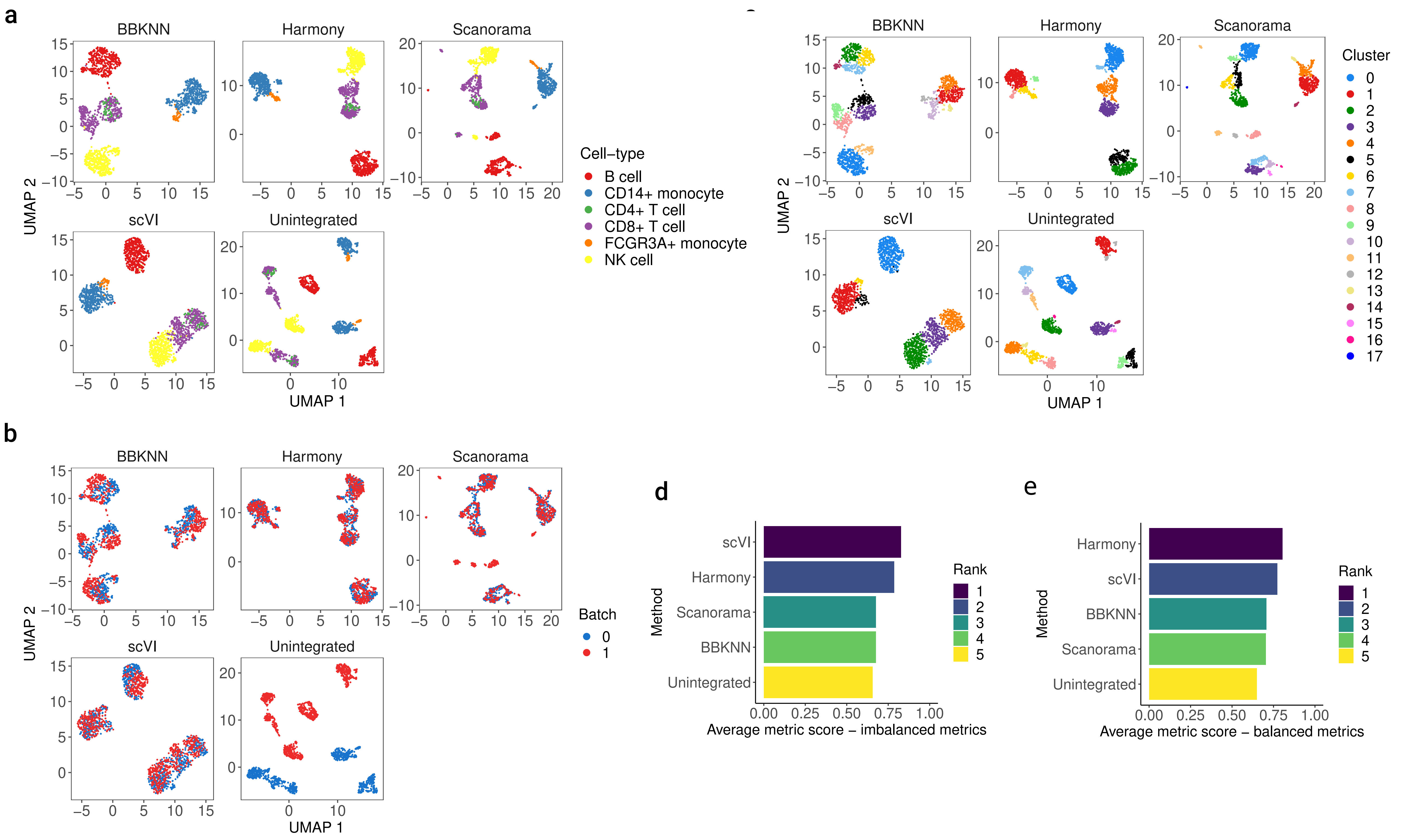

### Figure_8.pdf

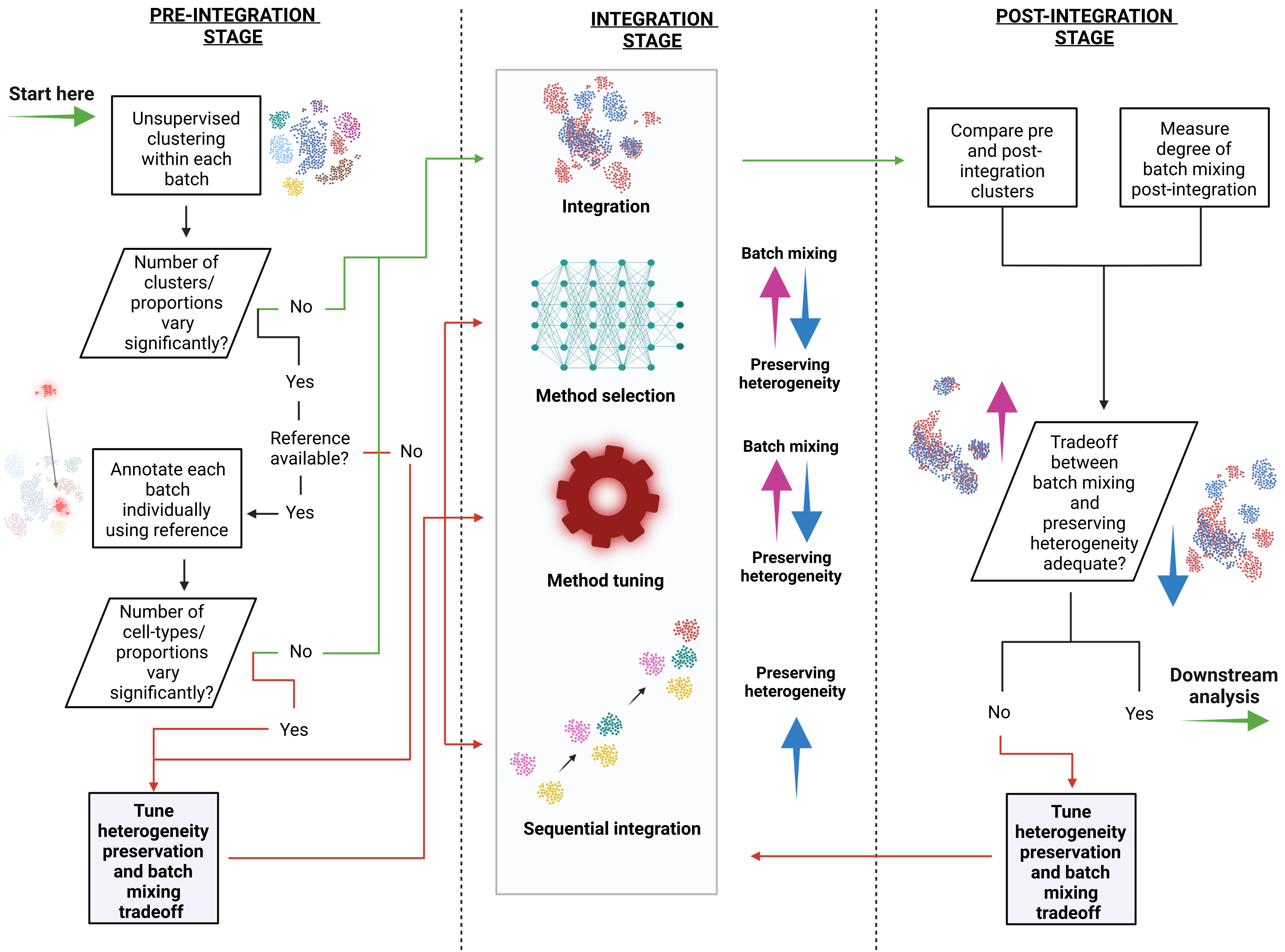

### Supplementary_Figure_1.pdf

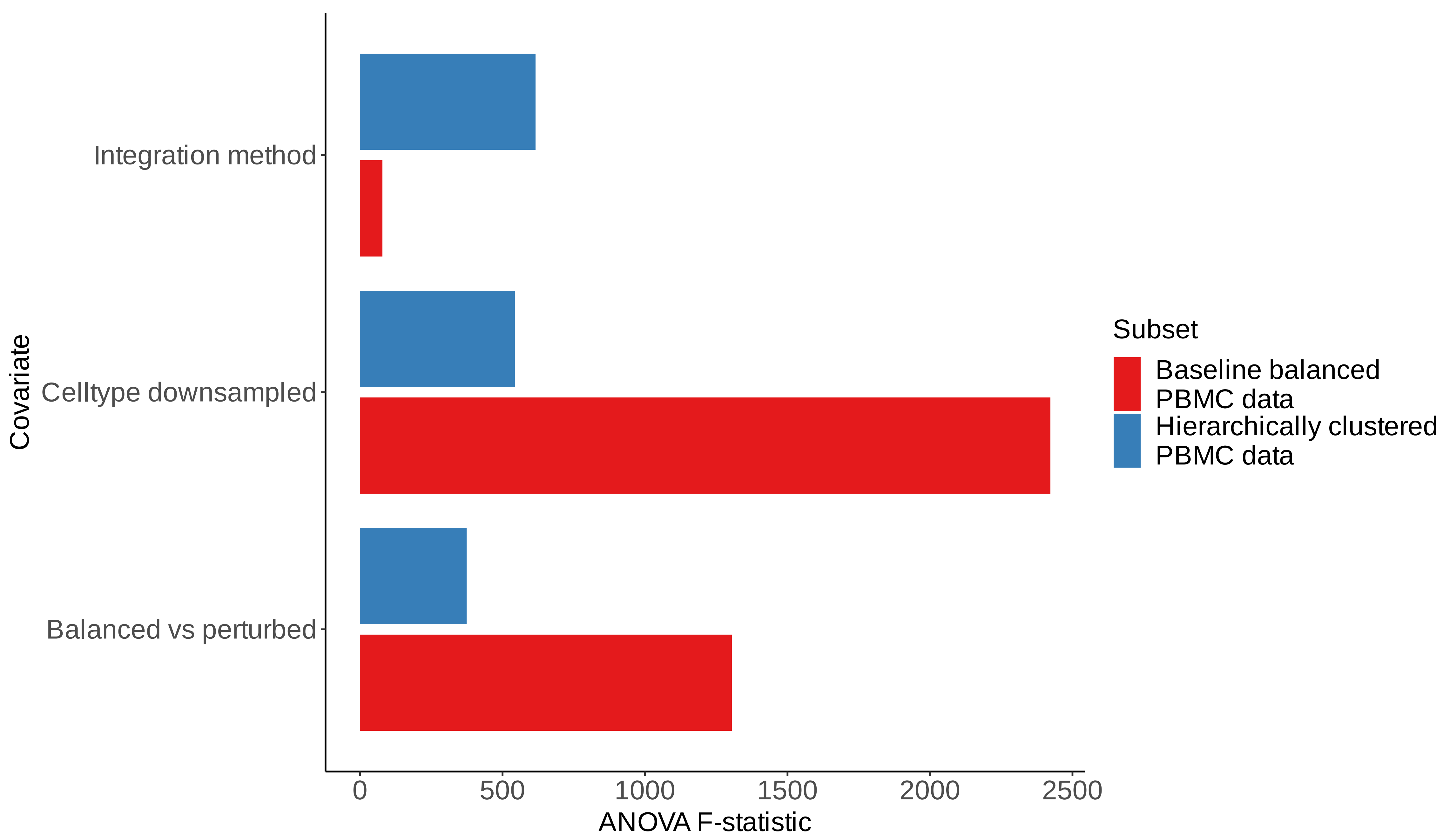

### Supplementary_Figure_2.pdf

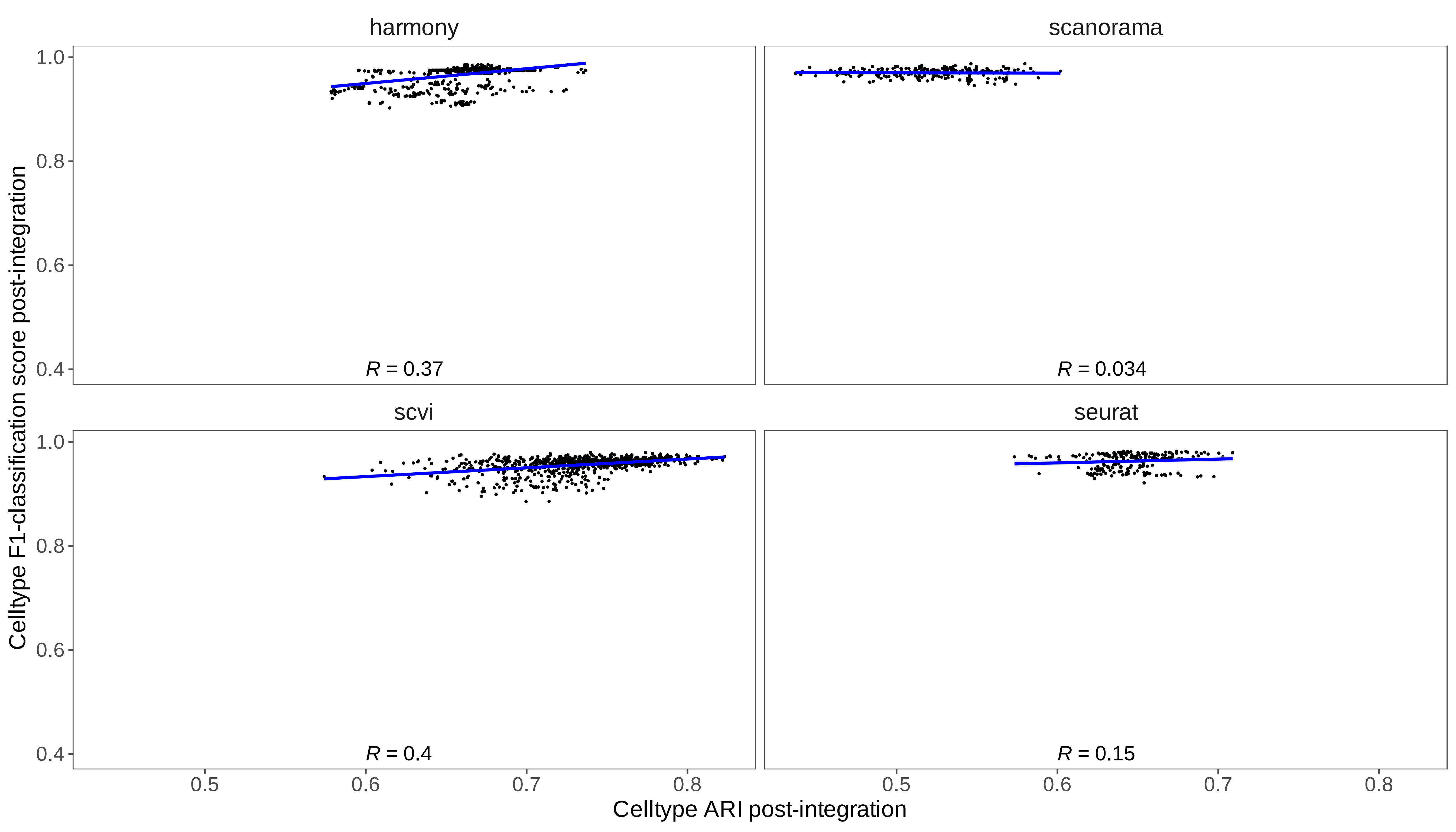

### Supplementary_Figure_3.pdf

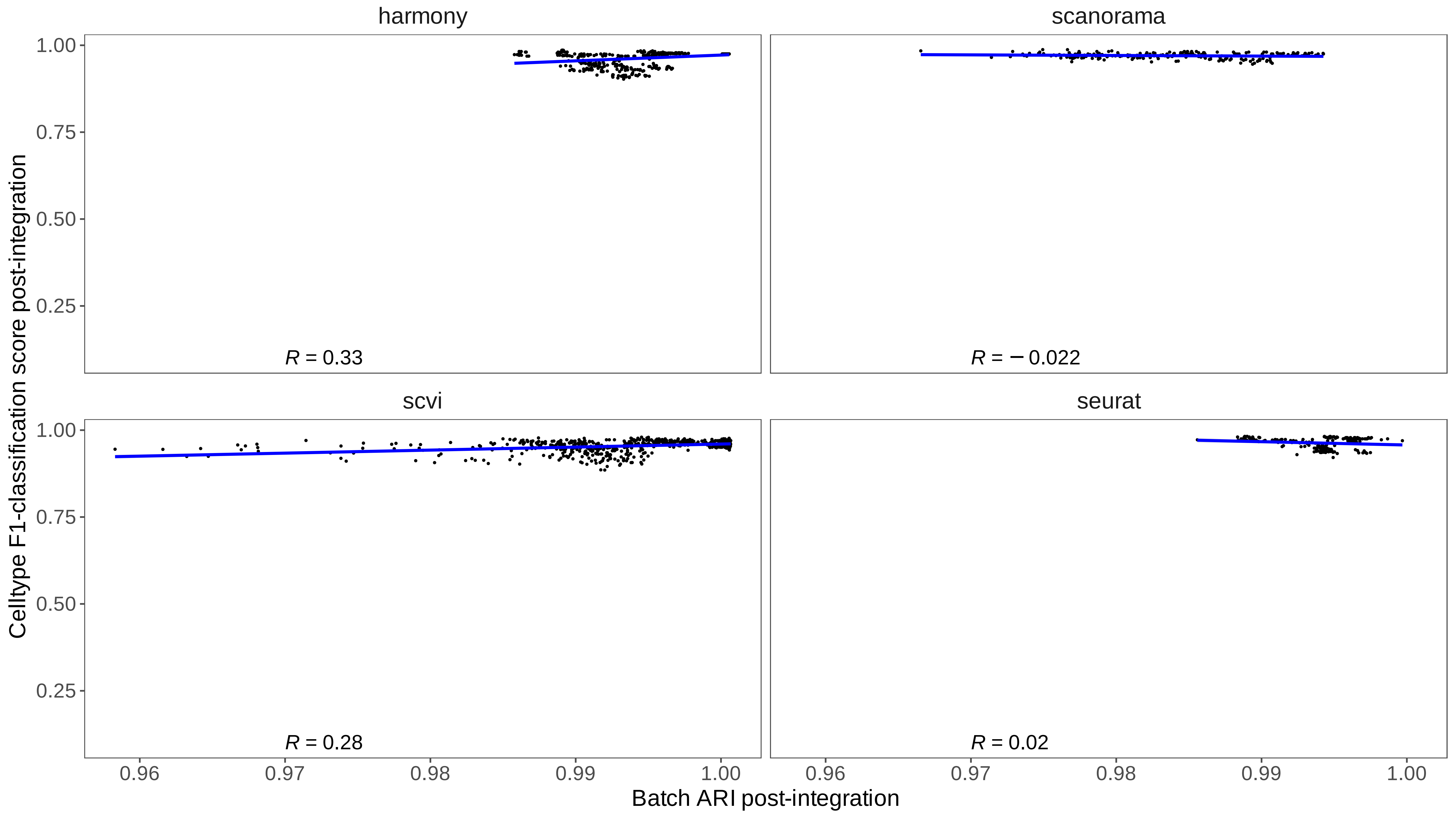

### Supplementary_Figure_4.pdf

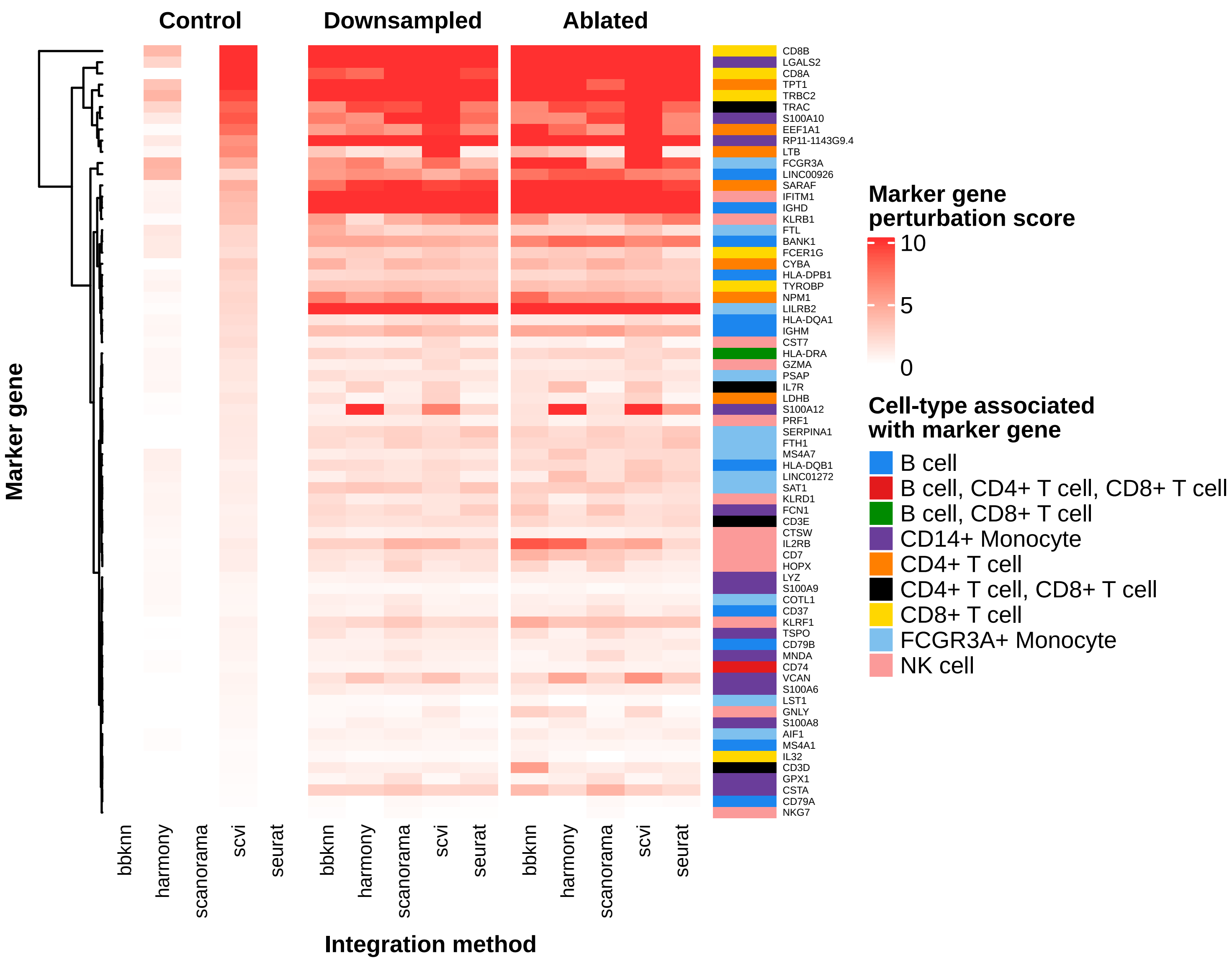

### Supplementary_Figure_5.pdf

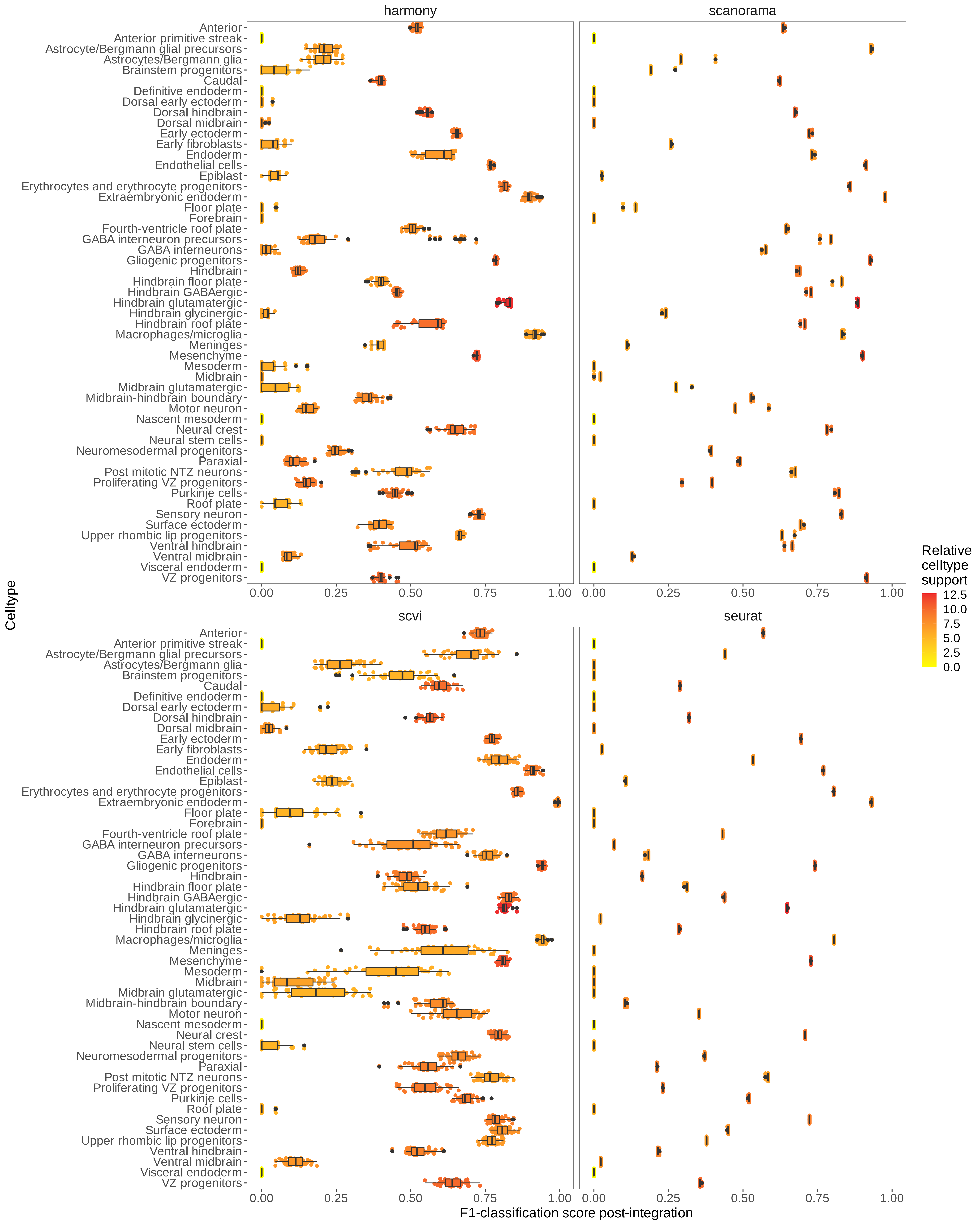

### Supplementary_Figure_6.pdf

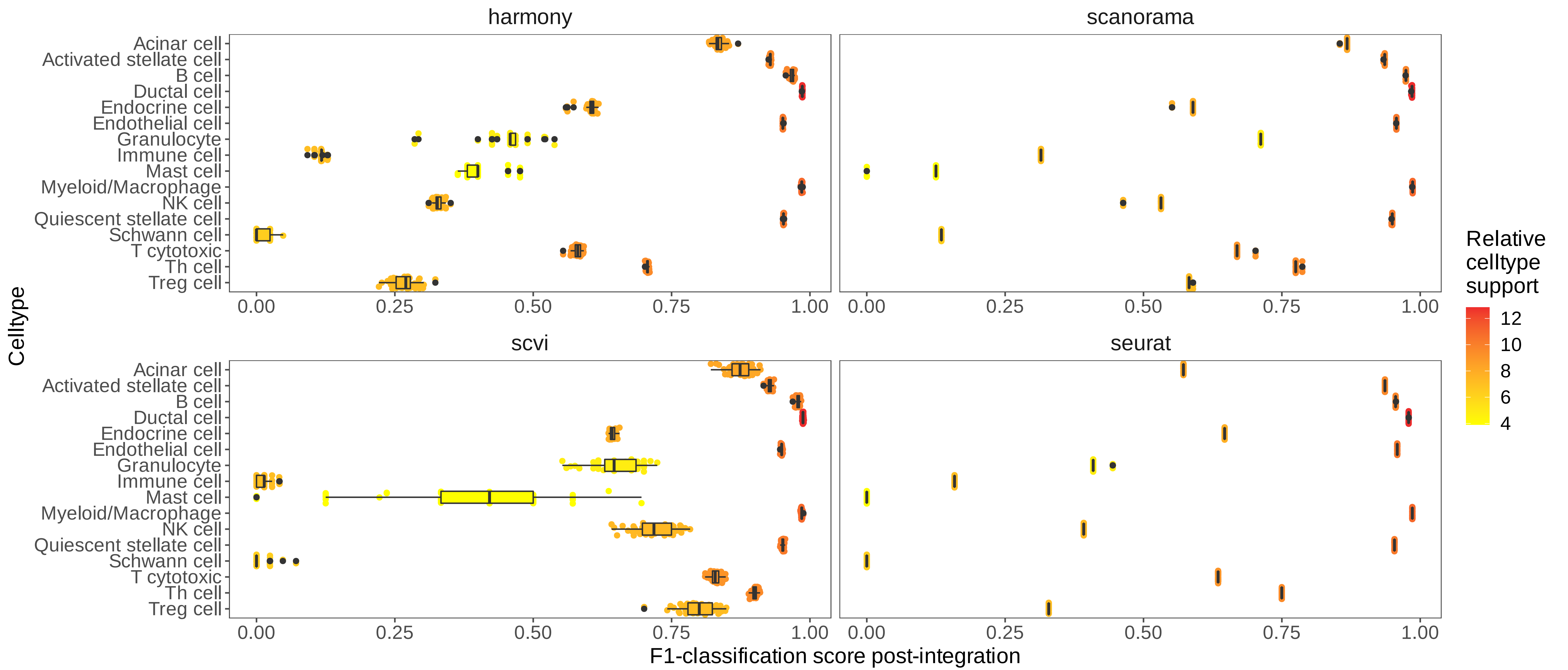

### Supplementary_Figure_7.pdf

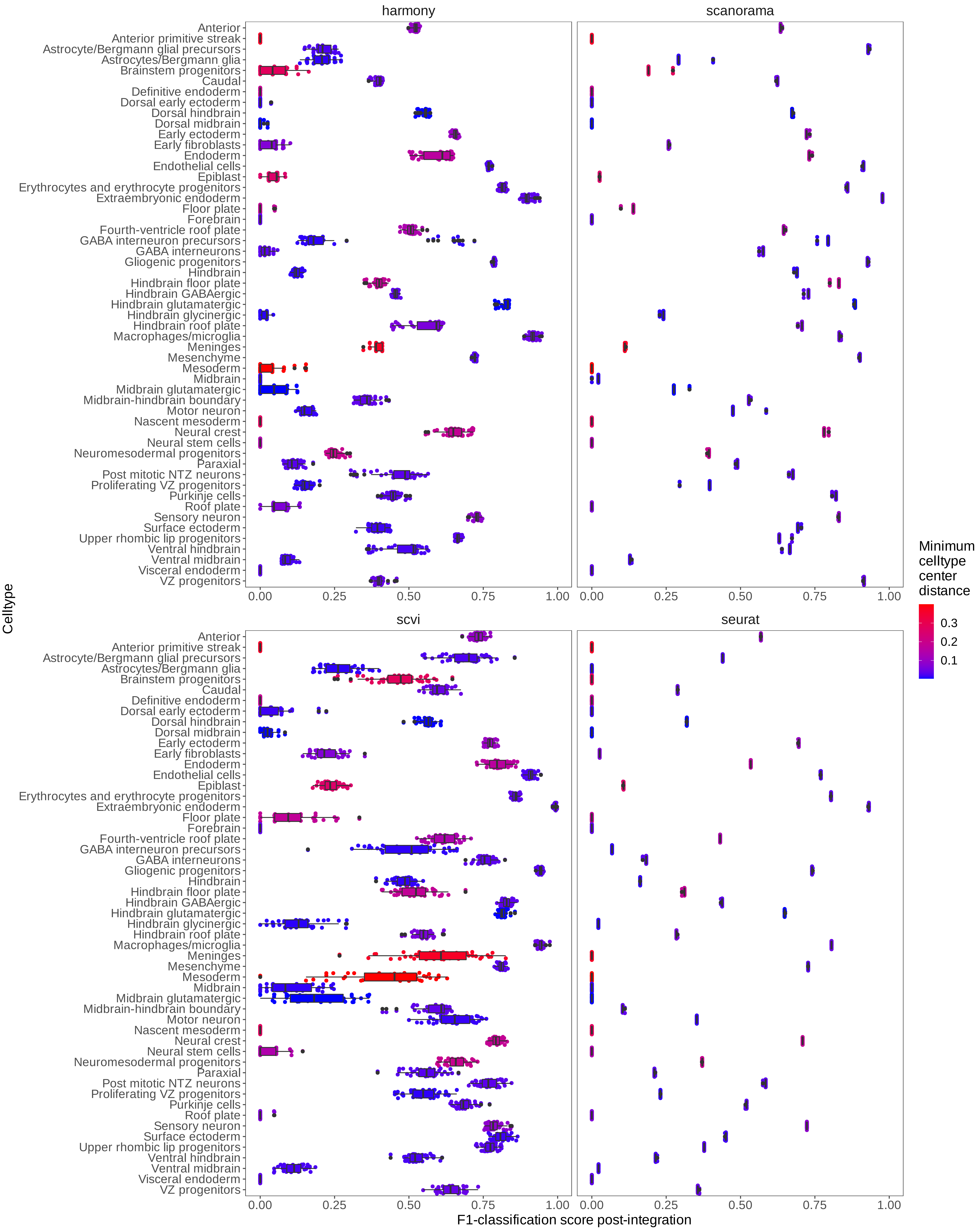

### Supplementary_Figure_8.pdf

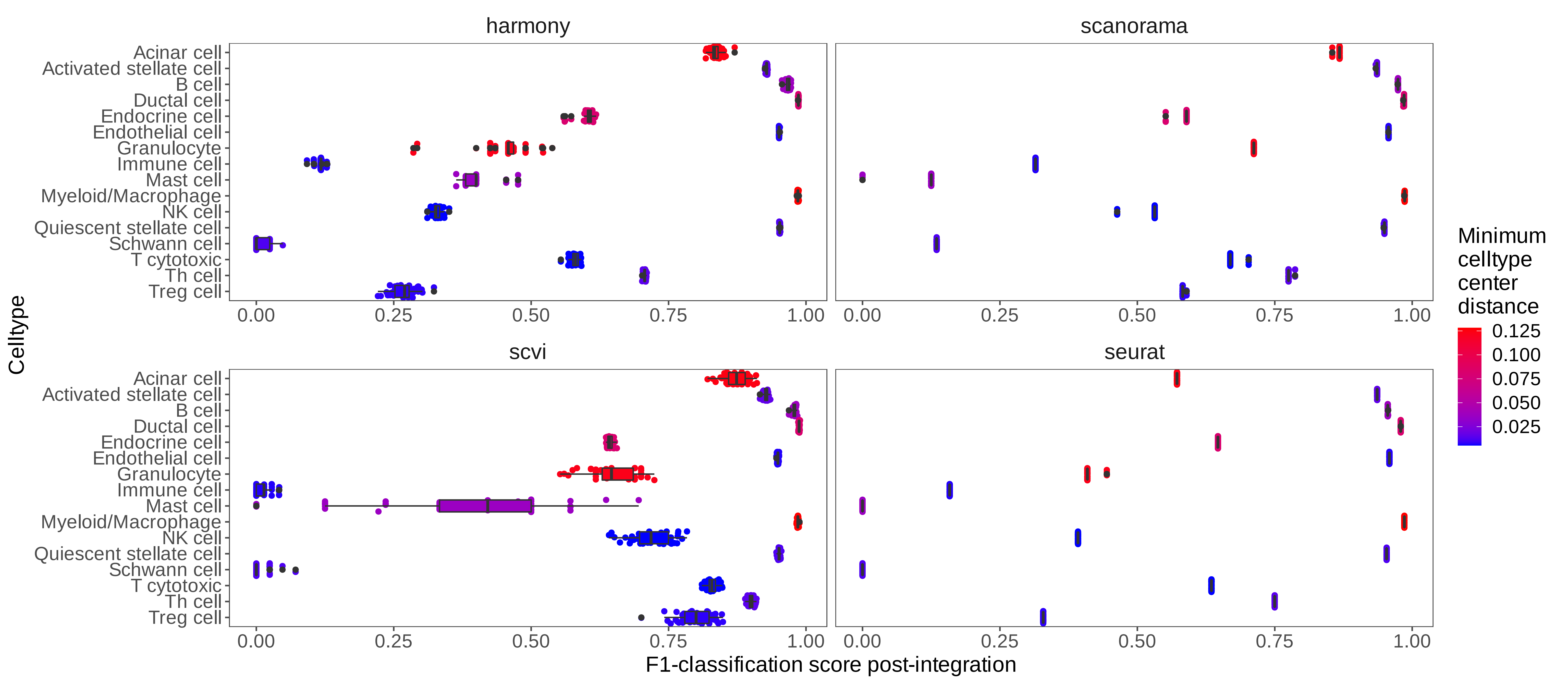

### Supplementary_Figure_9.pdf

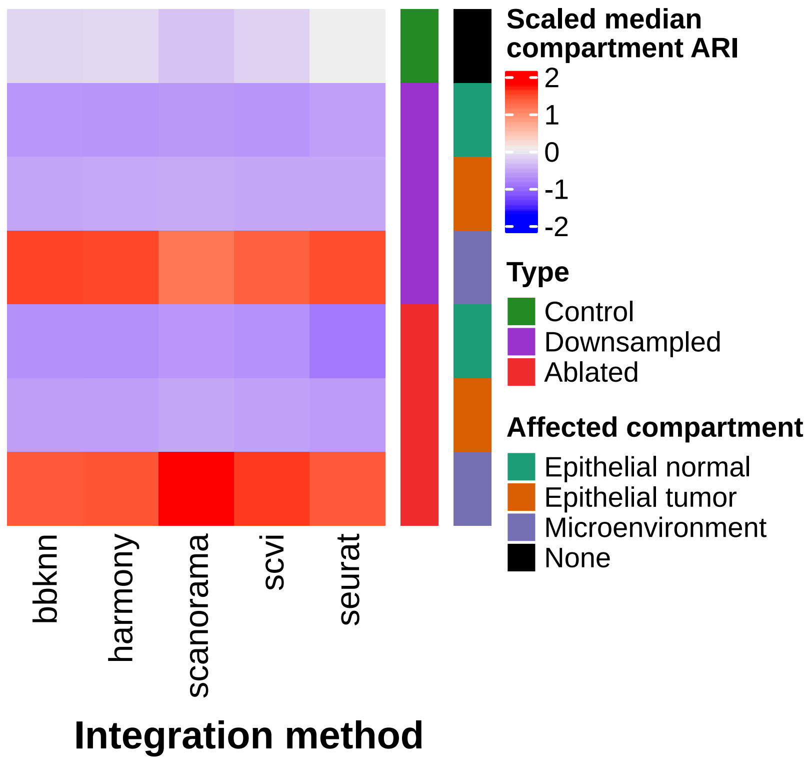

### Supplementary_Figure_10.pdf

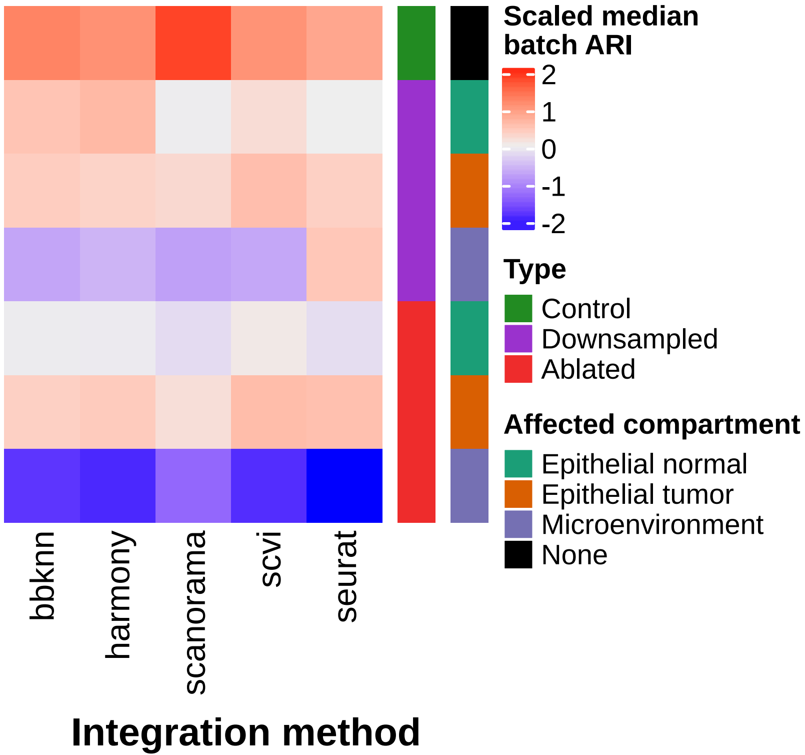

### Supplementary_Figure_11.pdf

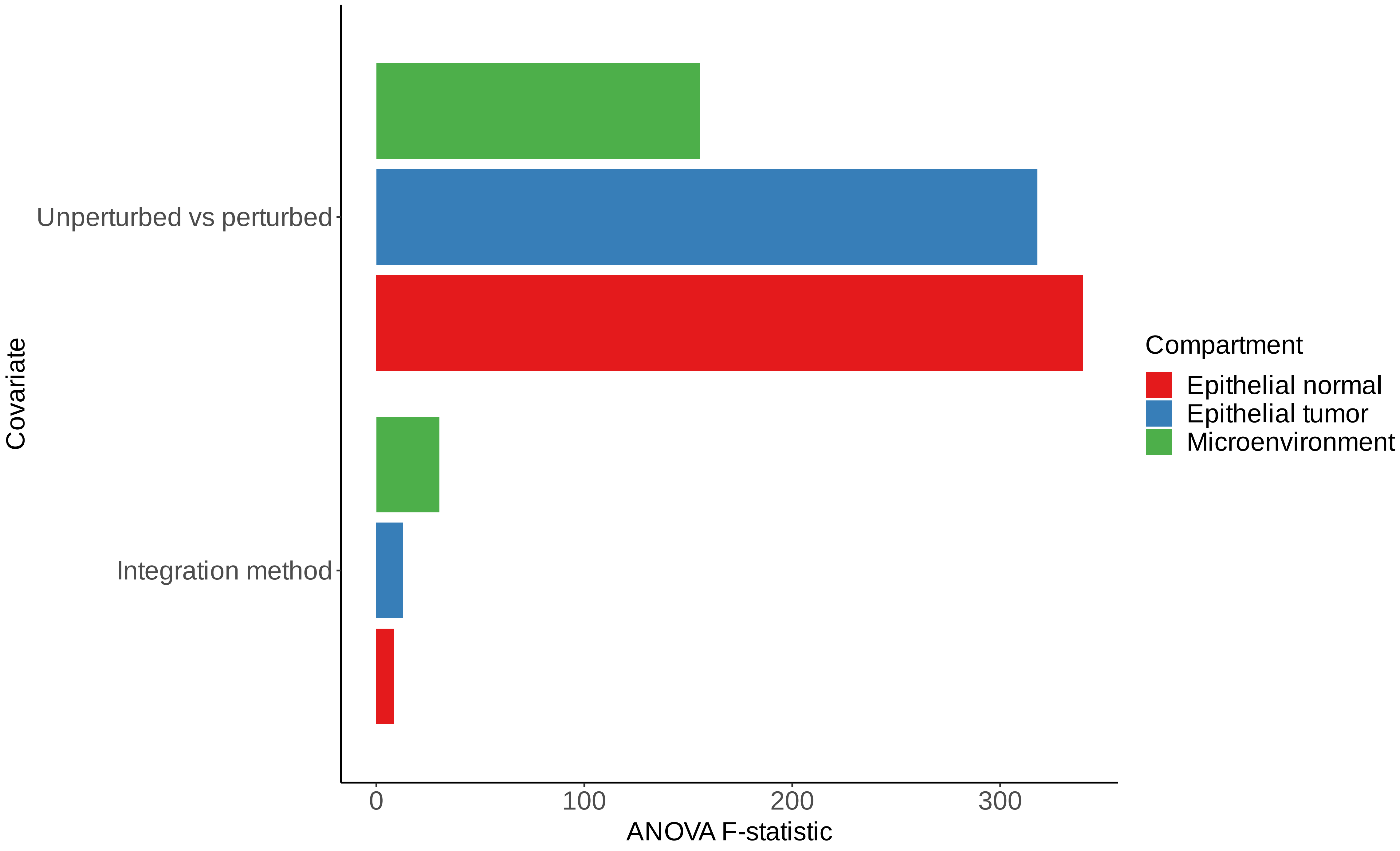

### Supplementary_Figure_12.pdf

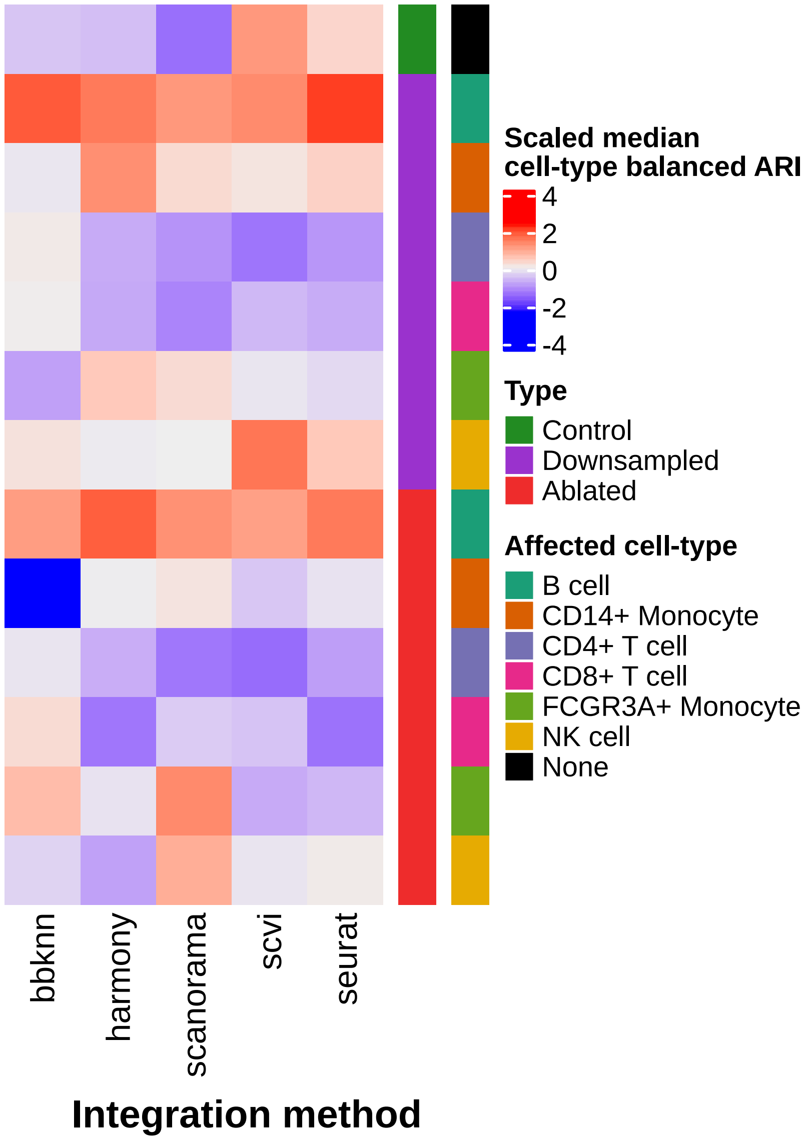

### Supplementary_Figure_13.pdf

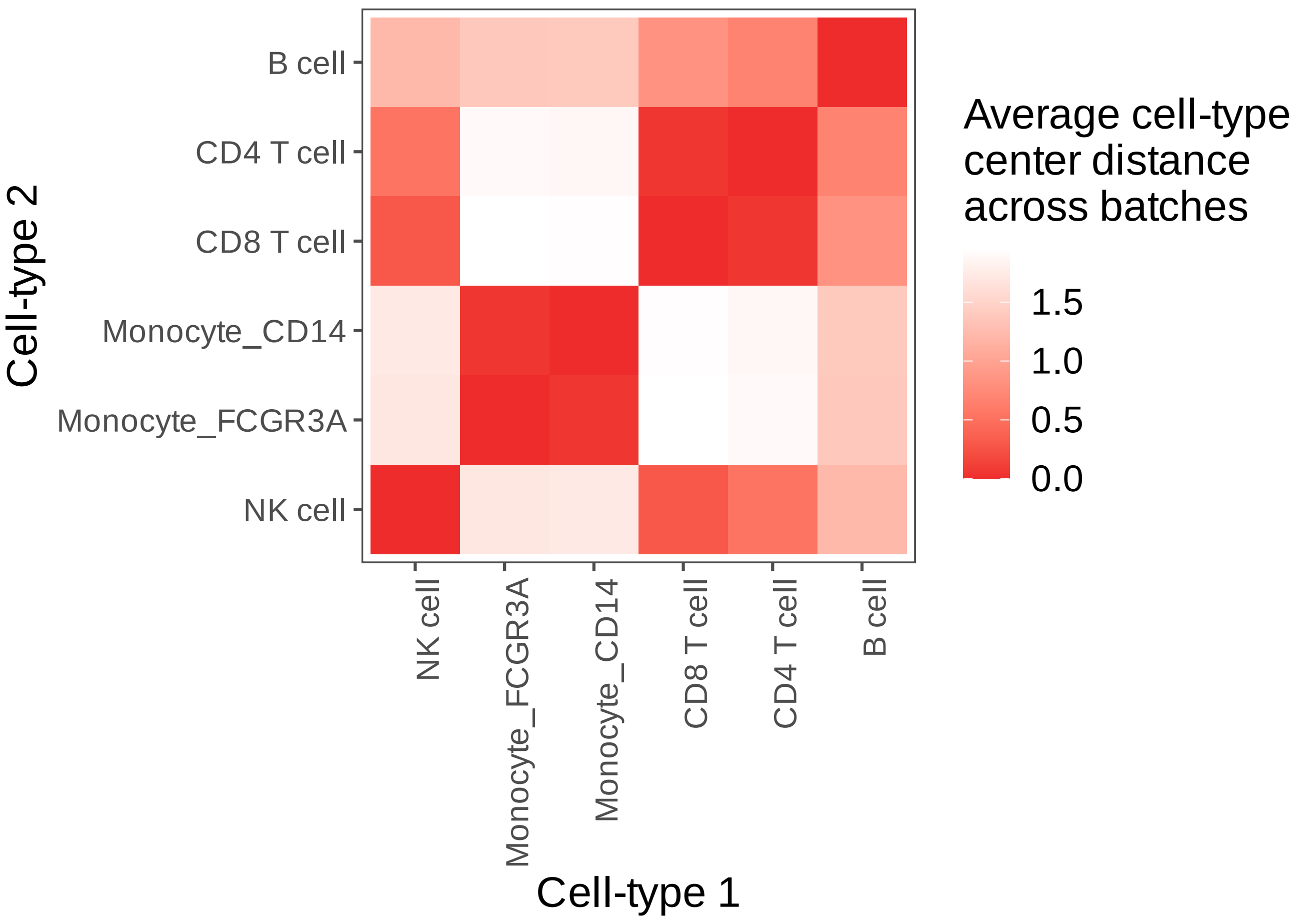

### Supplementary_Figure_14.pdf

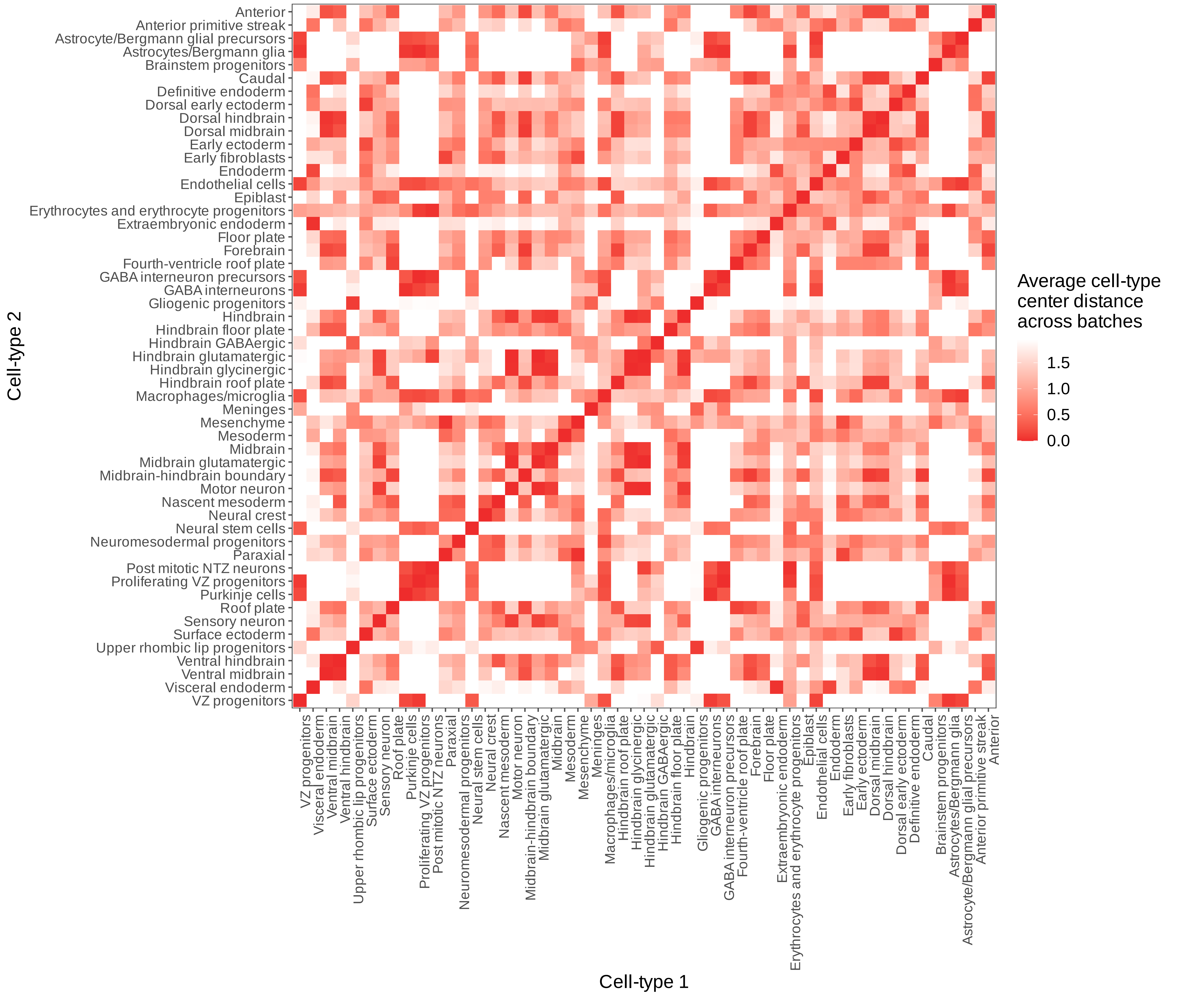

### Supplementary_Figure_15.pdf

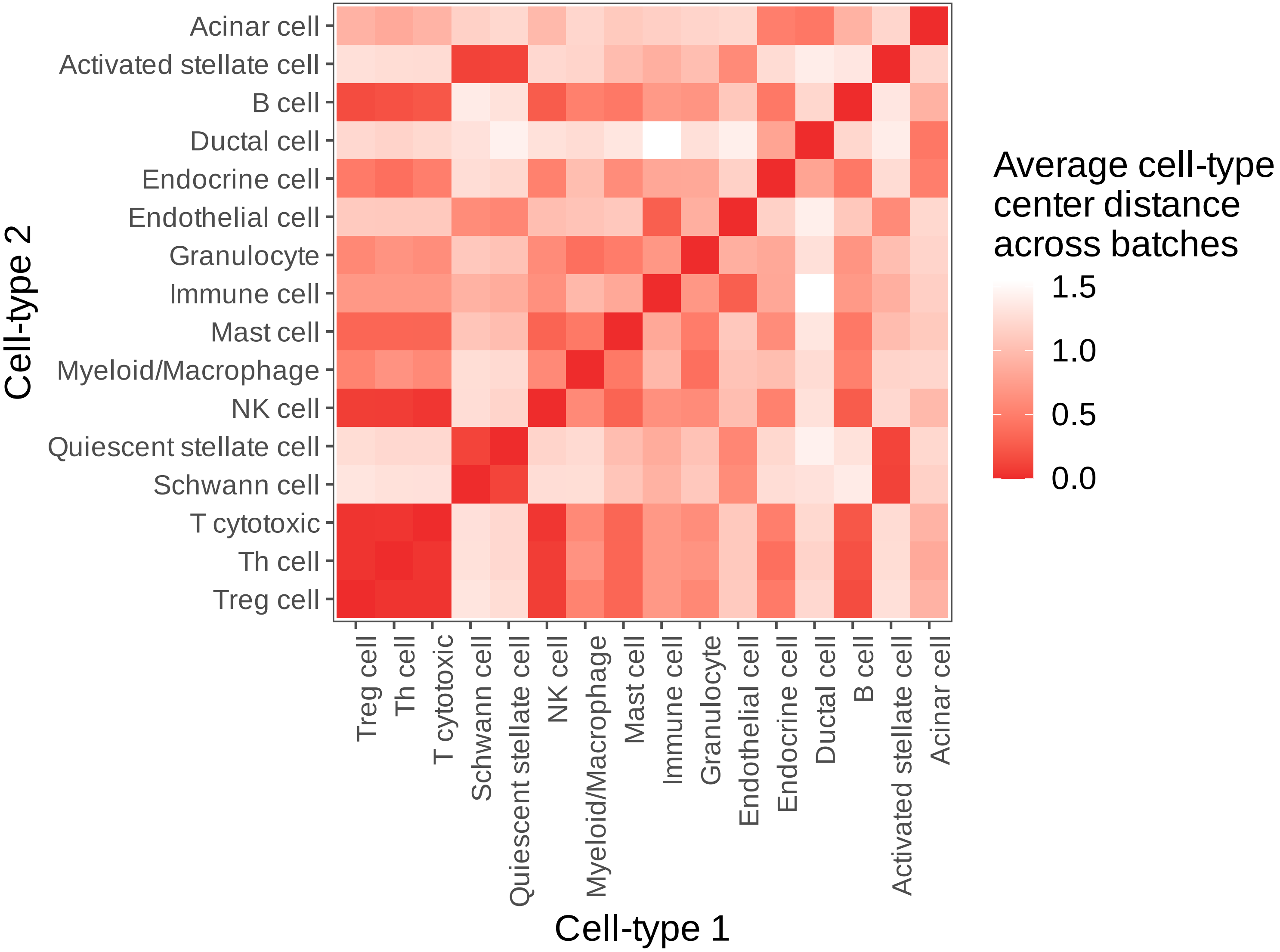

### Supplementary_Figure_16.pdf

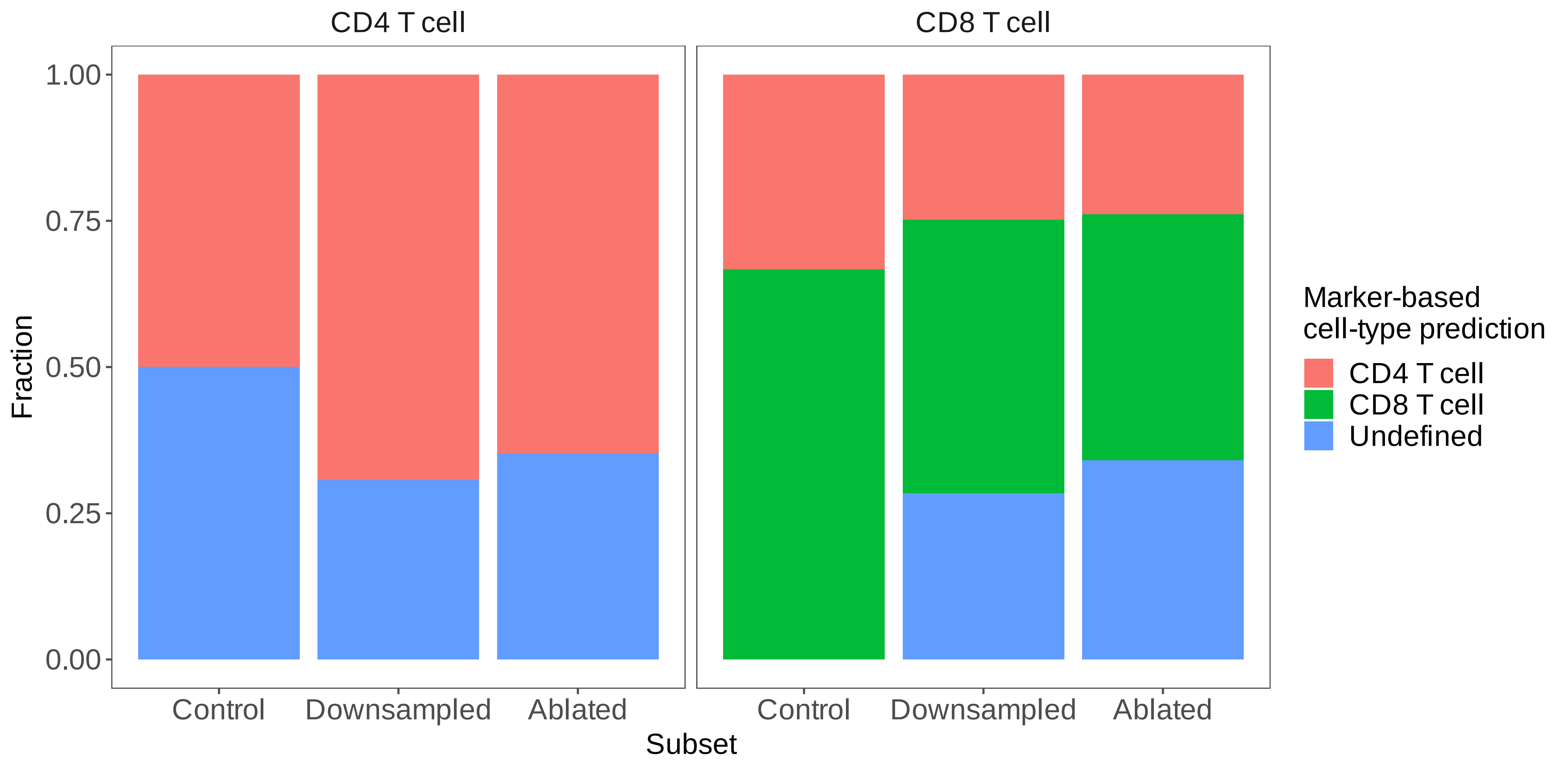
